# Demographics and Employment of Max-Planck Society’s Postdocs

**DOI:** 10.1101/2020.11.27.399733

**Authors:** M. Vallier, M. Mueller, P. Alcami, G. Bellucci, M. Grange, YX. Lu, S. Duponchel

## Abstract

The recently founded Max Planck PostdocNet brings together postdoctoral researchers (or postdocs) from the Max Planck Society (MPS), provides representation for the postdoctoral community across all Max Planck Institutes (MPI) and associated institutes, and advocates for their interests on their behalf.

At the 2019 founding meeting, MPS postdocs quickly raised their concerns about their employment situation and their associated social and working conditions. Subsequently, the PostdocNet conducted the first survey targeting exclusively postdoctoral researchers to gather information on their demographics, employment situations and social conditions.

This report presents the results of this survey, providing a thorough characterization of the postdoc demographics as well as the working conditions experienced by the postdoctoral community of the MPS. Remarkably, the survey analysis revealed a number of disparities in the access to employment type, wage level and social benefits. These results will guide future and present MPS postdoctoral researchers and their employers at the MPS to thrive for equality and fairness. Moreover, these results should be of help to the MPS to establish, improve and maintain optimal working conditions for MPS postdocs.

## I. Introduction

Founded in April 2019, the Max Planck PostdocNet brings together postdoctoral researchers (hereafter referred to as postdocs) from the Max Planck Society (MPS), provides representation for the postdoctoral community across all Max Planck Institutes (MPI) and associated institutes, and advocates for their interests on their behalf. The PostdocNet’s self-defined goal is to improve working conditions and scientific development of postdocs, and to enhance their career perspectives.

During the PostdocNet founding meeting, postdocs raised concerns about their employment and associated social and working conditions. Despite the consensus on the urgent need to tackle these issues, empirical data were needed to understand fully postdocs’ social and working conditions at the MPS, and foster constructive discussions for their improvement. Therefore, a survey group was formed within the PostdocNet to conduct the first comprehensive postdoc-led survey presented here. This survey aims to uncover the various social and employment difficulties faced by the postdoctoral community and to assess the current social landscape of postdocs within the MPS.

### Design of experiment

To obtain a meaningful picture of the postdocs and their work conditions within the MPS, a set of 32 questions were designed and submitted to the postdoctoral community between the 31 July and 27 August 2019. This questionnaire focused on three main topics: (i) demographic characteristics; (ii) past and overall research experiences and (iii) current employment conditions. Postdocs were also asked to indicate the MPS section (i.e. BM, Biology and Medical section; CPT, Chemistry, Physics and Technology section; HUM, Human Sciences section) to which their institutes belong. The topic on current employment conditions was subdivided to specifically address either postdocs on a fellowship (either a third-party fellowship or a MPS scholarship) or postdocs on a work contract based on the Collective Wage Agreement for the Civil Service (TVöD). Finally, the survey included free text boxes for postdocs to leave comments on their personal experiences and/or concerns at the end of the survey.

### What is a postdoc?

The definition of “postdoc” was at length discussed during the founding meeting and had to be taken into consideration during the establishment and analysis of the survey data. Therefore, the definition given in the “Guidelines for the Postdoc Stage”, internal document provided by the MPS, was chosen: postdocs are defined as “scientists who have obtained their doctorates and […] the postdoc phase is time-limited”^①^.

## II. Results

### A. Representativeness

This survey provides statistics on demographic aspects and employment conditions of MPS postdocs. The survey data was analyzed with the aim of comparing responses across region of origin, gender, research section, and, where relevant, past research experience. The questions used for the survey can be found in the supplementary **File S1**.

In total, 623 postdocs answered the survey partly or completely. This corresponds to 22.8% of all MPS postdocs (total number: 2,727; data provided on the 30.09.2019 by the MPS headquarters). Among these, 546 (87.6%) answered all “key” demographics and experience questions (gender, age, origin, graduation, experience, MPS section, employment type and funding source); 258/508 (50.8%) of contract holders answered all “key” questions about contracts; 38/82 (46.3%) of stipend holders answered all “key” questions about stipends. As participants did not always answer all questions, sample sizes for each question (or a combination thereof) may change and can be found in the respective figures.

Postdocs from 76 of the 87 MPIs within the MPS answered the survey. This could also be broken down further into the 3 sections to which they belong within the MPS: 32 of 33 MPIs from the Biology and Medical (BM) section, 25 of 33 MPIs from the Chemistry, Physics and Technology (CPT) section and 19 of 21 MPIs from the Human Sciences (HUM) section. The majority of responses came from postdocs working in the BM section (57.0%), followed by the CPT section (26.7%) and HUM section (16.3%) (**Fig 1A-C**).

**Figure 1.**
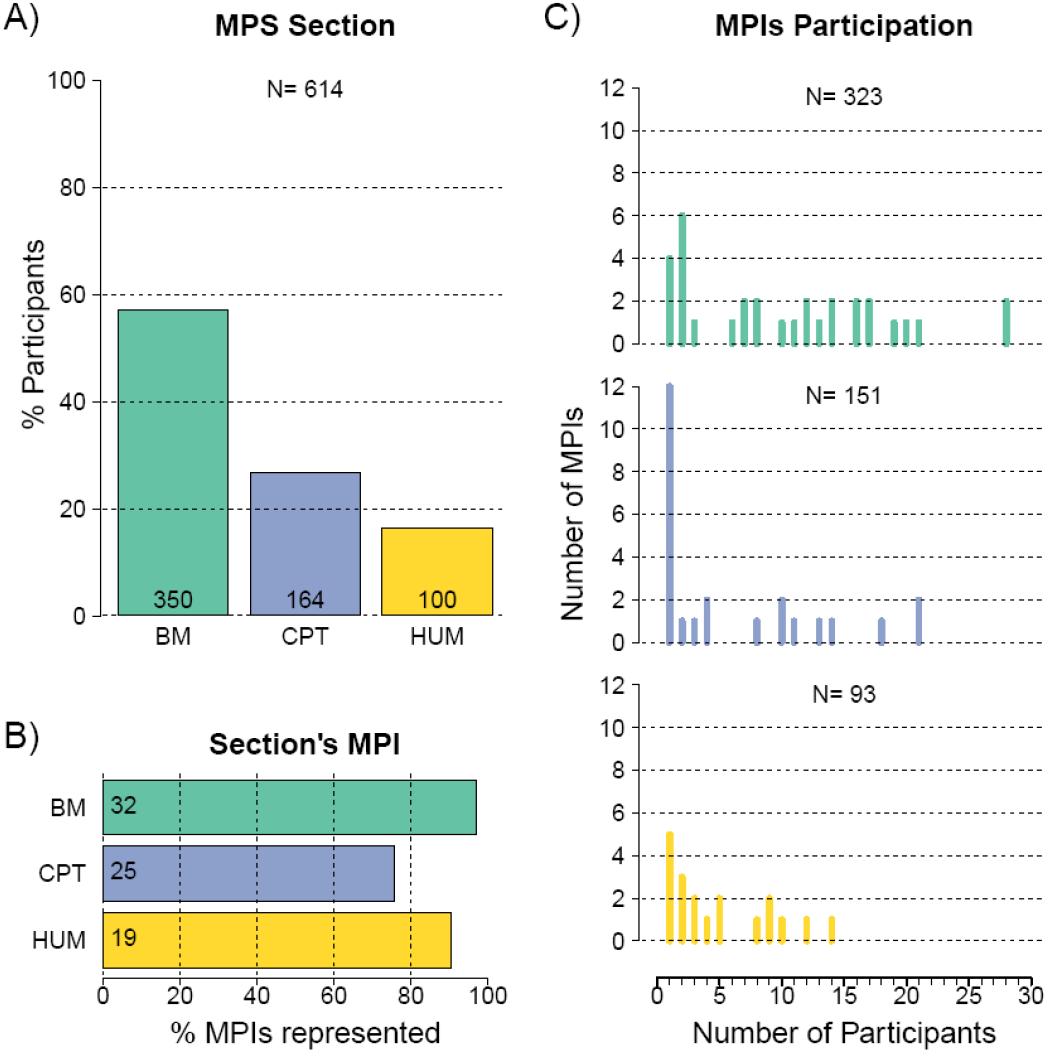
Representativeness of survey respondents across the MPS. Percentage of respondents belonging to each of the three MPS sections **(A)**. Percentage of MPIs represented per section **(B)**. Histogram of the number of participants per institute in each section **(C)**. Numbers in each bar graph refer to the number of answers given. BM, Biology and Medical section; CPT, Chemistry, Physics and Technology section; HUM, Human Sciences section; MPI, Max Planck Institute, MPS, Max Planck Society.

### B. Demographics

#### 1. Gender

Overall (**Fig 2A**), 58.1% of respondents identified as male, 41.4% as female and 0.5% subscribed to genders outside of these binary categories. Within the sections (**Fig 2B**), the HUM section had 52.6% male to 47.4% female, the CPT section had 71.7% male to 28.3% female and the BM section had 54.5% male to 45.5% female.

**Figure 2.**
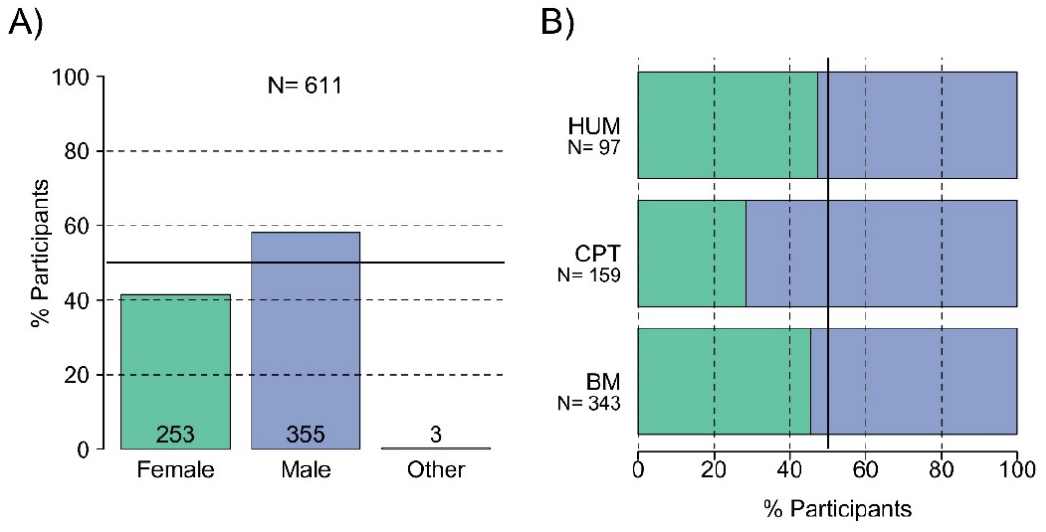
Gender* distribution. Gender distributions are given for the MPS overall **(A)** and for each MPS section **(B)**. Numbers in each bar graph refer to the number of answers given. The thick black line corresponds to 50%. BM, Biology and Medical section; CPT, Chemistry, Physics and Technology section; HUM, Human Sciences section. \**Although respondents were asked to answer, “male/female/other” in response to the question of their gender, the PostdocNet would like to state that it supports non-binary gender identity*.

A statistical analysis of the overall female and male distribution using chi-square test and Cramer’s V estimation revealed a significant overall male-bias of 16.8% (p<0.01) (**Tab 1**). When analyzed on a per-section basis, no significant male bias was found within the data from the BM and HUM sections (p>0.05), however the data from the CPT section revealed a significant male-bias of 43% (p=0.0001).

**Table 1.** Summary of the female to male distribution in the surveyed population. BM, Biology and Medical section; CPT, Chemistry, Physics and Technology section; HUM, Human Sciences section.

|  | Female | Male | Total | Male-bias | Chi-square to 50% | Cramer's V |
| --- | --- | --- | --- | --- | --- | --- |
| **Overall** | 253 | 355 | 608 | 16.8% | 0.0040 | 8.4% |
| **HUM** | 46 | 51 | 97 | 5.2% | 0.8294 | 2.6% |
| **CPT** | 45 | 114 | 159 | 43.4% | 0.0001 | 22.2% |
| **BM** | 156 | 187 | 343 | 9.0% | 0.2677 | 4.5% |

#### 2. Age

The significant majority of postdocs were between 30 and 40 years old (74.2%), with the next greatest proportion of postdocs being between 20 and 30 years old (16.3%).

With 9.5%, those postdocs older than 40 years old were the smallest group (**Fig 3A**). Similar distributions of age were observed in each MPS section (**Fig 3B**).

**Figure 3.**
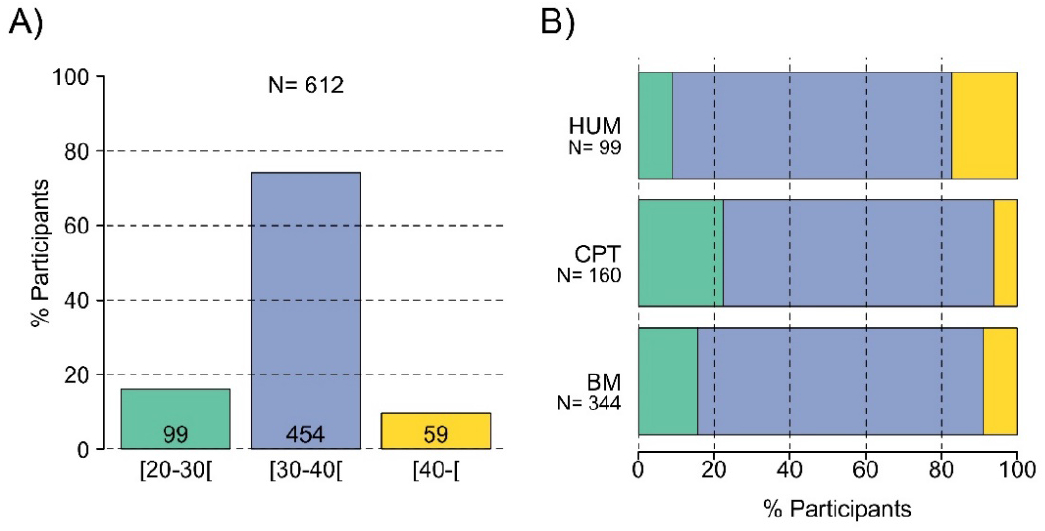
Age group distribution. Age group distributions are given for the MPS overall **(A)** and for each MPS section **(B)**. Numbers in each bar graph refer to the number of answers given. BM, Biology and Medical section; CPT, Chemistry, Physics and Technology section; HUM, Human Sciences section.

When considering the distribution of males and females depending on their age (**Fig 4, Tab S1**), there was no significant difference in the younger age group, between 20 and 30 years old. The gender distribution significantly differed from the 50% expectation in the middle age group, from 30 to 40 years old and was slightly lower for females in the older age group, from 40 years old on.

**Figure 4.**
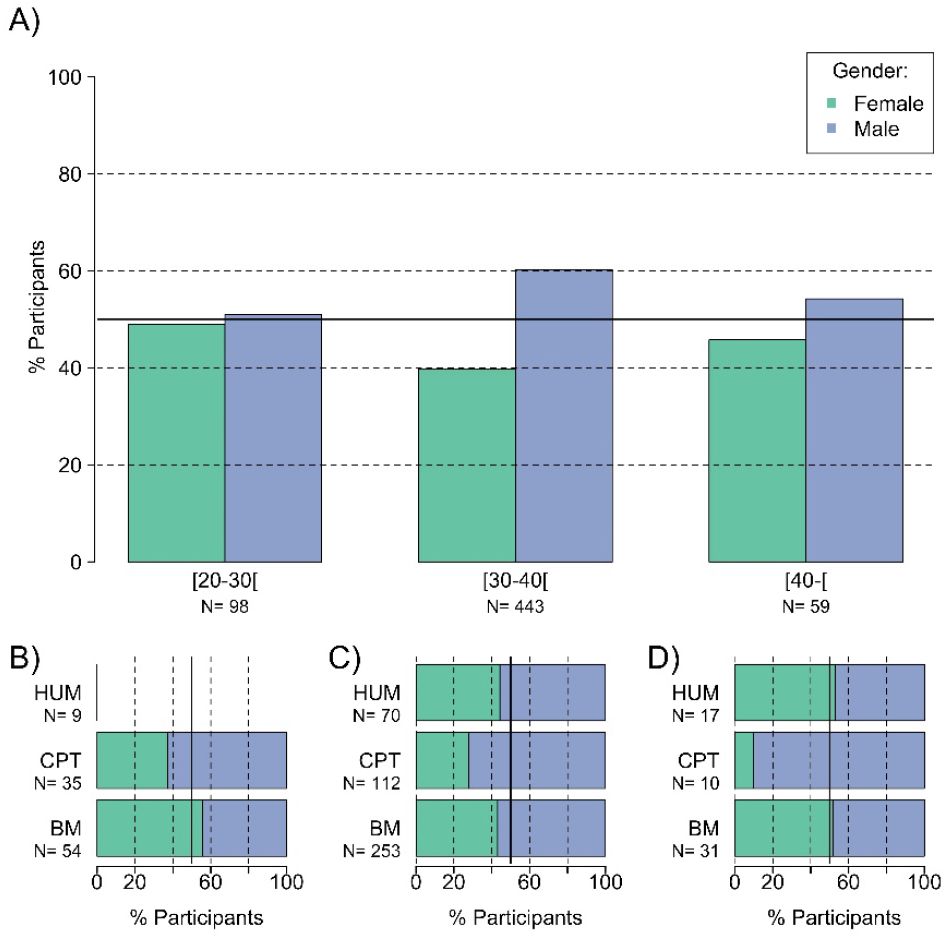
Gender and age group. Comparison of gender distribution based on age group in the MPS overall **(A)**. In each section, the gender distribution for each age groups is separated: younger age group [20-30[**(B)**, middle age group [30-40[ **(C)**, and older age group [40-[ **(D)**. Note, statistics with less than 10 data points are not shown on graphs. The black solid line corresponds to 50%. BM, Biology and Medical section; CPT, Chemistry, Physics and Technology section; HUM, Human Sciences section.

This distribution varied between MPS sections (**Fig 4B-D**). Within the BM and HUM sections, there was no significant difference between gender distributions across the three age groups, with a trend of more females in younger (**Fig 4B**) and older (**Fig 4D**) age groups. However, in the CPT section, the distribution deviated significantly from 50% towards a male bias across all age groups.

#### 3. Region of origin

The postdoctoral community was made up of employees coming from all regions of the world (**Fig 5A**). The least represented area was Africa with only 0.6% (4/607) of respondents coming from this continent, followed by Oceania with 2.5% (15/607). The majority of postdocs (62.9%, 382/607) originated from Europe, which includes Germany, other European Union countries (28 countries including UK^②^, referred to as EU) and European non-EU countries (Europe). Asia was the second most represented continent with 19% (115/607), while 15% (91/607) originated from the Americas (North and South America).

These observations remained the same when looking at the distribution of origins within MPS sections, with the notable exception for postdocs coming from Asia (**Fig 5B**, in pink). Their proportion was the biggest in the CPT section (25%, 40/160), followed by the BM (19.6 %, 67/341) and the HUM (7.1 %, 7/98) sections.

**Figure 5.**
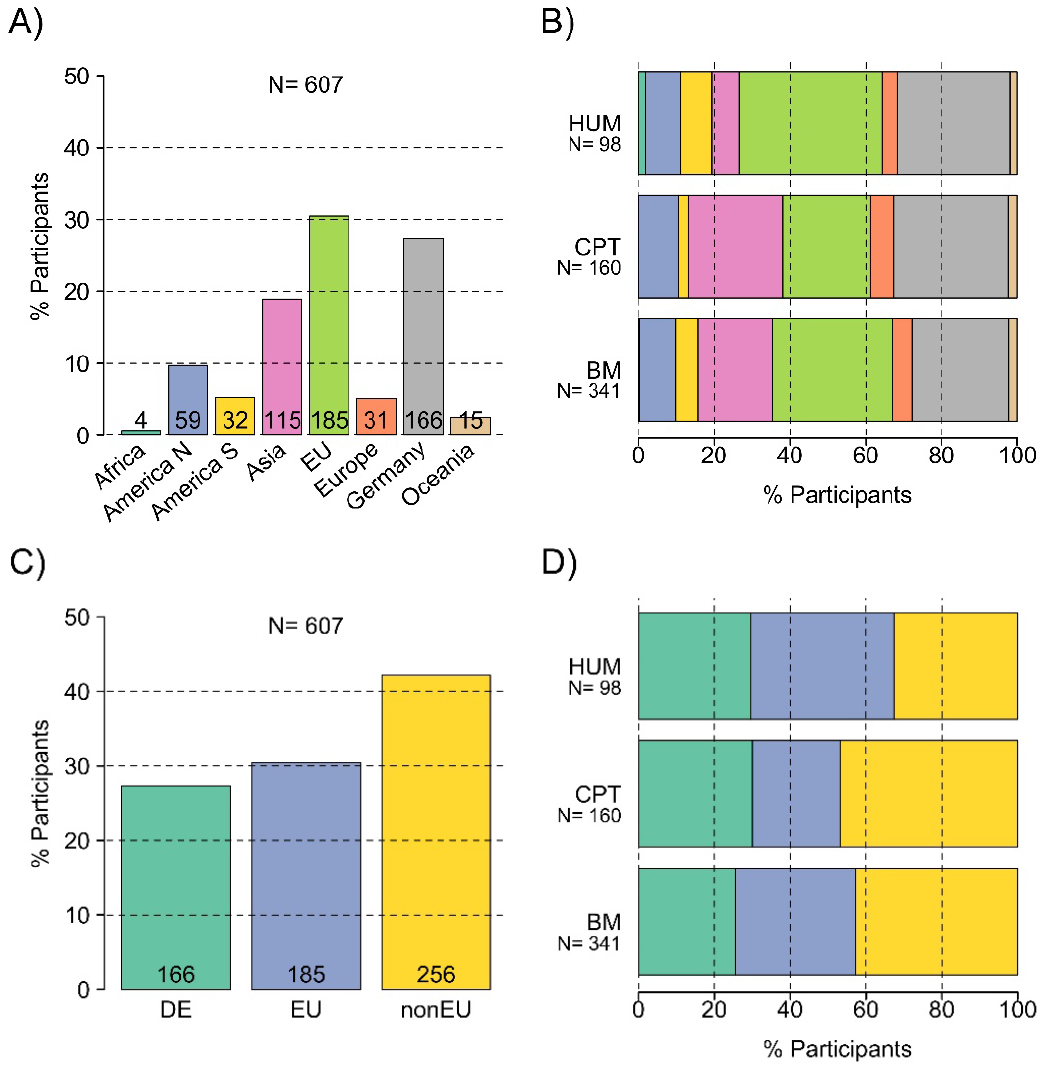
Origins distributions. Origins distributions are given for the MPS overall **(A)** and for each MPS section **(B)**. The data were combined to show the proportion of German postdocs (DE), European Union but non-German postdocs (EU) and postdocs from the rest of the world (nonEU) in the MPS overall **(C)** and in each section **(D)**. Numbers in each bar graph refer to the number of answers given. BM, Biology and Medical section; CPT, Chemistry, Physics and Technology section; HUM, Human Sciences section.

To easily compare postdocs’ origins and their impact on postdocs demographic and work conditions, the regions of origin were merged into three classes: German (DE), EU and non-EU (rest of the world, referred to as nonEU) origins. Overall (**Fig 5C**), the vast majority of postdocs (72.6%, 441/607) were from outside Germany. This was also the case in each MPS section (**Fig 5D**) in which the proportion of international (non-German) postdocs remained above 70%.

When comparing age groups and postdocs’ origins (**Fig 6A**), the proportion of German postdocs increased with age. In the younger age group ([20-30[), only 16.2% of postdocs were German, while in the older age group, 50.9% of postdocs were German. This suggests that the distribution of postdocs’ origins varies strongly with age.

**Figure 6.**
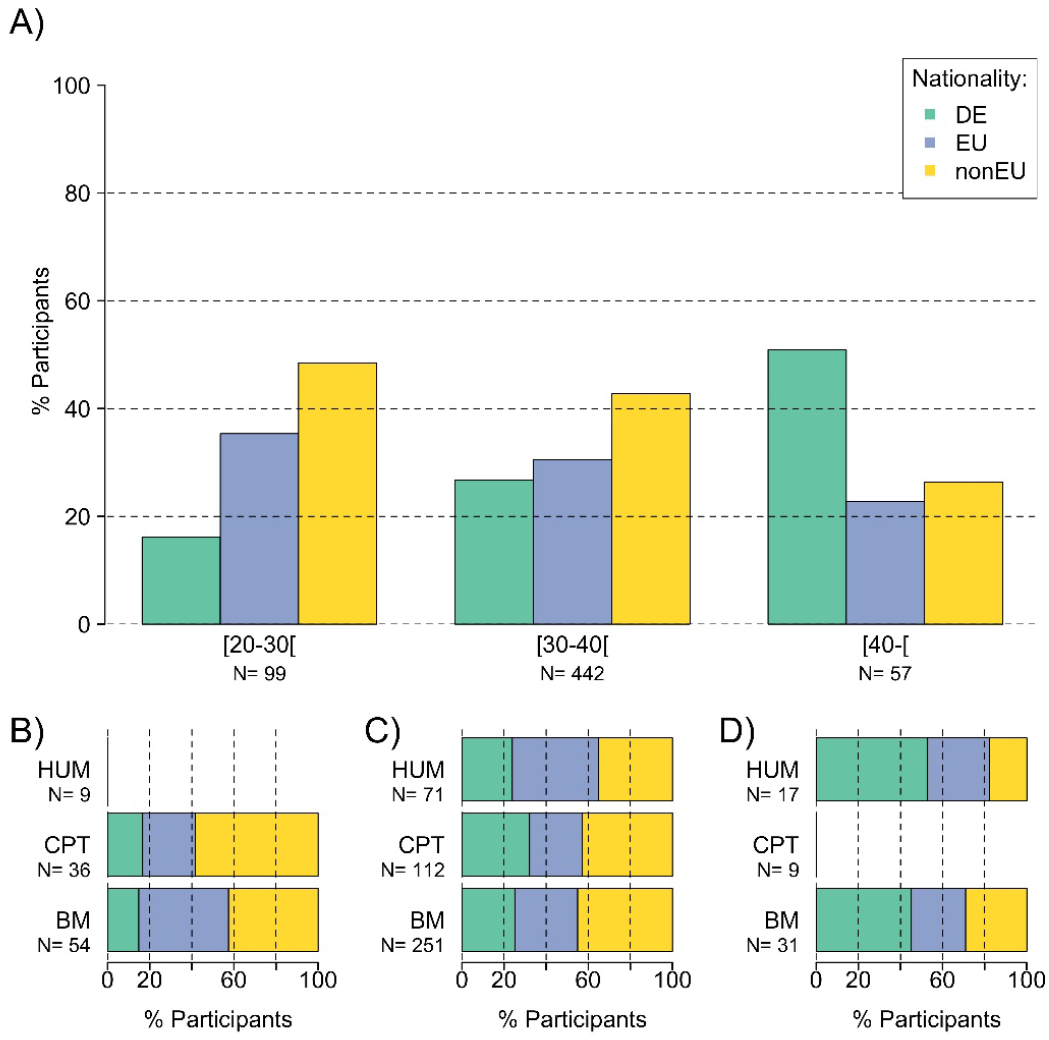
Origins and age group. Comparison of origins distribution based on age group in the MPS overall **(A)**. For each section, the origin distributions are separated by age group in the lower graphs as follows: postdocs from 20 to less than 30 years old **(B)**, postdocs from 30 to less than 40 years old **(C)**, and postdocs from 40 years old on **(D)**. Note, statistics with less than 10 data points are not shown on graphs. DE, Germany; EU, European Union; nonEU, countries outside of the European Union; BM, Biology and Medical section; CPT, Chemistry, Physics and Technology section; HUM, Human Sciences section.

Similarly, within the MPS sections, German postdocs were proportionally less represented in the younger (22%, **Fig 6B**) and middle (32%, **Fig 6C**) age groups. Accordingly, international postdocs represented the majority of both age groups with more than 77% in the younger and more than 67% in the middle age group. German proportions increased to more than 52.9% in the older age group, while the proportion of international postdocs decreased in this age group (**Fig 6D**).

An overall comparison of origin with gender revealed significant differences from the 50% expectation towards more males, which was strongest for the group of postdocs with non-EU origin. Distributions of origin versus gender within sections and a comparison to age can be found in the supplementary Figures S1 and S2.

##### Summary of demographic data

- Significantly more male postdocs than female postdocs
- Majority of postdocs were between 30 and 40 years old
- German postdocs were more represented in the more than 40 years old group
- 73% of postdocs were from outside of Germany

### C. Experience

We surveyed past and present working experiences of the postdoctoral community. Postdocs answered on the geographic region from which they obtained their PhD (referred to as PhD graduation or PhD Lab) by choosing between (i) MPS institutes, (ii) other German institutes, (iii) EU institutions and (iv) non-EU institutions (answers were merged in a comparative analysis). Participants also provided i) their overall postdoctoral experience and ii) their postdoctoral experience within the MPS. For these two questions, respondents had to choose between 4 different categories: (i) less than 1 year, i.e. [0-1[, (ii) 1 to less than 3 years, i.e. [1-3[, (iii) 3 to less than 5 years, i.e. [3-5[and (iv) more than 5 years, i.e. [5-[.

#### 1. Geographic region of PhD graduation

When looking at the geographic region of PhD graduation (also referred to as PhD lab), the majority of postdocs (34.7% – 213/613) graduated in a European country, followed by 30.8% of the postdocs (189/613) who obtained their PhD degrees from non-EU institutions (**Fig 7A**). Less than 20% of the postdocs graduated in Germany (19.6% – 120/613) or within the MPS (14.5% – 91/613).

**Figure 7.**
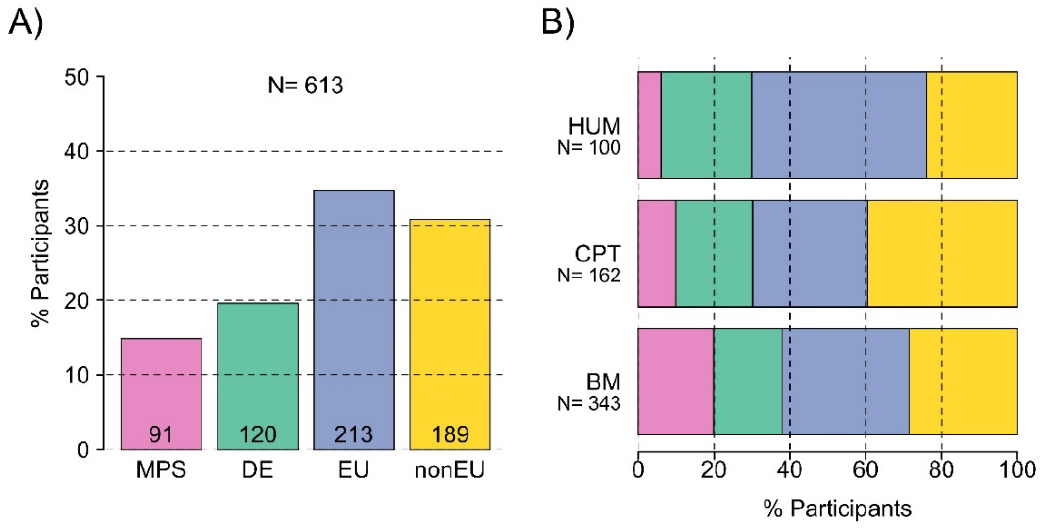
PhD lab. PhD lab distributions are given for the MPS overall **(A)** and for each MPS section **(B)**. Postdocs could choose between fours options: PhD graduation from one MPI **(MPS)**, from a German institution **(DE),** from a European institution **(EU)** or from a non-European institution **(nonEU).** Numbers in each bar graph refer to the number of answers given. BM, Biology and Medical section; CPT, Chemistry, Physics and Technology section; HUM, Human Sciences section; MPS, Max Planck Society.

However, origin of PhD lab varies across MPS sections (**Fig 7B-D**). In the CPT and HUM sections, most postdocs obtained their PhD outside Europe (respectively 39.5% – 64/162 and 46% – 46/100), while in the BM section the proportion of EU and non-EU PhD degrees was relatively similar (respectively 33.5% – 115/343 and 28.6% – 98/343). The BM section had more postdocs that graduated from the MPS (19.8% – 68/343) compared to the CPT (9.9% – 16/162) and the HUM (6% – 6/100) sections. The tendency was reverse for the German PhD degree, as the HUM section had more postdocs that graduated from a German institution (24% – 24/100) compared to the CPT (20.4% – 33/162) and the BM (18.1% – 62/343) sections.

We compared the data obtained on the origin of PhD lab with the postdocs’ origin (**Fig 8A**). Statistical analysis using chi-square test and Cramer’s V estimation showed that these two sets of data were significantly correlated (**Tab S2**).

**Figure 8.**
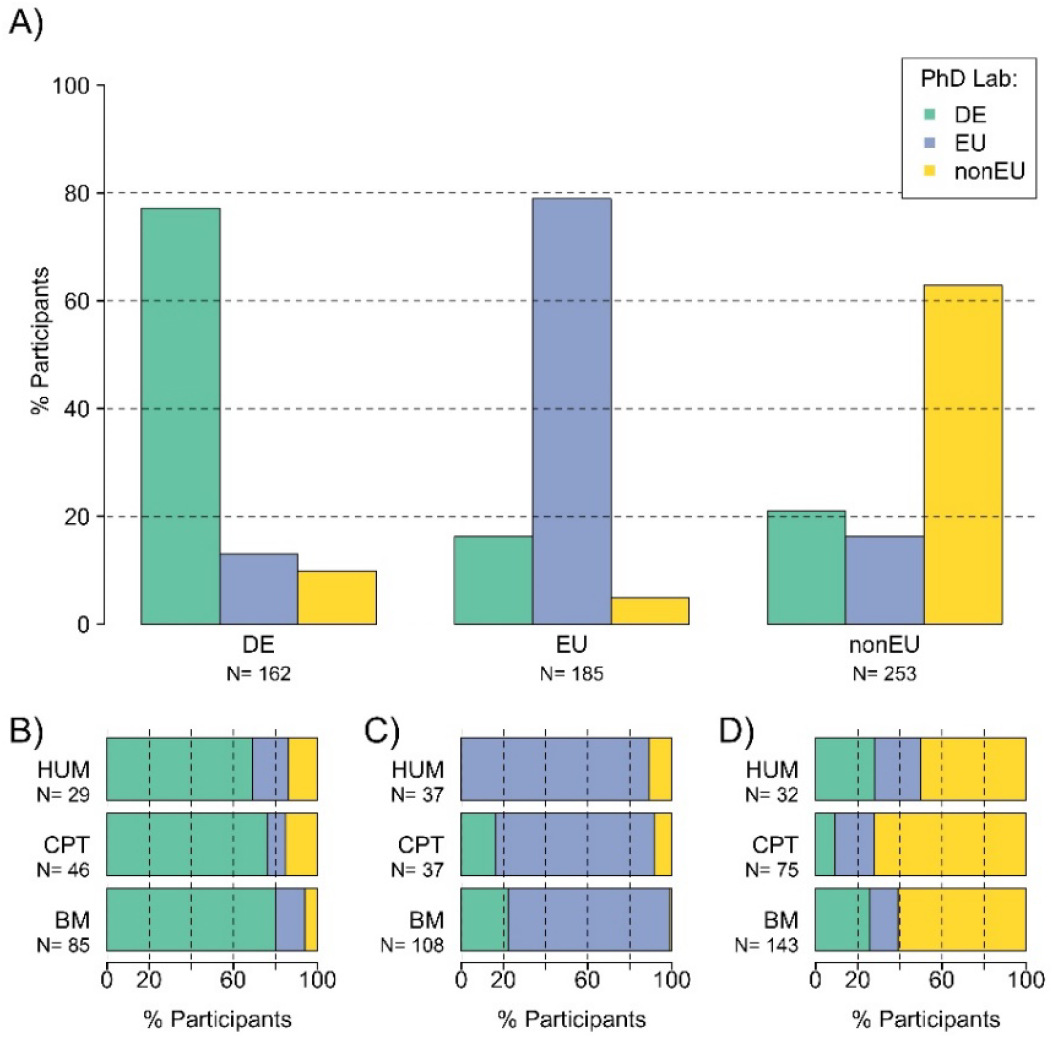
PhD lab and origins. PhD lab distribution in the MPS overall **(A)**, and in each MPS section among Germans **(B)**, Europeans **(C)** or non-Europeans **(D)**. Note the colors of the figure refer to the region where postdocs did their PhD. DE, Germany; EU, European Union; nonEU, countries outside of the European Union; BM, Biology and Medical section; CPT, Chemistry, Physics and Technology section; HUM, Human Sciences section.

German postdocs graduated significantly more from German institutions (77.2% – 125/162) and less from EU (12.9% – 21/162) or non-EU institutions (9.8% – 16/162) (**Fig 8A**). The same observation can be made in each section (**Fig 8B – Tab S3**).

EU postdocs graduated significantly more often from EU institutions (78.9% – 146/185) and more from German institutions (16.2% – 30/185; **Fig 8A**). It was also the case in the MPS sections (**Fig 8C – Tab S3**). Most EU postdocs graduated from EU institutions in all three sections (BM: 83/108; CPT: 28/37; HUM: 33/37). The HUM section differed from the BM and CPT sections regarding German PhD obtained by EU postdocs. Likewise, 22.2% (24/108) and 17.6 % (6/34) of EU postdocs from respectively the BM and CPT section obtained their PhD in a German institution, while no EU postdoc from the HUM section graduated in Germany.

Non-EU postdocs graduated significantly more from non-EU institutions (62.8% – 159/253) and more from German institutions (20.9% – 53/253) than from EU institutions (16.2% – 41/253) (**Fig 8A**). In the MPS sections also (**Fig 8D – Tab S3**), most non-EU postdocs graduated outside EU (BM: 87/143; CPT: 54/75; HUM: 16/32). The CPT section differed from the BM and HUM sections regarding German and EU PhD obtained by non-EU postdocs. In the BM and HUM sections, there were more German PhD holders (BM: 37/143; HUM: 9/32) than EU PhD holders (BM: 19/143; HUM: 7/32). By contrast, in the CPT section, there were more EU PhD holders (14/75) than German PhD holders (7/75).

We compared the data obtained for PhD lab with the postdocs’ age (**Fig S3A**). As already observed in **Fig 6A** for postdocs’ origins, the proportion of German PhD holders increased with increasing age: 25.3% (25/99) were between 20 and 30 years old, while 53.4% (31/58) were older than 40. The tendency was again reverse for the non-German (EU and nonEU) postdocs as 74.7% (74/99) were between 20 and 30 years old, while 46.6% (27/58) were older than 40. As previously observed (see **Fig 3A**), the majority of postdocs were between 30 and 40 and there was no significant difference based on the origin of their PhD.

In the sections, similar distributions to the MPS overall distribution could be observed (**Fig S3B - D**).

We also compared the PhD lab with gender (**Fig S4A**). As the data on origins of postdocs and origins of PhD lab strongly correlated, the distribution was here similar to the distribution observed in **Fig 5A**. Likewise the gender distribution differs from the 50% expectation, with the most uneven distribution for postdocs with non-EU PhD (120 males for 64 females) followed by postdocs with EU (117 males for 91 females) and German PhD (111 males for 96 females).

Performing this analysis for each section separately (**Fig S4B-D**) revealed that the HUM and BM sections had a gender distribution very close to the 50% expectation for postdocs with German and EU PhDs, while for non-EU PhD holders in the CPT section, an uneven gender distribution was observed.

#### 2. Overall experience

Answers to the question “How many years of research experience do you have, excluding your PhD years?” showed that over one third of the postdocs had more than 5 years of experience (35.6% -219/614), followed by postdocs with 1 to 3 (28.1% -173/614), 3 to 5 (22.6% -138/614) and 0 to 1 (13.7% -84/614) years of experience (**Fig 9A**).

**Figure 9.**
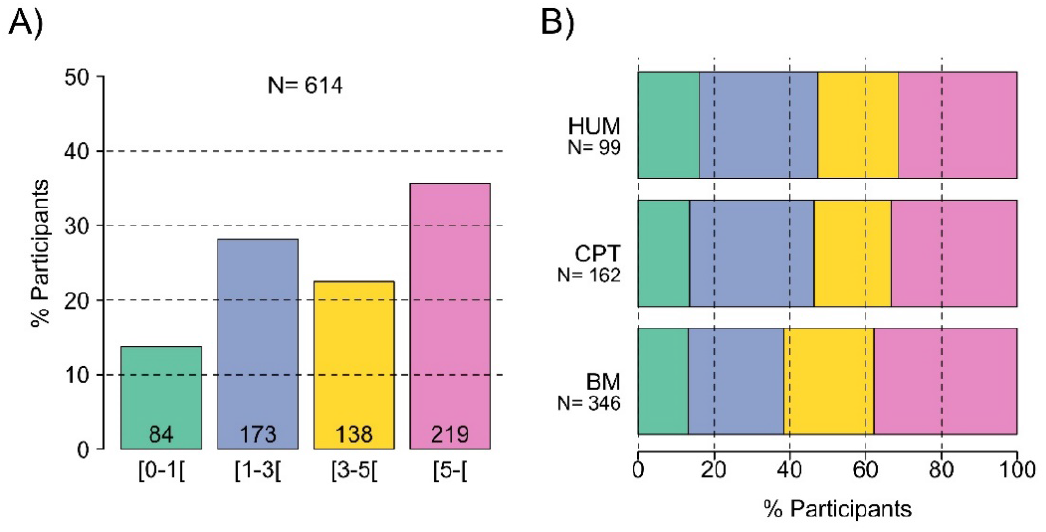
Overall postdoctoral experience. Experience after PhD completion of postdocs in the MPS overall **(A)** and in each section **(B)**. Numbers in each bar graph refer to the number of answers given. BM, Biology and Medical section; CPT, Chemistry, Physics and Technology section; HUM, Human Sciences section.

Similar distributions of years of experience were observed across MPS sections (**Fig 9B**).

With respect to gender (**Fig 10**), a similar proportion of females (49.6%) and males (50.4%) had between 0 to 3 years of experience (i.e. 123/248 and 125/248 respectively; **Fig 10A**). The proportion of females differed significantly from the 50% expectation in the group with more than 3 years of experience (**Tab S4**). Males (64% – 226/353) were significantly more likely to have 3 to more than 5 years of experience than women (35.9 % – 127/353).

**Figure 10.**
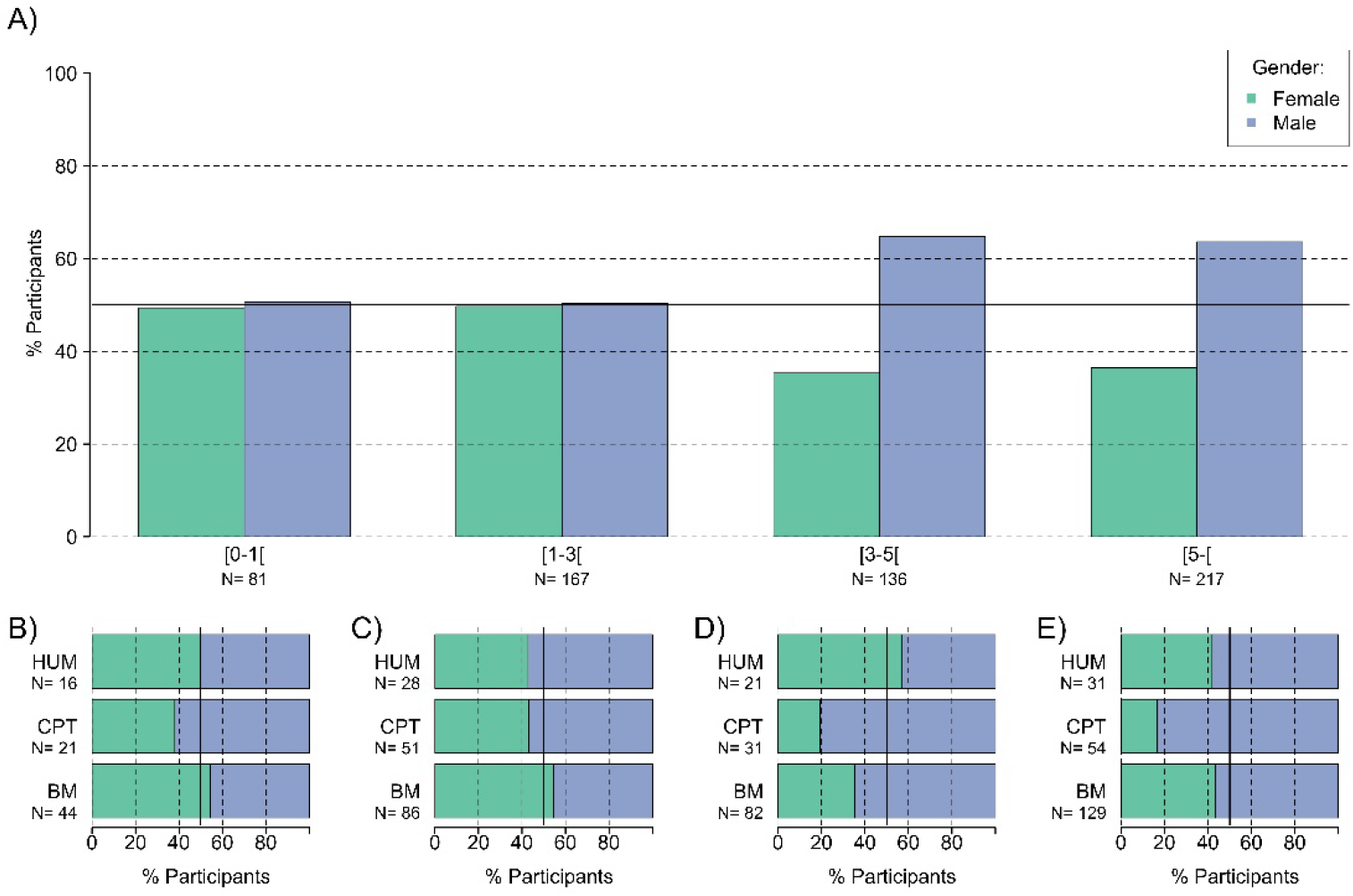
Gender and overall postdoctoral experience. Gender distribution in the MPS overall **(A)** and in each MPS section depending on the overall experiences: less than one year of experience **(B)**, less than 3 years **(C)**, less than 5 years **(D)** and more than 5 years overall experience **(E)**. Overall experience represents the number of years worked after completing the PhD. The continuous line corresponds to 50%. BM, Biology and Medical section; CPT, Chemistry, Physics and Technology section; HUM, Human Sciences section.

For postdocs with less than 3 years of experience, a similar gender distribution was observed across MPS sections (**Fig 10B-C, Tab S4**) with no significant difference from the 50% expectation. Both CPT and HUM sections had more males than females, while the BM section had a slightly higher proportion of females with less than 3 years of experience.

For the more experienced postdocs, there were some variations in the MPS sections (**Fig 10D-E, Tab S4**). In the BM and HUM sections, the gender distributions did not differ significantly from the 50% expectation for postdocs with more than 3 years of postdoctoral experience (**Fig 10D-E**). However, the proportion of males was higher in the BM section (+ 19.4% of males). In the CPT section, the gender distribution significantly deviated from the 50% expectation with more than 61% (6 females for 25 males) and 66% (4 females for 45 males) males in the postdocs with 3 to 5 (**Fig 10D**) and more than 5 years of overall experience respectively (**Fig 10E**). We compared the origin of postdocs depending on their experience (**Fig 11**). As seen in the origins distribution (**Fig 5**), non-German postdocs were the majority throughout all experience groups (**Fig 11A**). The proportion of German postdocs was close to 30% in the groups with 0 to 1, 3 to 5 and more than 5 years of experience (respectively 28% – 23/82; 30.4% – 41/135 and 31% – 68/216) while it was below 20% for postdocs with 1 to 3 years of experience (19.3% – 32/166). The proportion of EU postdocs was quite stable around 30% in all experiences groups (sorted by increasing experience: 26.8% – 22/82; 30.1% – 50/166; 30.4% – 41/135 and 33% – 71/216). Despite a higher proportion of nonEU postdocs in each experience group, the proportion tended to decrease with growing experience (sorted in order of increasing experience: 45.1% – 37/82; 50.6% – 84/166; 39.3% – 53/135 and 36% – 77/216).

**Figure 11.**
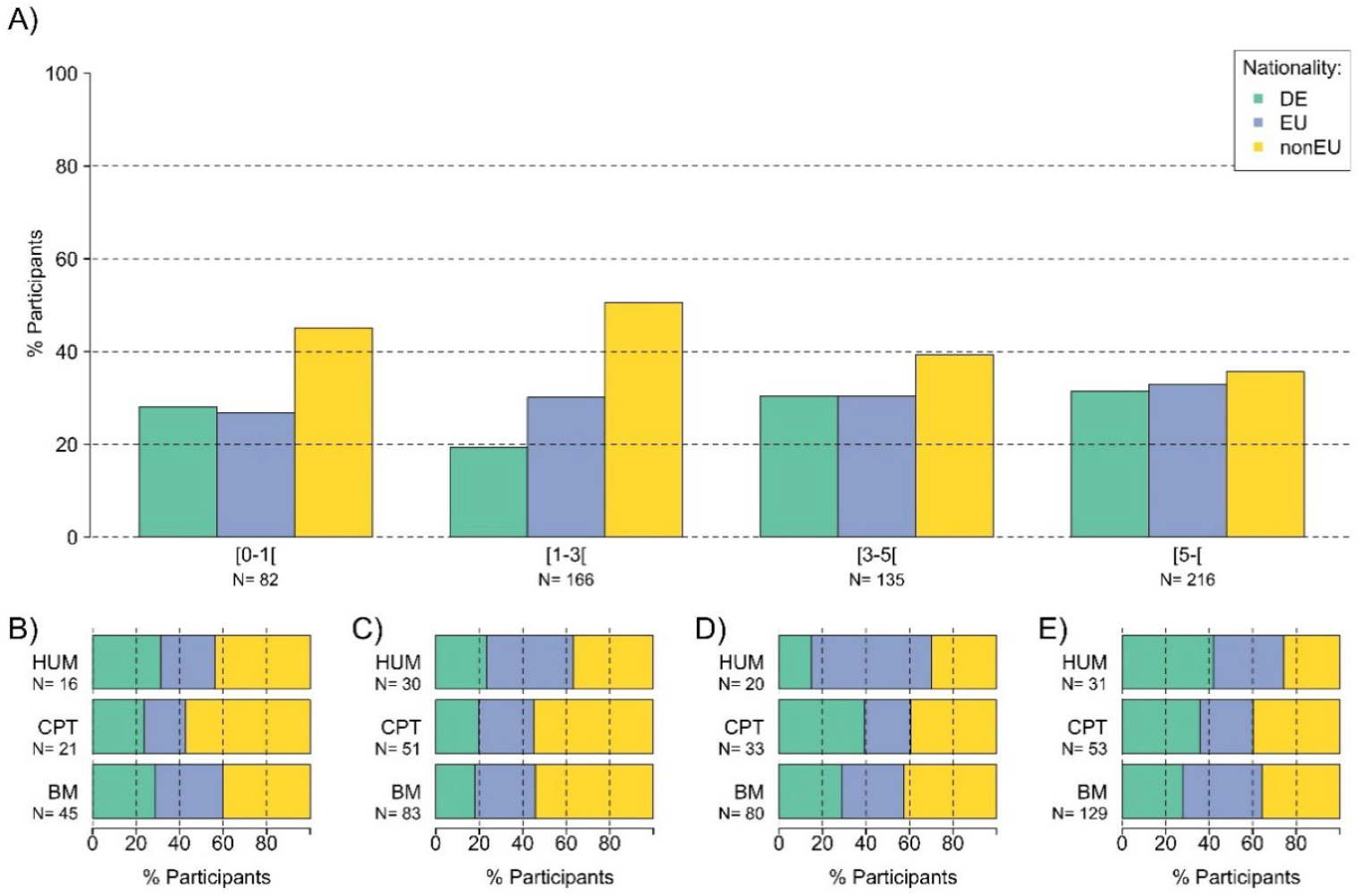
Origins and overall experience. Origin distribution in the MPS overall **(A)** and in each MPS section depending on the overall experiences: less than one year of experience **(B)**, less than 3 years **(C)**, less than 5 years **(D)** and more than 5 years overall experience **(E)**. Overall experience represents the number of years worked after completing the PhD. DE, Germany; EU, European Union; nonEU, countries outside of the European Union; BM, Biology and Medical section; CPT, Chemistry, Physics and Technology section; HUM, Human Sciences section.

The MPS sections showed only minor variations from the results of the overall population (**Fig 11B-E**). Specifically, in the CPT section, the proportion of postdocs with 3 to 5 years of experience was identical for non-EU and German postdocs (39.4% – 13/33), while the HUM section had a majority of EU postdocs (55% – 11/20) within the same experience group. In the HUM section, postdocs with more than 5 years of experience were mainly from Germany (41.9% – 13/31).

We also examined the distributions of postdocs based on their years of experience as a function of where they completed their PhD (i.e., PhD lab; **Fig 12**). Postdocs with 0 to 1 and 3 to 5 years of experience more likely graduated from a German institution (39.3% – 33/84 and 39.1% – 54/138 respectively). For these two experience groups, the proportion of postdocs who had obtained their PhD from an EU institution was the second largest (respectively 32.1% – 27/84 and 32.6% – 45/138). In the group with 1 to 3 years of experience, postdocs graduated more from non-EU (37.5% – 63/168) and EU institutions (36.3% – 61/168). On the contrary, in the group with more than 5 years of experience, postdocs graduated more from EU (37% – 79/215) and German institutions (36% – 77/215).

**Figure 12.**
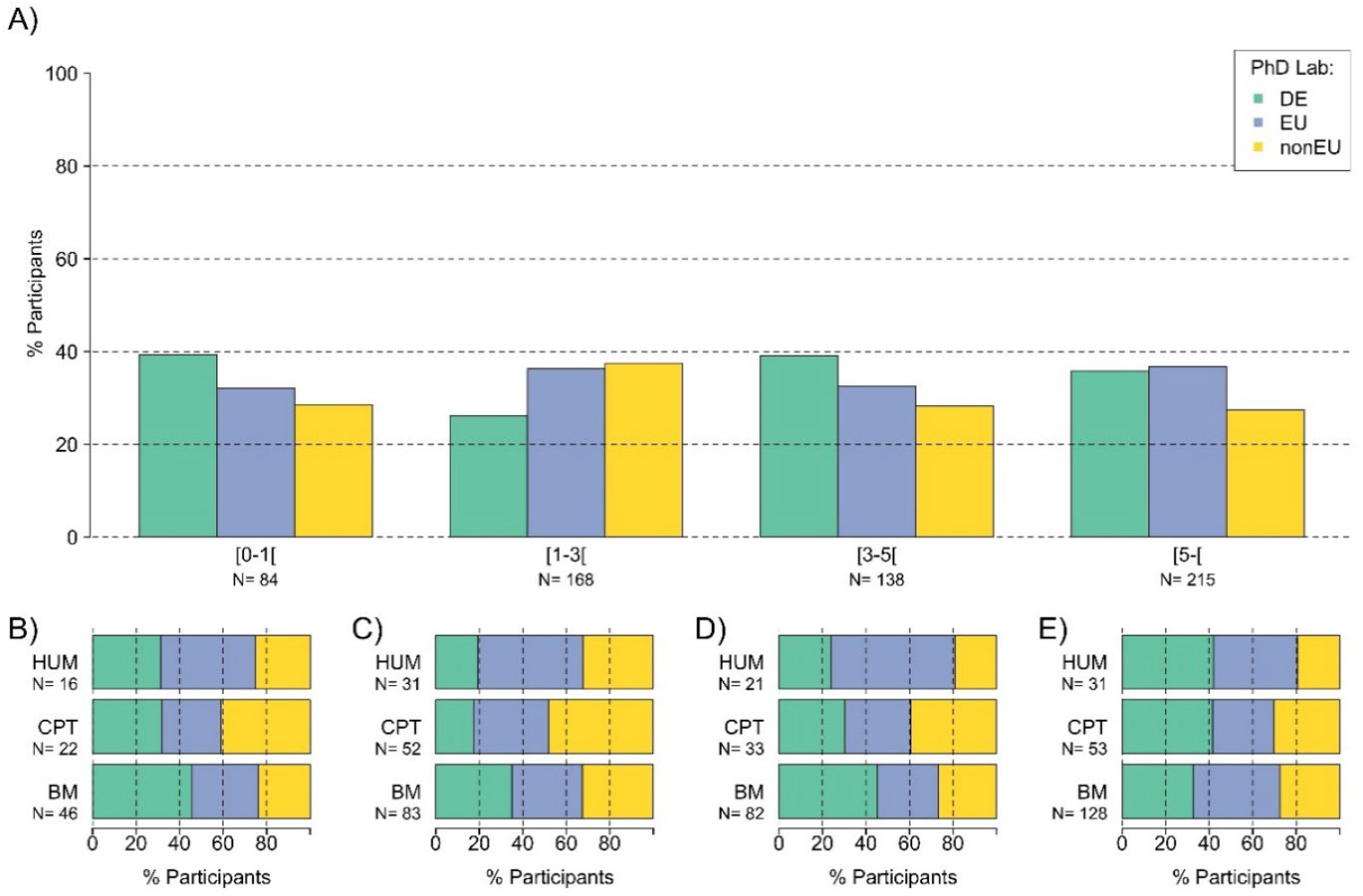
PhD lab and overall experience. PhD lab distributions in the MPS overall **(A)** and in each MPS section depending on the overall experiences: less than one year of experience **(B)**, less than 3 years **(C)**, less than 5 years **(D)** and more than 5 years overall experience **(E)**. Overall experience represents the number of years worked after PhD graduation. Note the colors of the figure refer to the region where postdocs obtained their PhD. DE, German institutions; EU, European Union institutions; nonEU, countries outside of the European Union institutions. BM, Biology and Medical section; CPT, Chemistry, Physics and Technology section; HUM, Human Sciences section.

The MPS sections showed only minor variations from the results of the overall population (**Fig 12B-E**). For postdocs with less than a year of experience (**Fig 12B**), non-EU PhDs represented the majority in the CPT section (40.9% – 9/22) while the proportion of EU PhDs was the highest in the HUM section (43.8% – 7/16). For postdocs with 1 to 3 years of experience (**Fig 12C**), the proportion of German PhDs was the highest in the BM section (34.9% – 29/83). For postdocs with 3 to 5 years of experience (**Fig 12D**), nonEU PhDs represented the majority in the CPT section (39.4% – 13/33). For postdocs with more than 5 years of experience (**Fig 12E**), German PhDs represented the majority in the CPT and HUM sections (41.5% – 22/53 and 41.9% – 13/31 respectively).

#### 3. MPS experience

Answers to the question “How long have you been working at the MPS as a Postdoc” (referred to as MPS experience) showed that most postdocs had been in a MPI for 1 to 3 years (39.9% – 244/621), followed by postdocs with less than a year of MPS experience (24.3% -149/612). Postdocs with 3 to 5 (19.9% -122/612) and more than 5 (15.8% -97/612) years of experience were the least (**Fig 13A**). Similar distributions were observed across MPS sections (**Fig 13B**).

**Figure 13.**
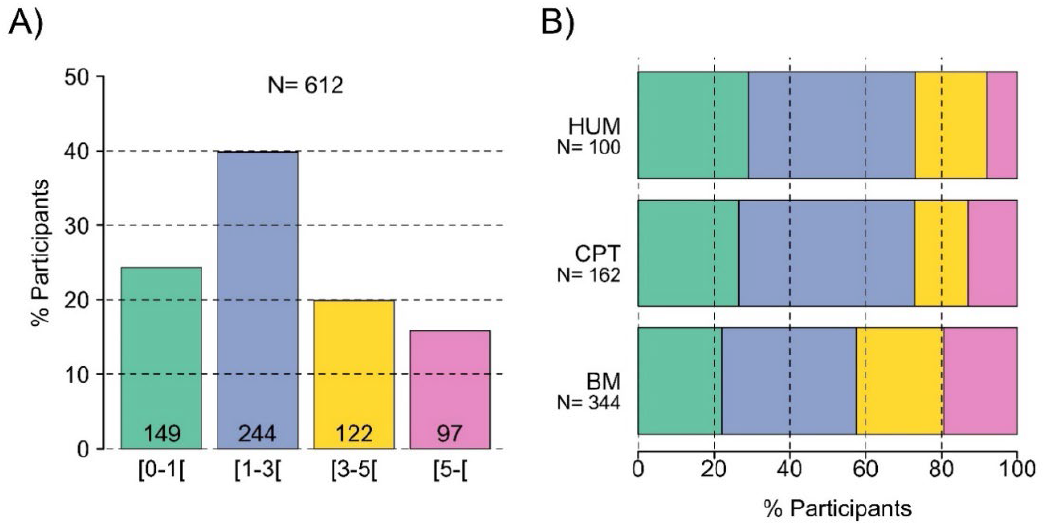
MPS postdoctoral experience. Experience within the MPS after PhD graduation of postdocs in the MPS overall **(A)** and in each section **(B)**. Numbers in each bar graph refer to the number of answers given. BM, Biology and Medical section; CPT, Chemistry, Physics and Technology section; HUM, Human Sciences section.

Comparing postdocs’ overall experience with their MPS experience (**Fig 14A**), we found that the two group of experience were significantly correlated (chi-square test and Cramer’s V estimation; **Tab S5**).

**Figure 14.**
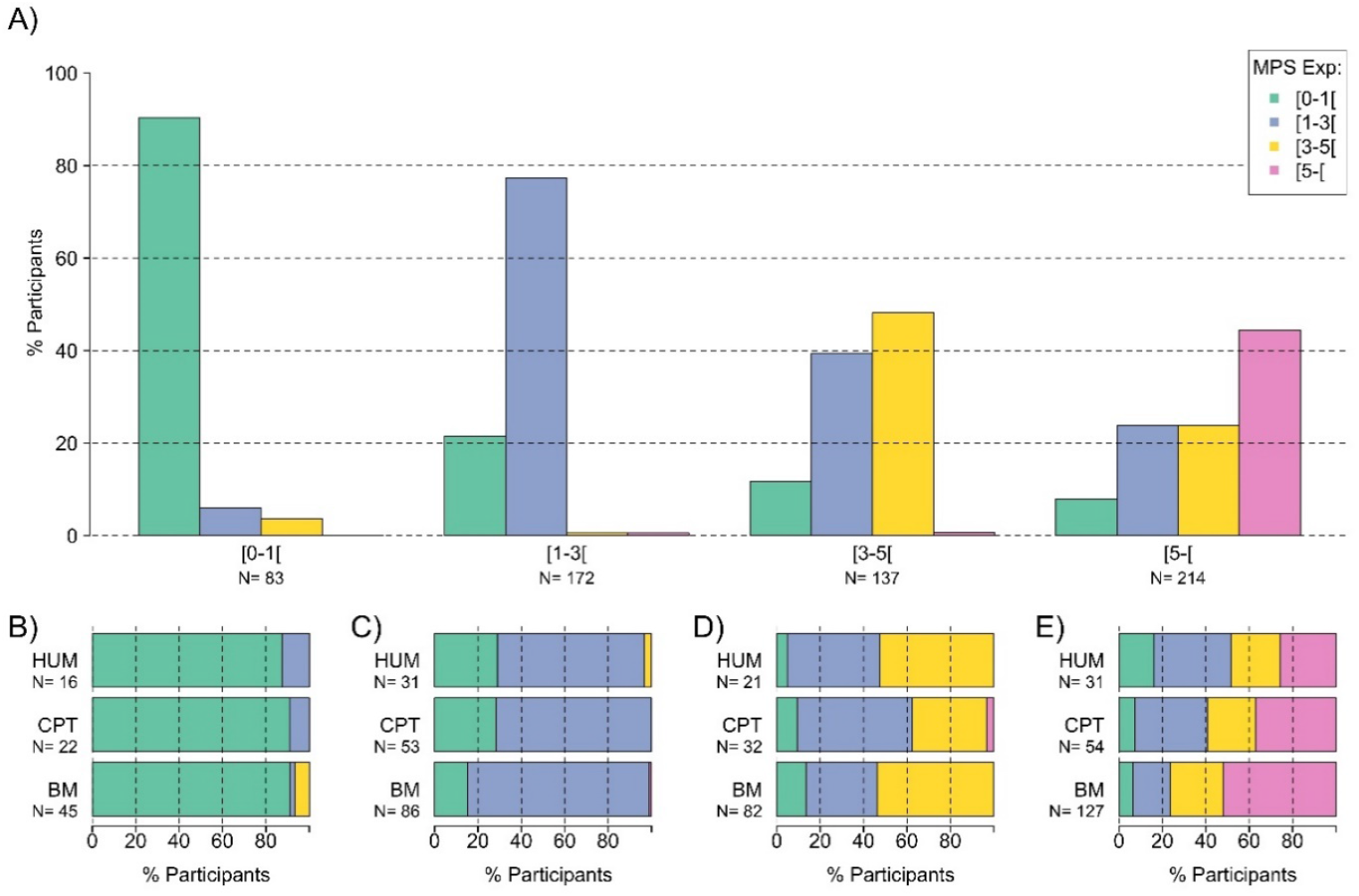
MPS and overall postdoctoral experience. Comparison of postdocs distribution based on their MPS postdoctoral experience in the MPS overall **(A)** and in each MPS section depending on their overall experiences: less than one year of experience **(B)**, less than 3 years **(C)**, less than 5 years **(D)** and more than 5 years overall experience **(E)**. Overall and MPS experiences represent the number of years worked after PhD graduation. BM, Biology and Medical section; CPT, Chemistry, Physics and Technology section; HUM, Human Sciences section.

With respect to their gender (**Fig 15A**), more male than female postdocs were found for postdocs with less than a year (+19.4% males), 1 to 3 (+3.4% males) and more than 5 (+15.5% males) years of MPS experience. This proportion differed significantly from the 50% expectation only in the group of postdocs with 3 to 5 years of MPS experience (+ 39.5% males; Chisq to 50% p<0.01 **Tab S6**).

**Figure 15.**
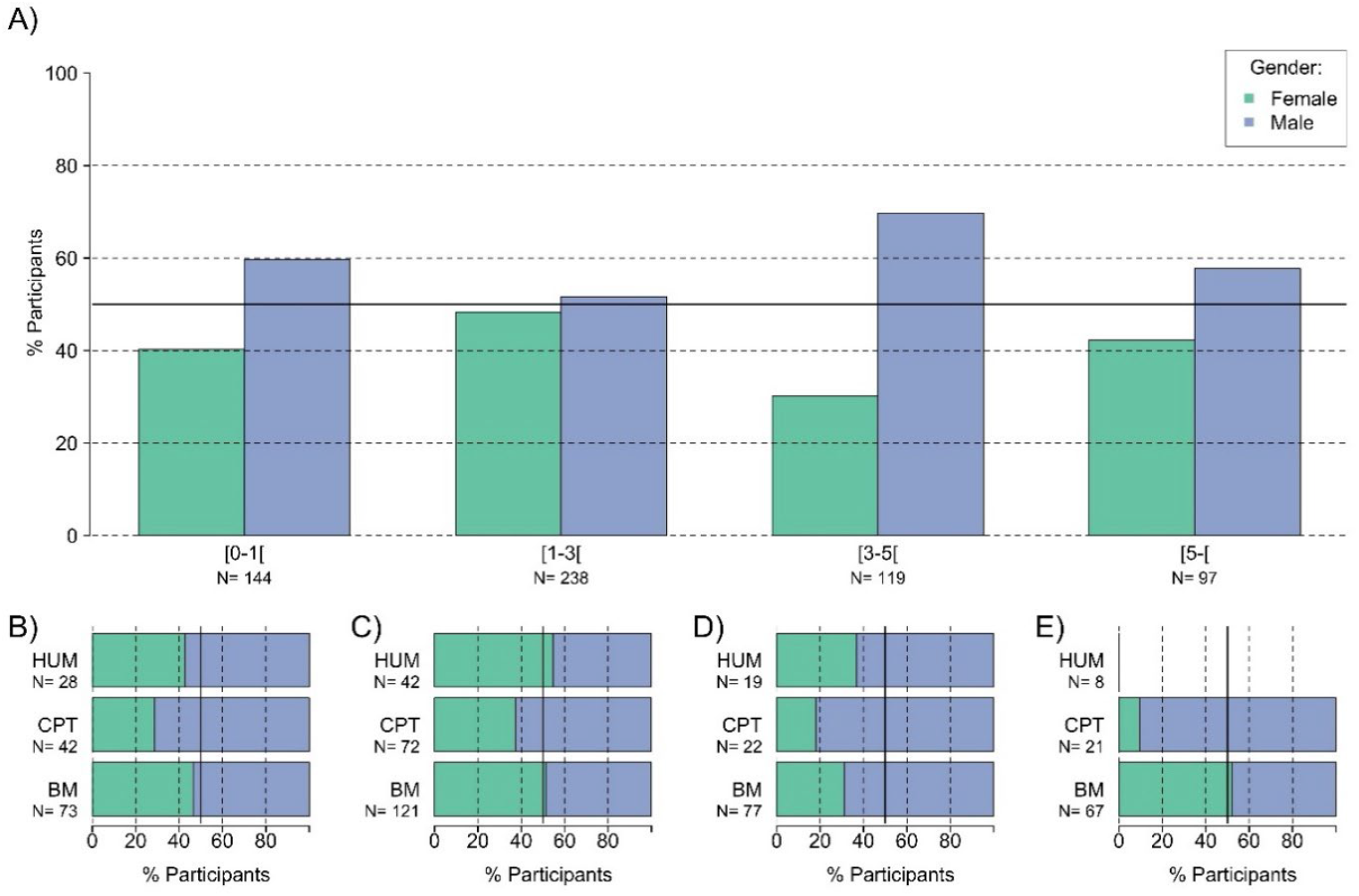
Gender and MPS postdoctoral experience. Gender distribution in the MPS overall **(A)** and in each MPS section depending on the MPS experiences: less than one year of experience **(B)**, less than 3 years **(C)**, less than 5 years **(D)** and more than 5 years of MPS experience **(E)**. MPS experience represents the number of years worked within the MPS after PhD graduation. The thick line corresponds to 50%. Note, statistics with less than 10 data points are not shown on graphs. BM, Biology and Medical section; CPT, Chemistry, Physics and Technology section; HUM, Human Sciences section.

Similar gender distributions were observed across sections, even though the CPT section stood out as having the highest male bias (**Fig 15B-E, Tab S6**). In the BM section, there were two MPS experience groups that had more female than male postdocs. The group with 1 to 3 years (62/121 females) and the group with more than 5 years (35/67 females) of MPS experience (**Fig 15C & E**). In the HUM section, the group with 1 to 3 years of MPS experience had 54.7% (23/42) female postdocs (**Fig 15C**).

Further, as seen in the previous origins distribution (see **Fig 5**), international postdocs represented the majority of postdocs with less than 3 years of MPS experience (75.5% – 289/383; **Fig 16A**).

**Figure 16.**
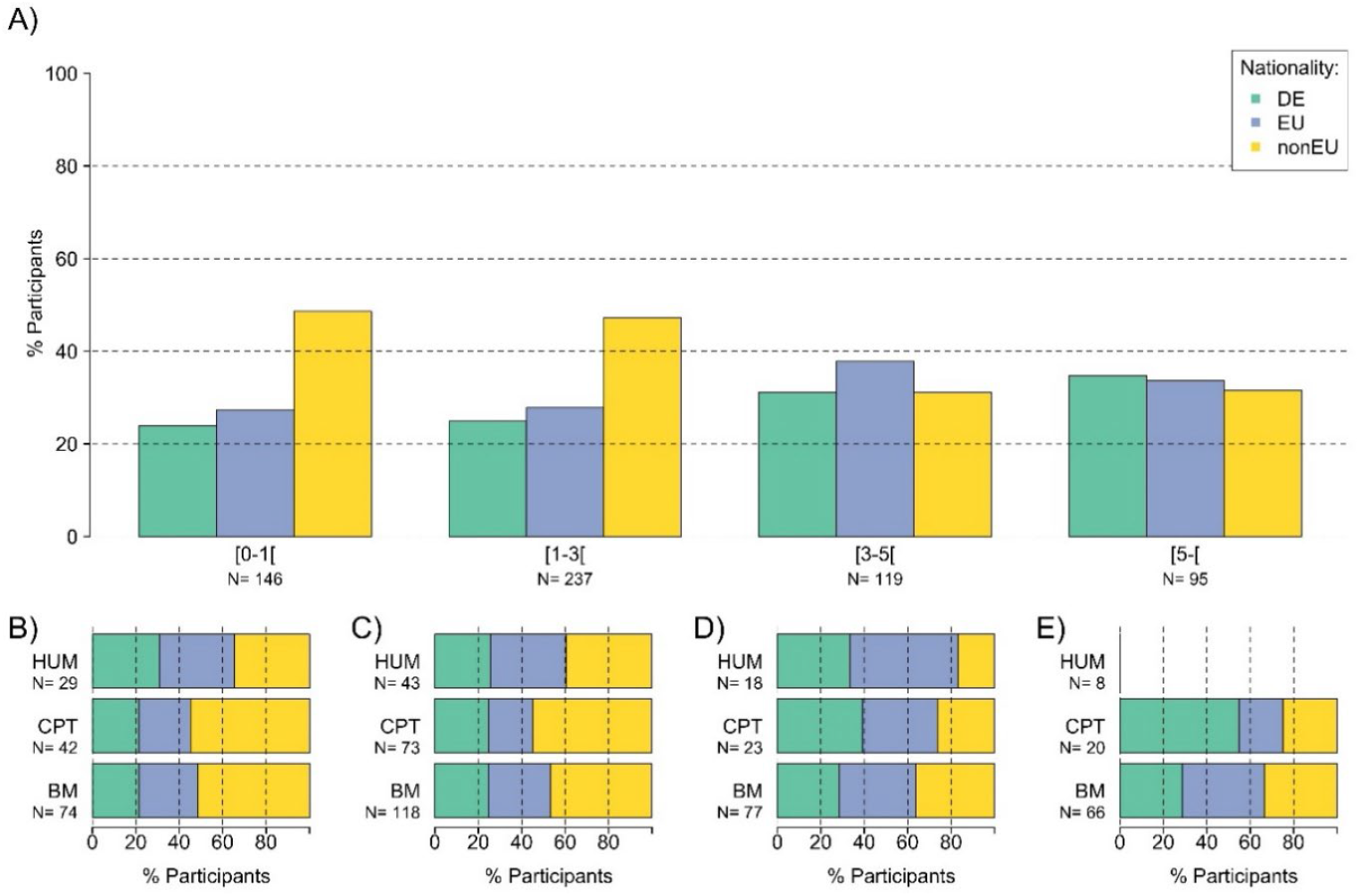
Origins and MPS experience. Origin distribution in the MPS overall **(A)** and in each MPS section depending on the MPS experiences: less than one year of experience **(B)**, less than 3 years **(C)**, less than 5 years **(D)** and more than 5 years of MPS experience **(E)**. MPS experience represents the number of years worked within the MPS after PhD graduation. Note, statistics with less than 10 data points are not shown on graphs. DE, Germany; EU, European Union; nonEU, countries outside of the European Union; BM, Biology and Medical section; CPT, Chemistry, Physics and Technology section; HUM, Human Sciences section.

For postdocs with less than or more than 3 years of MPS experience, the proportion of Germans was 24.5% (94/383) and 32.7% (70/214), respectively. Similar distributions were observed across sections (**Fig 16B-E**).

With respect to their PhD lab (**Fig 17A**), about 20% of postdocs with either less than a year or 3 to 5 years of MPS experience did their PhD in one MPI (18.9% – 28/148 and 19% – 23/121, respectively). This proportion dropped to around 10% for postdocs with either 1 to 3 years or more than 5 years of MPS experience (11.7% – 28/240 and 12% – 12/97, respectively). The proportion of postdocs that graduated from a German or an EU institution increased for postdocs with more years of MPS experience (for German PhD lab: 13.5% – 20/148; 20% – 48/240; 22.3% – 27/121 and 25% – 24/97; for EU PhD lab: 35.1% – 52/148; 32.5% – 78/240; 34.7% – 42/121 and 40% – 39/97). On the contrary, the proportion of postdocs that graduated from a non-EU institution decreased for postdocs with more years of MPS experience (32.4% – 48/148; 35.8% – 86/240; 24% – 29/121 and 23% – 22/97). Similar distributions were observed across sections (**Fig 17B-E**).

**Figure 17.**
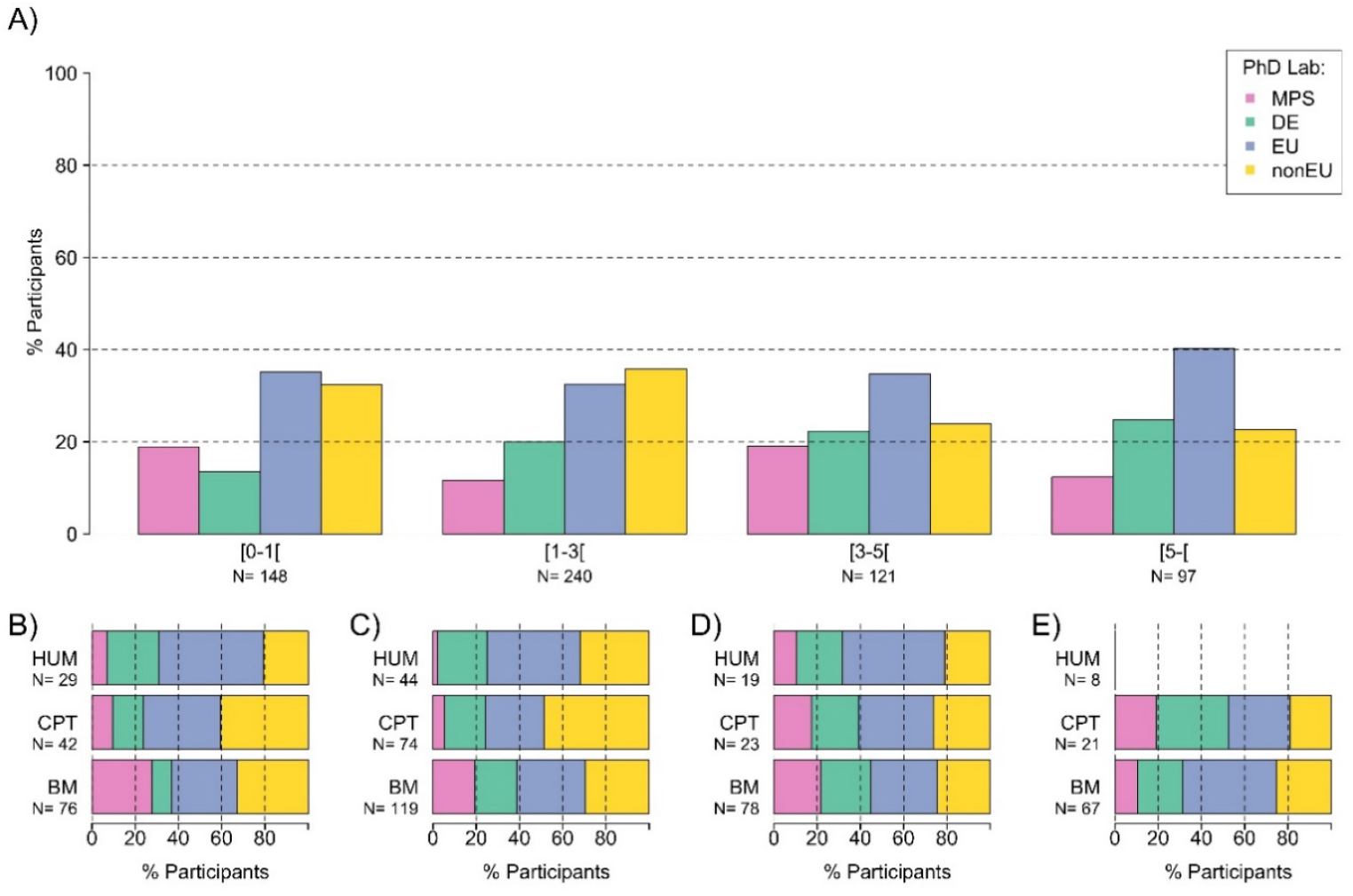
PhD lab and MPS experience. PhD lab distributions in the MPS overall **(A)** and in each MPS section depending on the MPS experiences: less than one year of experience **(B)**, less than 3 years **(C)**, less than 5 years **(D)** and more than 5 years of MPS experience **(E)**. MPS experience represents the number of years worked within the MPS after PhD graduation. Postdocs could choose between fours options: graduation from one MPI (MPS), from a German institution (DE), from a European institution (EU) or from a non-European institution (nonEU). Note, statistics with less than 10 data points are not shown on graphs. BM, Biology and Medical section; CPT, Chemistry, Physics and Technology section; HUM, Human Sciences section.

##### Summary of experience data

- Strong correlation between postdocs’ origin and the geographic region of PhD graduation
- Strong correlation between postdocs’ ages and their postdoctoral experience
- Over one third of postdocs had more than 5 years of postdoctoral experience
- Significantly fewer females postdocs with more than 3 years of postdoctoral experience
- Male postdocs had slightly more postdoctoral experience than female postdocs
- German postdocs had slightly more postdoctoral experience than non-German postdocs

### D. Employment conditions in the MPS

To assess the current employment situation of postdocs, participants could choose between three different employment conditions:

1. **on a stipend** defined as “a payment made for living expenses (e.g. scholarship, fellowship or grant). As it is not considered wages, you are not paying for Social Security or Medical Taxes on it and your employer will not withhold any income taxes from the stipend. However, it can still be counted as taxable income for income tax purposes in certain cases”. For description and analysis, this group was later separated between postdocs employed on a MPS scholarship (also referred to as scholarship) and postdocs employed on a third-party fellowship (also referred to as fellowship).
2. **on a fixed-term contract** defined as “governed by the Collective Wage Agreement for the Civil Service (i.e. TVöD); referred to as fixed-term contract or contract. You pay taxes and social contributions (e.g. health insurance, unemployment money, retirement money) on your salary (pre *vs* post taxes) and the institute pays half of your health insurance. Your contract is time-limited.”
3. **on a permanent employment contract** defined as “same as fixed-term but without end”.

*Note: the definitions above were established by the PostdocNet and do not engage in any case the responsibility of the MPS.*

#### 1. General employment conditions

The vast majority of postdocs (84.1% – 508/604) were employed on a fixed-term contract, while 13.6% (82/604) of postdocs received a stipend and 2.3% (14/604) of postdocs were employed on a permanent contract (**Fig 18A**).

**Figure 18.**
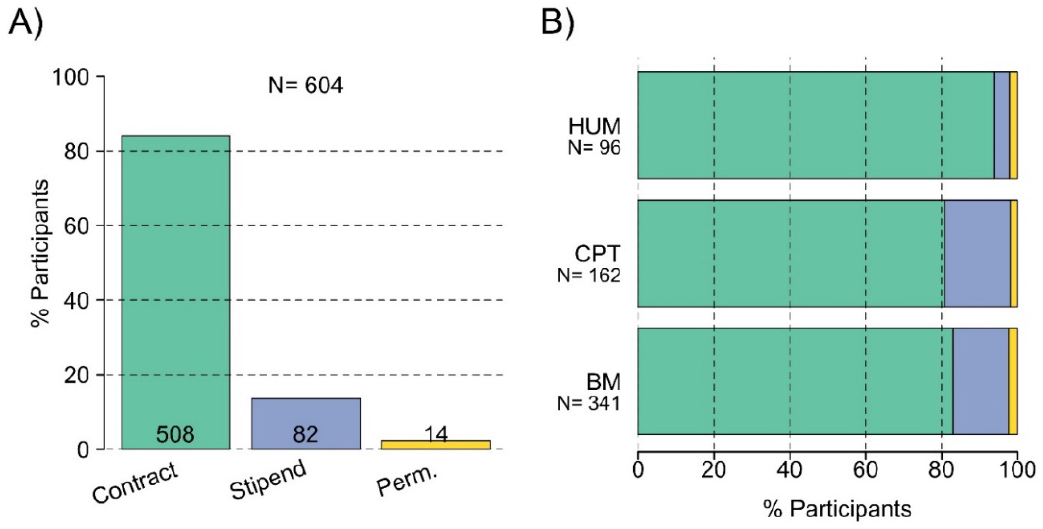
Employment conditions. Employment conditions in the MPS overall **(A)** and in each section **(B)**. Numbers in each bar graph refer to the number of answers given. Contract, fixed term TVöD-based employment; stipend, scholarship or fellowship; perm, permanent TVöD-based employment. BM, Biology and Medical section; CPT, Chemistry, Physics and Technology section; HUM, Human Sciences section.

Variations in the employment conditions were observed across MPS sections (p<0.01, Cramer’s V = 12.7%). While all sections showed a similar number of permanent positions (2.3% – 8/341 for BM; 1.9% – 3/162 for CPT; 2.1% – 2/96 for HUM), the proportion of stipends varied from 4.2% (4/96) in the HUM, to 14.7% (50/341) in the BM and 17.3% (28/162) in the CPT sections (**Fig 18B**).

Scientists on permanent positions represented a very small number of participants (2.3%), as they were not the target of this survey. They were excluded from further analysis, to avoid drawing biased conclusions based on an insufficient sample size (see Discussion).

##### a) Employment conditions & origins

A strong correlation (p<0.001, Cramer’s V = 27.6%) between type of employment and origin was observed (**Fig 19A**). German postdocs were rarely employed on a stipend (1.9% – 3/155) while the situation was more common for Europeans (9.7% – 17/175) and even more frequent for non-Europeans (24.4% – 60/246).

**Figure 19.**
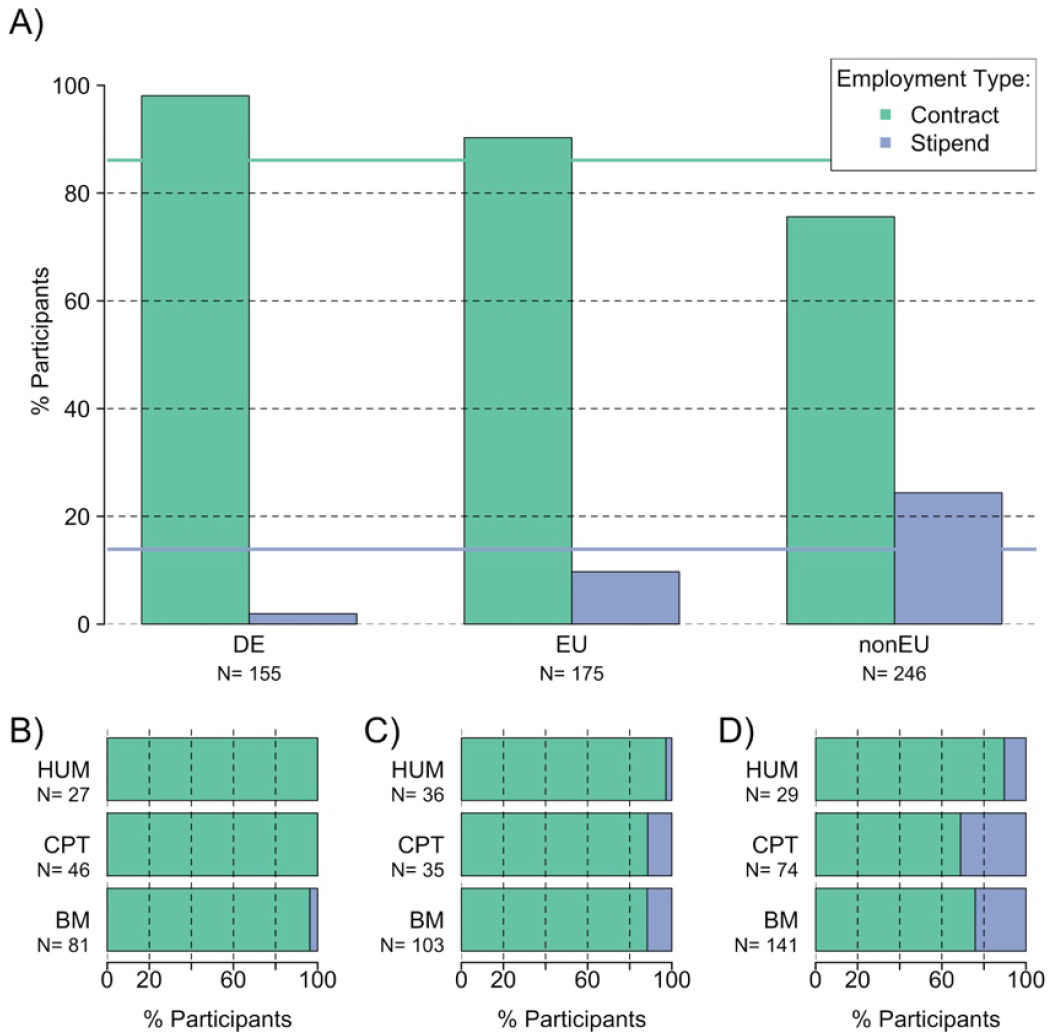
Employment conditions and origins. Distribution of employment conditions in the MPS **(A)** and in each MPS section depending on the origin: German **(B** - DE); European **(C** - EU); Non-European **(D** - nonEU). The green and blue lines correspond to the proportion in the overall population of fixed term TVöD-based contracts and stipends respectively. BM, Biology and Medical section; CPT, Chemistry, Physics and Technology section; HUM, Human Sciences section.

In the MPS sections, the distribution of fixed-term contracts was similar to the overall population. The small number of German postdocs on stipends exclusively belonged to the BM section (**Fig 19B**), while the majority of European stipend holders belonged equally to the CPT and BM sections and only a small fraction was working in the HUM section (**Fig 19C**). Finally, the comparison revealed that the highest number of non-European stipend holders worked in the CPT section, followed by the BM section. Again, the smallest number of non-European stipend holders was found in the HUM section (**Fig 19D**).

As expected from the previously observed correlation between origin and PhD lab, we additionally observed a correlation between employment conditions and PhD Lab (p<0.001, Cramer’s V = 28.4% – **Fig S5**). Postdocs that graduated in Germany were rarely employed on a stipend (5.7% – 5/87 for MPS PhD; 3.6% – 4/110 for other German PhD) while this was more common for postdocs that graduated in Europe (10.6% – 21/199) and frequent for postdocs that graduated outside of Europe (28.1% – 52/185). A section-wise comparison revealed similar results.

##### a) Employment conditions & gender

We examined the relationship between gender and employment type (**Fig 20A**). A slightly higher proportion of male postdocs were employed on a stipend (15.1% – 51/337) compared to female postdocs (10.9% – 26/238) but this difference did not reach significance (p>0.1).

**Figure 20.**
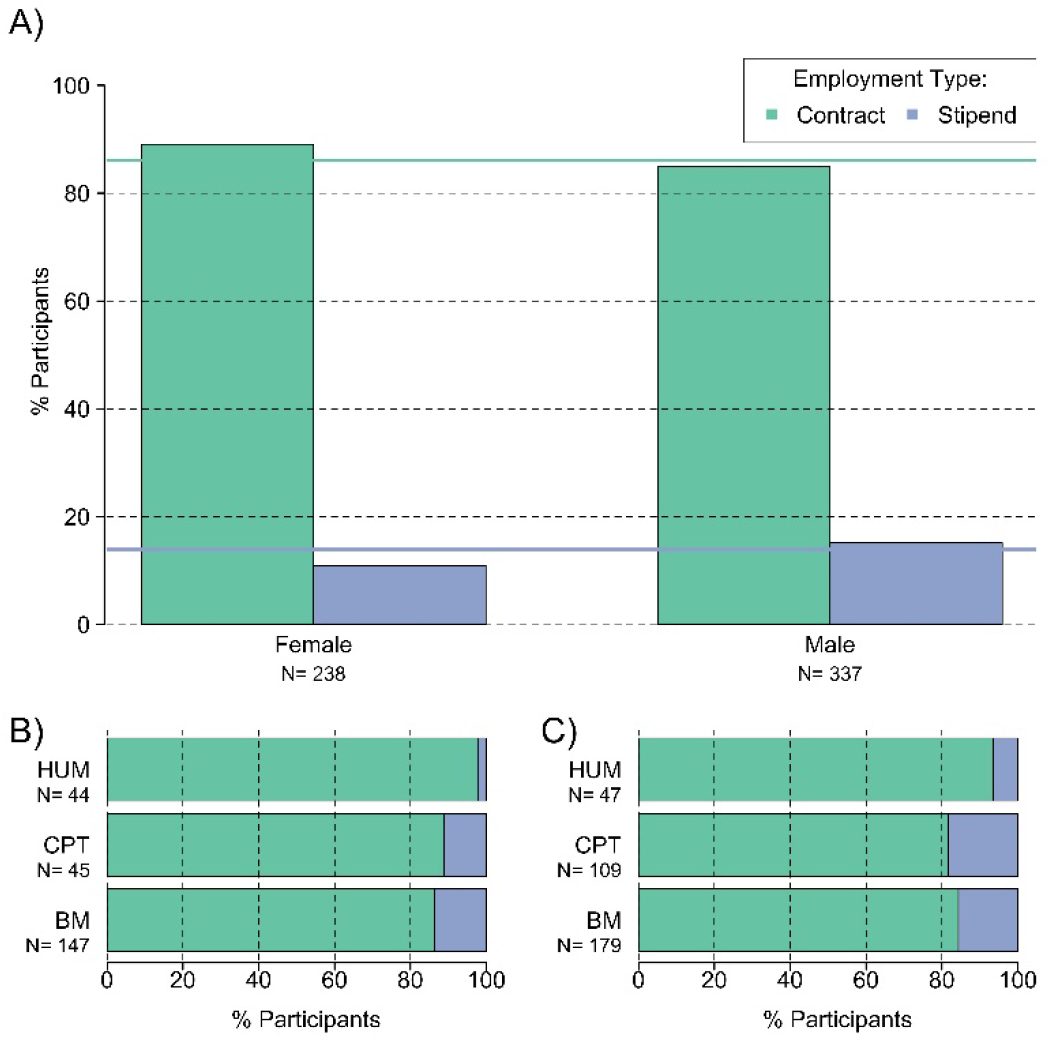
Employment conditions and gender. Distribution of employment conditions in the MPS **(A)** and in each MPS section depending on the gender: female **(B)** and male **(C)**. The green and blue lines correspond to the proportion in the overall population of fixed term TVöD-based contracts and stipends respectively. BM, Biology and Medical section; CPT, Chemistry, Physics and Technology section; HUM, Human Sciences section.

Within the sections (**Fig 20B-C**), the distribution of employment type was similar to the overall population (**Fig 18B**), with no significant differences in employment type according to gender.

##### b) Employment conditions & age and experience

A comparison of the employment type with age (**Fig 21A**) showed a significant correlation (p<0.001, Cramer’s V = 19.9%). Younger postdocs were more likely to be employed on stipends than older postdocs. For stipend holders, 27.7% (26/94) belonged to the younger age group between 20 and 30 years, 11.6% (51/440) belonged to the middle age group between 30 and 40 years and only 2.1% (1/48) were older than 40 years old.

**Figure 21.**
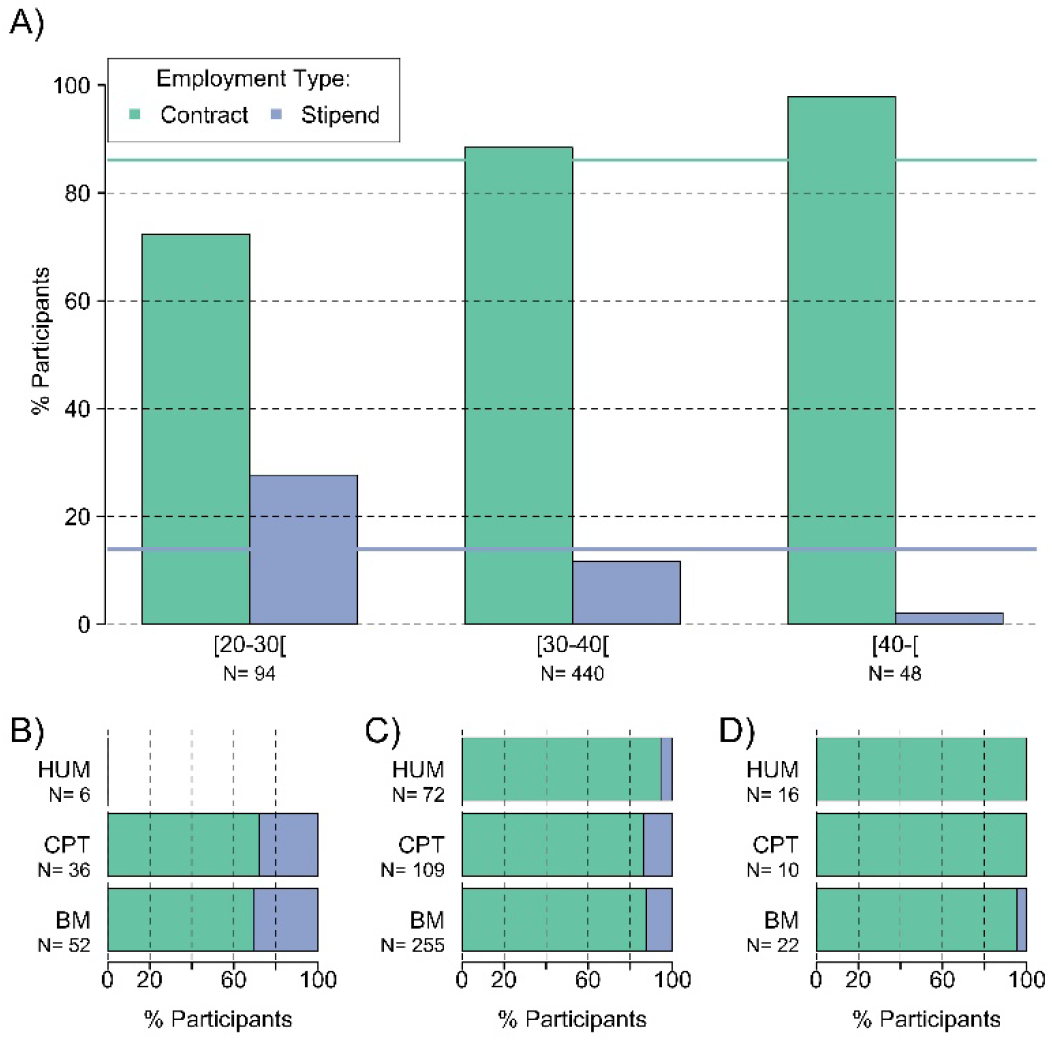
Employment conditions and age group. Distribution of employment conditions in the MPS **(A)** and in each MPS section depending on the age group: postdocs from 20 to less than 30 years old **(B)**, postdocs from 30 to less than 40 years old **(C)**, and postdocs from 40 years old on **(D)**. The green and blue lines correspond to the proportion in the overall population of fixed term TVöD-based contracts and stipends respectively. Note, statistics with less than 10 data points are not shown on graphs. BM, Biology and Medical section; CPT, Chemistry, Physics and Technology section; HUM, Human Sciences section.

Within the MPS sections (**Fig 21B-D**), the distribution of employment type was similar to the overall population.

Similarly, we observed a correlation between employment type and previous work experience. The correlation was stronger for MPS experience (**Fig S6**, p<0.001, Cramer’s V = 18.8%). In particular, postdocs with less than 3 years of MPS experience were more frequently on stipends (18.6%% – 70/ 376) than postdocs with more than 3 years of MPS experience (5.4% – 11/205). The correlation with the overall experience (**Fig S7**, p<0.01, Cramer’s V = 14.5%) was slightly weaker. Postdocs with more than 5 years of experience were less likely to hold a stipend (6.9% – 14/202) than postdocs with less than 5 years of experience (17.0% – 65/382).

#### 2. Fixed-term TVöD-based employment

Our analysis of fixed-term contract conditions focused on four topics: (i) funding source; (ii) salary distributions; (iii) wage group and level distributions; and, finally, (iv) recognition of experience for wage level assignment.

##### a) Funding source

First, the main funding source of contracts was the MPS with 278 out of 495 (56%) postdocs being paid directly by the MPS (**Fig 22**). The remaining postdocs reported that they received funding from other sources (30% – 149/495) or that they did not know where their funding came from (14% – 68/495). Funding source distributions significantly varied across MPS sections. The HUM section had the highest number of postdocs with MPS funding (81% – 71/88), while in the CPT and BM sections postdocs with MPS funding were roughly half of the respondents (CPT: 46% – 57/124; BM: 51% – 141/279). Further, 10% (9/88) of postdocs in the HUM section, 44% (54/124) of postdocs in the CPT section and 33% (91/279) of postdocs in the BM reported to be funded by other sources. Finally, a large portion of postdocs in the BM section did not know where their funding came from (17% – 47/279) while this number was smaller in the CPT (11% – 13/124) and HUM (9% – 8/88) sections.

**Figure 22.**
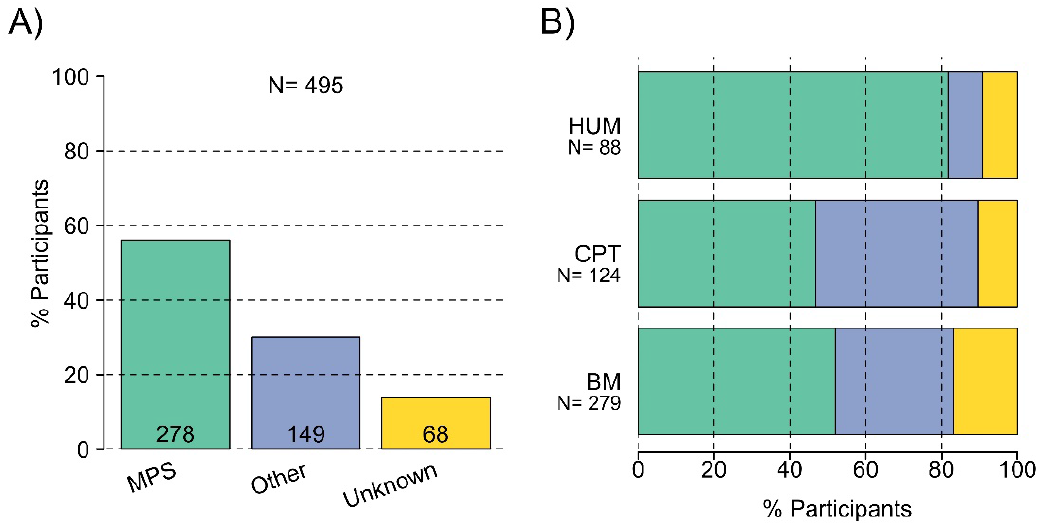
Contract funding sources. Distribution of funding sources for postdocs in the MPS overall **(A)** and in each MPS sections **(B)**. BM, Biology and Medical section; CPT, Chemistry, Physics and Technology section; HUM, Human Sciences section; MPS, Max Planck Society

##### b) Salary distributions

Salary distributions (**Fig 23A**) did not show any relevant bias with respect to MPS sections (**Fig 23B**), gender (**Fig 23D**), origin (**Fig 23E**) or PhD lab (**Fig 23F**). As expected, differences were found for age (**Fig 23C**) and overall experience (**Fig 23G**). In particular, older postdocs with more experience tended to earn more.

**Figure 23.**
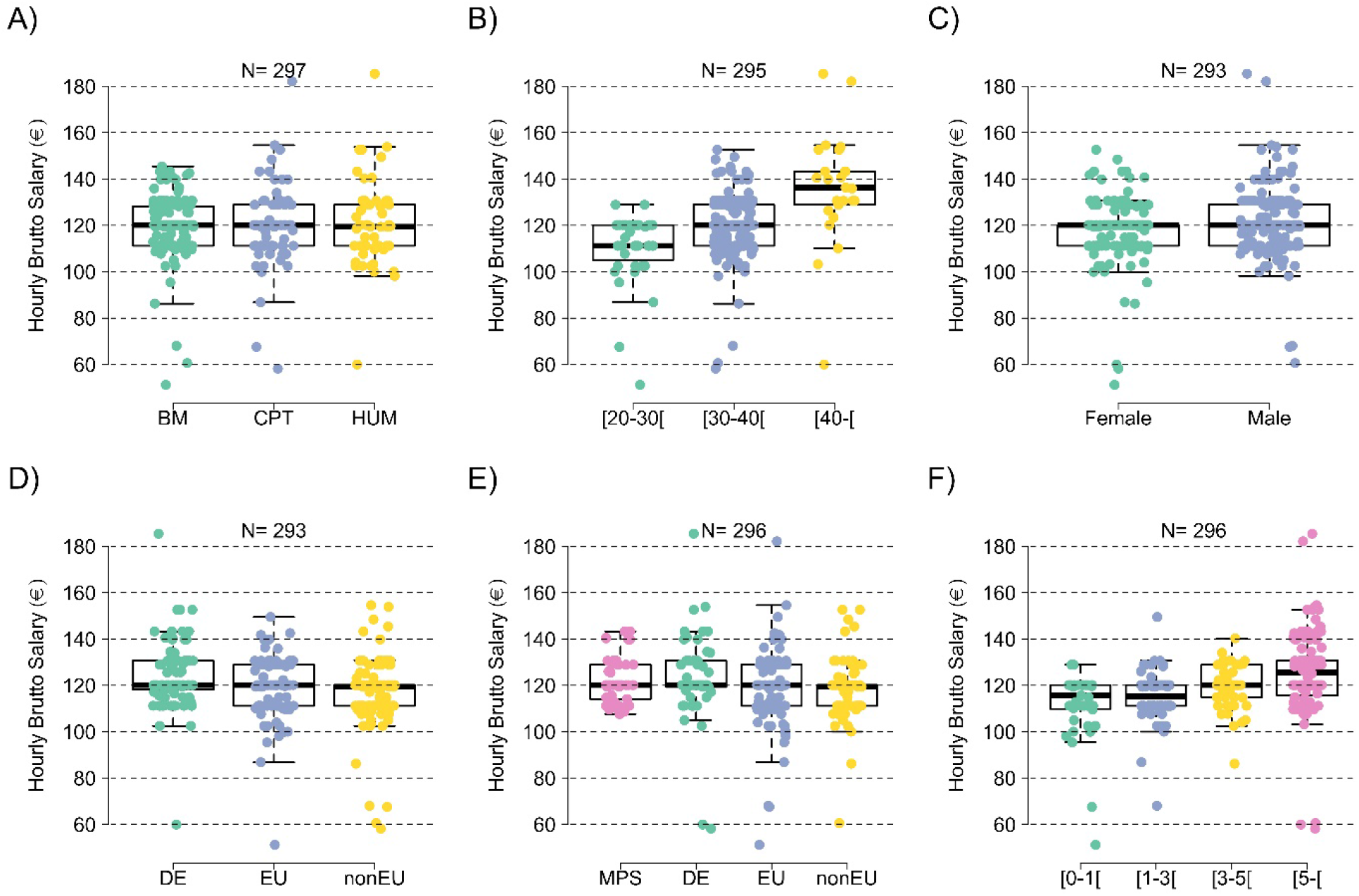
Salary distributions. Salary distributions of postdocs on contract by MPS sections **(A)**, age **(B),** gender **(C),** origin **(D),** PhD lab **(E)** and overall experience as postdocs **(F)**. Gross salary (in euros) per hour is displayed on the y-axis. Age is displayed in years and overall experience in years post PhD graduation. Origin corresponds to Germany (DE), a country in the European Union (EU) or countries outside of the European Union (nonEU). PhD lab corresponds to graduation from one MPI (MPS), from a German institution (DE), from a European institution (EU) or from a non-European institution (nonEU). Note, data points slightly change across these measurements, as not the same number of respondents answered all survey questions regarding personal salary. BM, Biology and Medical section; CPT, Chemistry, Physics and Technology section; HUM, Human Sciences section; MPS, Max Planck Society.

##### c) TVöD system

In Germany, scientists on a federal contract are employed under the *Tarifvertrag für den öffentlichen Dienst* (TVöD). This system is subdivided into wage groups (Entgeltgruppe - E) that are assigned based on the final degree and the job description. Having a master’s degree or equivalent, postdocs should be in E13. Each wage group is itself subdivided into wage levels (Stufe) that depend on the time spent in the previous wage level (**Tab 2**).

**Table 2.**
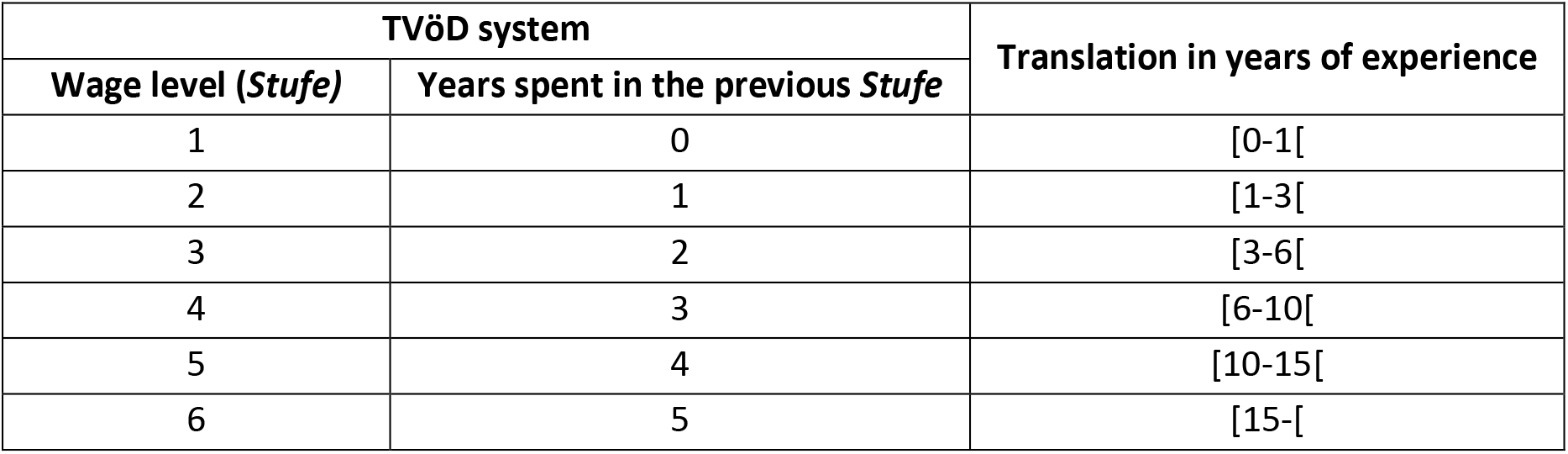
Wage levels (*Stufe)* assignment based on the TVöD system. The six possible wage levels and the number of relevant work experience in years according to the TVöD system^③^. The “Translation in years of experience” represents an interpretation of the TVöD regulations, assuming all years of research experience are taken into account.

##### d) Wage group (*Entgeltgruppe*) distributions

Analysis of the wage group (*Entgeltgruppe*) distributions among postdocs (**Fig 24**) revealed that overall (**Fig 24A**), most postdocs were in the wage group E13 (80% – 320/400). However, almost one fifth of postdocs (18% – 72/400) reported to be in the wage group E14 and a small minority in wage group E15 (4 postdocs), EU contracts (3 postdocs), WII (1 postdoc) or E12 (1 postdoc – non displayed on the graph). In particular, the biggest proportion of postdocs with an E14 contract were in the HUM section (20/67 – 30%), followed by the CPT (21% – 20/97) and BM (32/226 – 14%) sections (**Fig 24B**). Postdocs who reported to be on EU or WII contracts were not considered in the subsequent data analysis due to the size of the data set.

**Figure 24.**
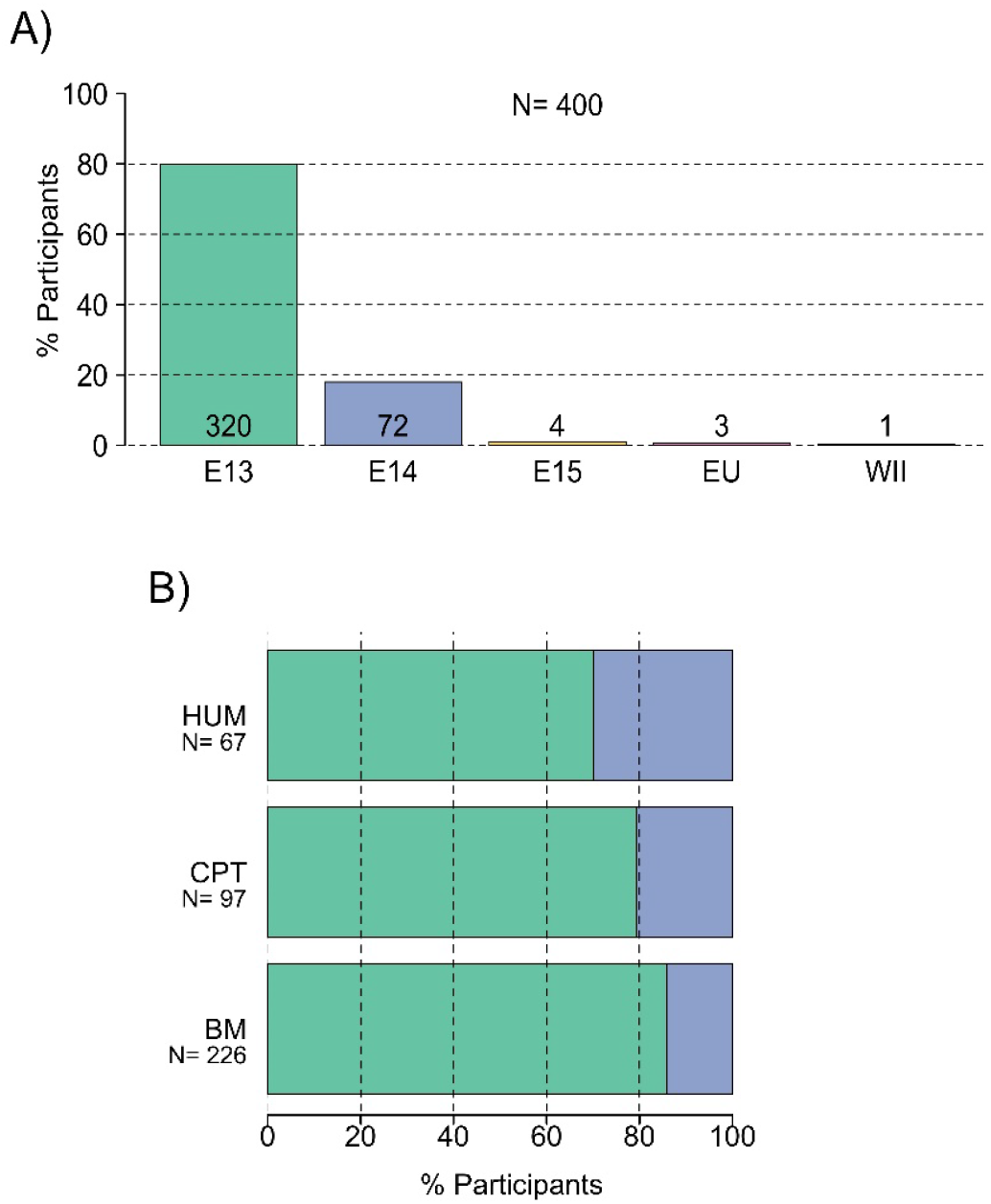
Wage group (*Entgeltgruppe*). Distribution of postdocs depending on their wage groups in the MPS overall **(A)** and in each MPS section **(B)**. Postdocs who reported to be on EU or WII contracts were not considered in the subsequent data analysis. BM, Biology and Medical section; CPT, Chemistry, Physics and Technology section; HUM, Human Sciences section.

Differences in the assignment of wage group (*Entgeltgruppe*) were found between postdocs of different origins. In particular, with respect to the fraction of postdocs on wage groups E13 and E14, a smaller fraction of German postdocs (75% – 96/128) were assigned to wage group E13 than international postdocs from either other EU countries (81% – 96/118) or from outside Europe (86% – 118/137) (**Fig 25**). Similar observations were visible in the sections.

**Figure 25.**
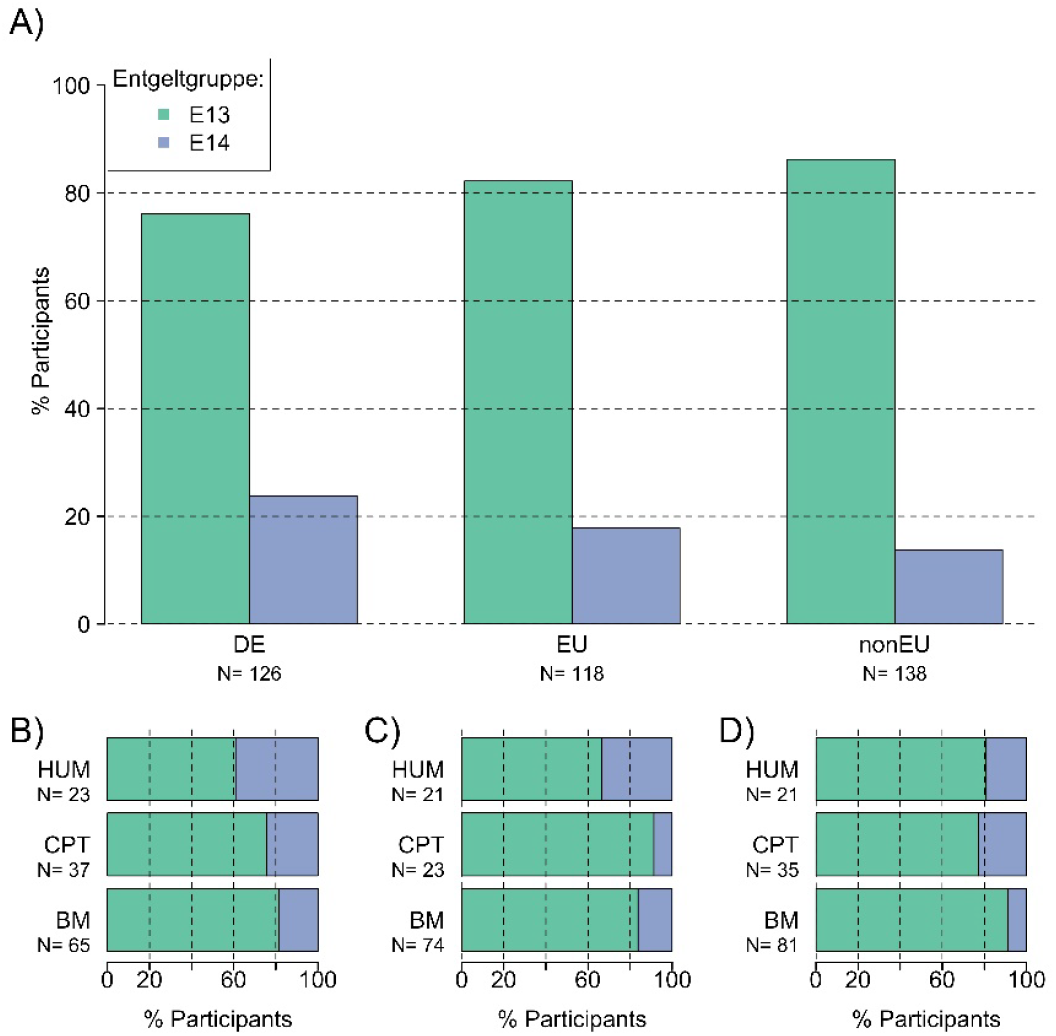
Wage groups (*Entgeltgruppe*) and origin. Fraction of German (DE), European Union (EU) and non-European Union (non-EU) postdocs in wage groups E13 and E14. BM, Biology and Medical section; CPT, Chemistry, Physics and Technology section; HUM, Human Sciences section.

As we scale this with experience, the tendency towards a greater proportion of German postdocs being bracketed into E14 wage group (*Entgeltgruppe)* continues (**Fig 26**). Among postdocs with up to 1 years of experience, German postdocs exceeded by 3% EU postdocs and by 4% non-EU postdocs. Further, among postdocs with between 1 and 3 years of experience, German postdocs exceeded by 10% EU postdocs and by 2% non-EU postdocs (**Fig 26** top panel). Finally, among postdocs with between 3 to 5 years of experience, German postdocs exceeded by 12% all international postdocs and among postdocs with more than 5 years of experience, German postdocs exceeded by 3% EU postdocs and by 20% non-EU postdocs (**Fig 26** bottom panel). Similar distributions were observed in the wage group E15, despite the small fraction of postdocs in this wage group. In particular, the fraction of German postdocs in E15 (4%) was higher than the fraction of postdocs from both EU (~2%) and outside Europe (0%). However, these differences in wage groups between German and international postdocs did not reach significance (Fisher exact test; p>0.05).

**Figure 26.**
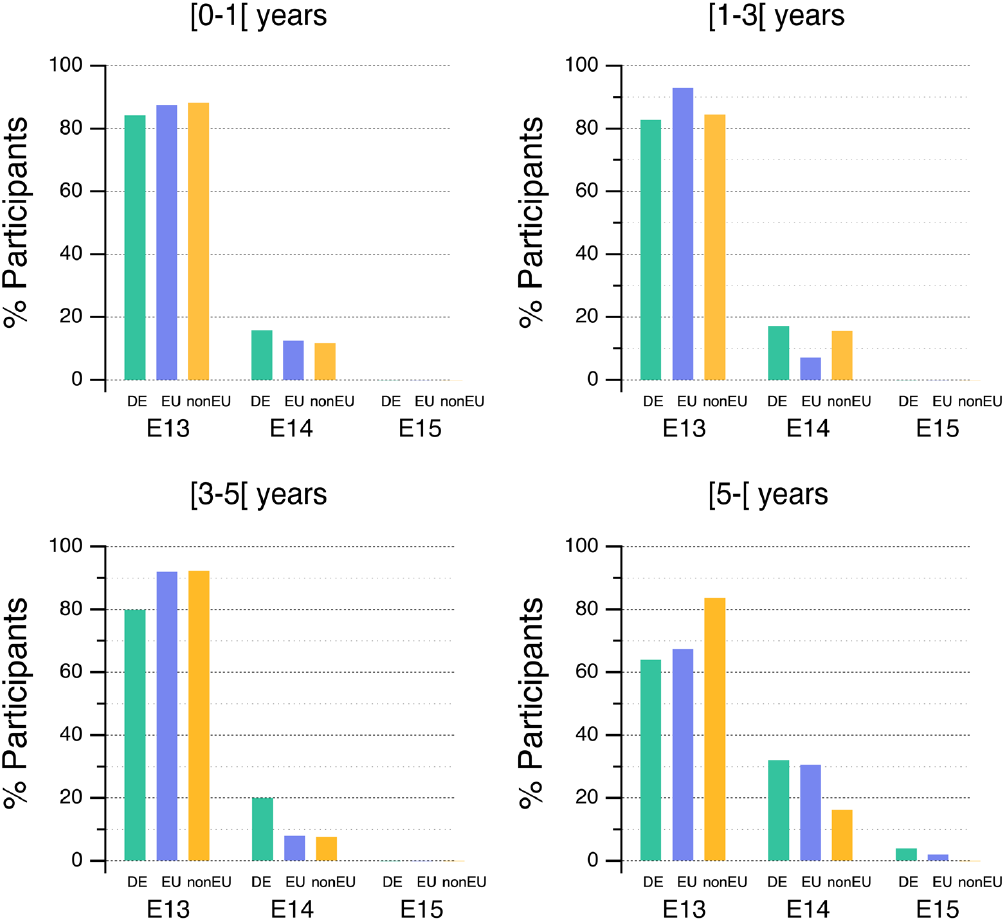
Origin, overall experience and wage groups (*Entgeltgruppe*). Fraction of German (DE, green), European Union (EU, blue) and non-European Union (non-EU, orange) postdocs in wage groups E13, E14 and E15 across years of postdoctoral experience.

##### e) Wage level (*Stufe*) distributions

With respect to the wage level (*Stufe*) of TVöD contracts (**Fig 27**), wage levels followed a normal distribution with a peak at level 3 (**Fig 27A**) with 44% (160/360) of postdocs being in this level. The other two most common wage levels were wage level 2 (26% – 95/360) and 4 (19% – 68/360). A small minority of postdocs was on wage level 1 (0.05% – 19/360) and 5 (0.05% – 18/360). Slightly different wage level distributions were observed across MPS sections (**Fig 27B**). In particular, the majority of postdocs from the BM section were on wage level 3 (51% – 107/208), followed by postdocs from the CPT (36% – 32/89) and the HUM (31% – 19/61) sections. Most postdocs in the HUM (level 2: 38 % – 23/61; level 1: 8% – 5/61) and the CPT (level 2: 33% – 29/89; level 1: 8% – 7/89) sections were on wage levels lower than 3. In comparison, the BM section had the least number of postdocs on wage levels 1 and 2 (level 2: 21% – 43/208; level 1: 3% – 7/208).

**Figure 27.**
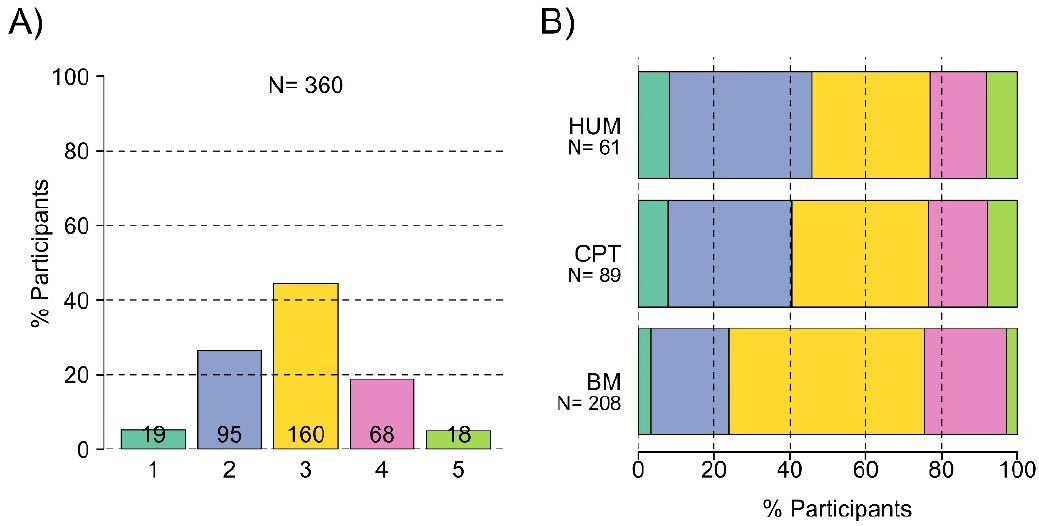
Wage level (*Stufe*). Distribution of postdocs depending on their wage levels in the MPS overall **(A)** and in each MPS section **(B)**. Answers from postdocs in wage groups E13 and E14 were used. BM, Biology and Medical section; CPT, Chemistry, Physics and Technology section; HUM, Human Sciences section.

The overall postdoctoral experience was compared to the wage levels assigned by the TVöD system (**Tab 2**).

Postdoctoral experience and wage level (*Stufe*) assignment do largely correlate with each other (**Fig 28**). However, there were also cases where postdoctoral experience and wage level assignment were not correlated For instance, a non-negligible proportion of postdocs on wage levels 1 (15.8% – 3/19) and 2 (34.7% – 33/95) reported to have more than 3 years of overall postdoctoral experience

**Figure 28.**
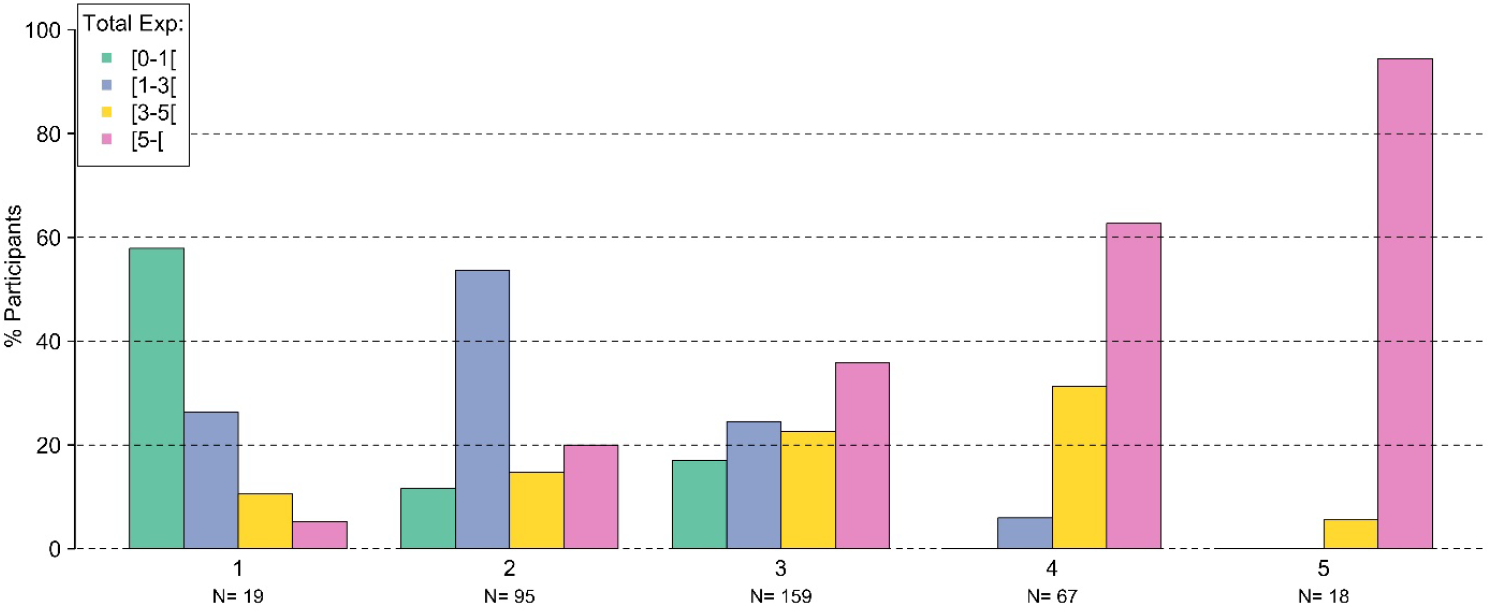
Overall experience and wage level *(Stufe*). Overall postdoctoral experience (years) according to wage levels. Answers from postdocs in wage groups E13 and E14 were used.

The tendency of higher wage levels (*Stufe* i.e. 1-6) being more frequently assigned to German relative to international postdocs was observed within each wage groups (i.e., E13, E14 or E15). In particular, the fraction of international postdocs found in lower wage levels was higher than the fraction of German postdocs (**Fig 29** and **Fig S8**). Among postdocs with less than one year of experience, 56% of EU postdocs and 50% of non-EU postdocs were assigned to the lowest wage levels (i.e., 1 and 2). On the contrary, only 32% of German postdocs were assigned to the lowest wage levels (**Fig 29**, top panel).

**Figure 29.**
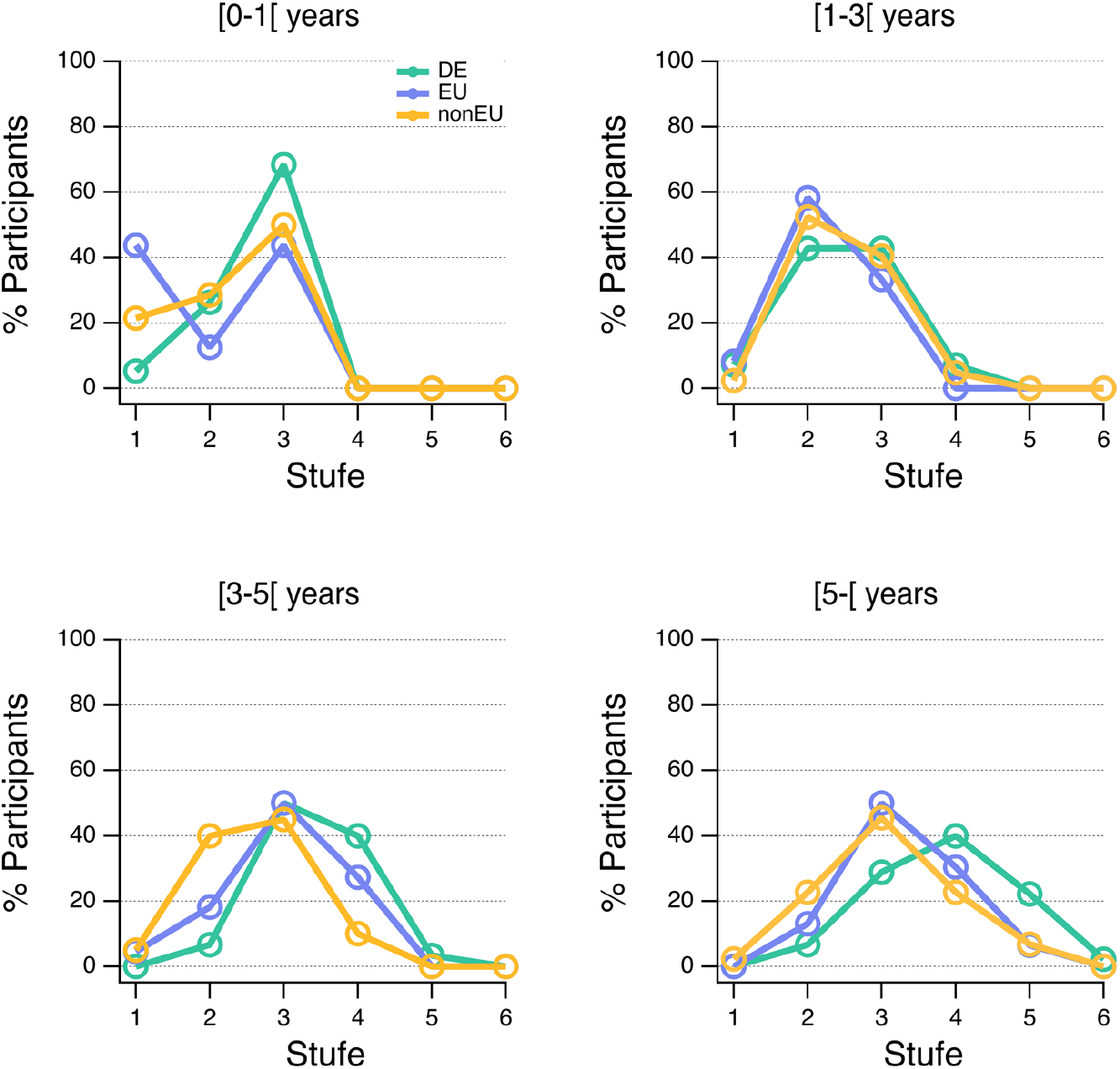
Origin, overall experience and wage levels (*Stufe*). Fraction of German (DE, green), European Union (EU, blue) and non-European Union (non-EU, orange) postdocs in wage levels 1 to 6 across years of postdoctoral experience. Answers from postdocs in wage groups E13, E14 and E15 were used.

Like for the wage groups, this tendency in wage level (*Stufe*) assignment between German and international postdocs increased with increasing years of postdoctoral experience. Specifically, this difference was at the highest among postdocs with 3-5 and more than 5 years of experience (**Fig 29**, bottom panel).

> *“I was told that only German experience counts for the salary category decision. Therefore I had to start for the lowest salary in E13, although I was hired based on my previous experience, knowledge and skills acquired in non-EU institute.”*
>
> — Participant’s comment

In the two last groups (3-5 and more than 5 years of experience), a shift to higher wage levels can be appreciated in the distribution of EU *vs.* non-EU researchers and for German *vs.* EU researchers. In particular, the fraction of international postdocs at wage levels 1 and 2 was 23% for EU and 45% for non-EU but only 7% for German postdocs with 3 to 5 years of experience. Conversely, only 27% of EU postdocs and 10% of non-EU postdocs, but up to 43% of German postdocs were found at wage levels higher than 3. Similar patterns were observed for postdocs with more than 5 years of experience. Specifically, the fraction of international postdocs at wage levels 1 and 2 was 25% for EU and 13% for non-EU but only 7% for German postdocs with more than 5 years of experience. On the contrary, only 37% of EU postdocs and 30% of non-EU postdocs but up to 64% of German postdocs were found at wage levels higher than 3. The proportion of postdocs in Stufe 1-2 *vs* 3-6 was statistically different comparing German and international postdocs in the experience category 3 to 5 years (Fisher exact test, p=0.009), and did not reach significance for other experience groups (Fisher exact test, P >0.05).

Finally, we examined in which wage levels (*Stufe*) postdocs should been assigned to if either their postdoctoral experience or their ‘PhD **and** postdoctoral’ experiences were taken into consideration (**Tab 3**). The obtained analysis displayed similar trends, thus only wage levels assignment when ‘PhD **and** postdoctoral’ experiences were recognized are presented (**Fig 30**). Wage levels (*Stufe*) assignment when overall postdoctoral experience was recognized can be found in supplementary **Fig S9**. To determine the ‘PhD **and** postdoctoral’ experience, we set the duration of the PhD experience to 3 years for all postdocs since it is the minimal duration. Accordingly, all new postdocs should receive a contract with wage level 3 after completion of their PhD (**Tab 3**).

**Table 3.** Expected wage levels (*Stufe*) according to postdoctoral experience, with and without recognition of PhD experience. The table shows in which wage levels postdoc should be placed in according to their postdoctoral experience (years), depending on whether the PhD experience is recognized as such. These expectations stem from our interpretation of the TVöD system presented in Tab 2. A PhD duration of 3 years was assumed.

| Postdoctoral experience (years) | Expected wage level (*Stufe*) with and without PhD experience recognition |  |
| --- | --- | --- |
|  | Without recognition | With recognition |
| [0-1[ | 1 | 3 |
| [1-3[ | 2 | 3/4 |
| [3-5[ | 3 | 4 |
| [5-[ | 3+ | 4+ |

**Figure 30.**
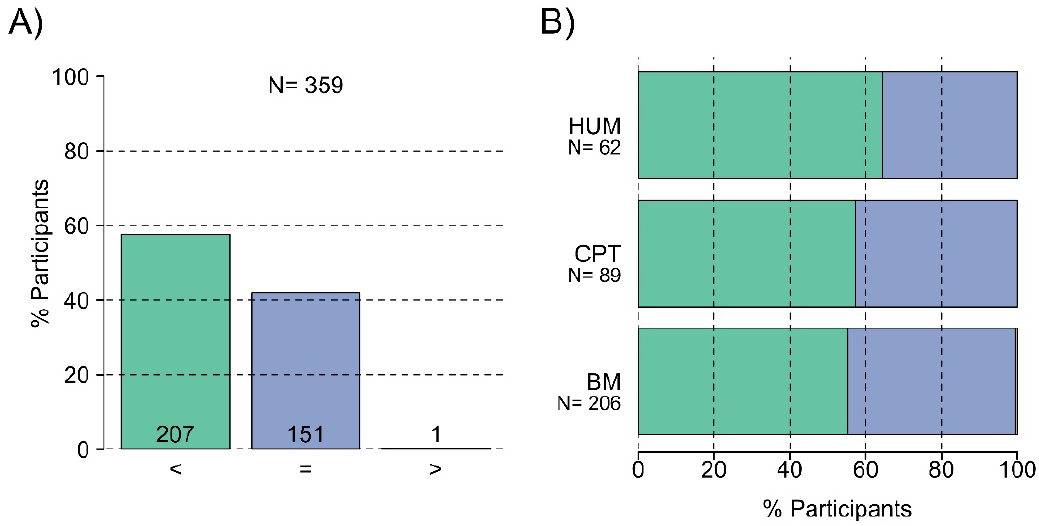
Appropriateness of wage level (*Stufe*) assignment. Fraction of postdocs placed in the appropriate wage level after recognition of a three-year doctoral experience as a function of postdoctoral experience (in years) in the MPS overall **(A)** and in each MPS section **(B)**. Postdocs found either in lower (green, <), appropriate (blue, =) or higher (yellow, >) wage level. BM, Biology and Medical section; CPT, Chemistry, Physics and Technology section; HUM, Human Sciences section.

Results show that 58% of postdocs (207/359) had a wage level (*Stufe*) lower than their ‘PhD **and** postdoctoral’ experience (**Fig 30A**). Results across MPS sections (**Fig 30B**) show that postdocs were less likely to have their PhD experience recognized in the HUM section (40/62, 65%) as opposed to the CPT (57% – 51/89) and BM (55% – 114/206) sections.

> *“Should PhD count as an experience in deciding the Stufe for our salary? It is very annoying if certain labs within MPI count them and some labs don’t. It will be good to have a consistent policy for counting PhD as work experience.”*
>
> — *Participant’s comment*

Postdocs whose PhD experience was not recognized were mainly between 20 and 30 years (60% – 28/47) or between 30 and 40 years (62% – 172/279). On the contrary, only 10% of postdocs older than 40 years (3/62) were placed in a lower wage level (*Stufe*) than their overall experience (**Fig 31A**). This overall pattern was further reflected in each of the MPS sections (**Fig 31B**).

**Figure 31.**
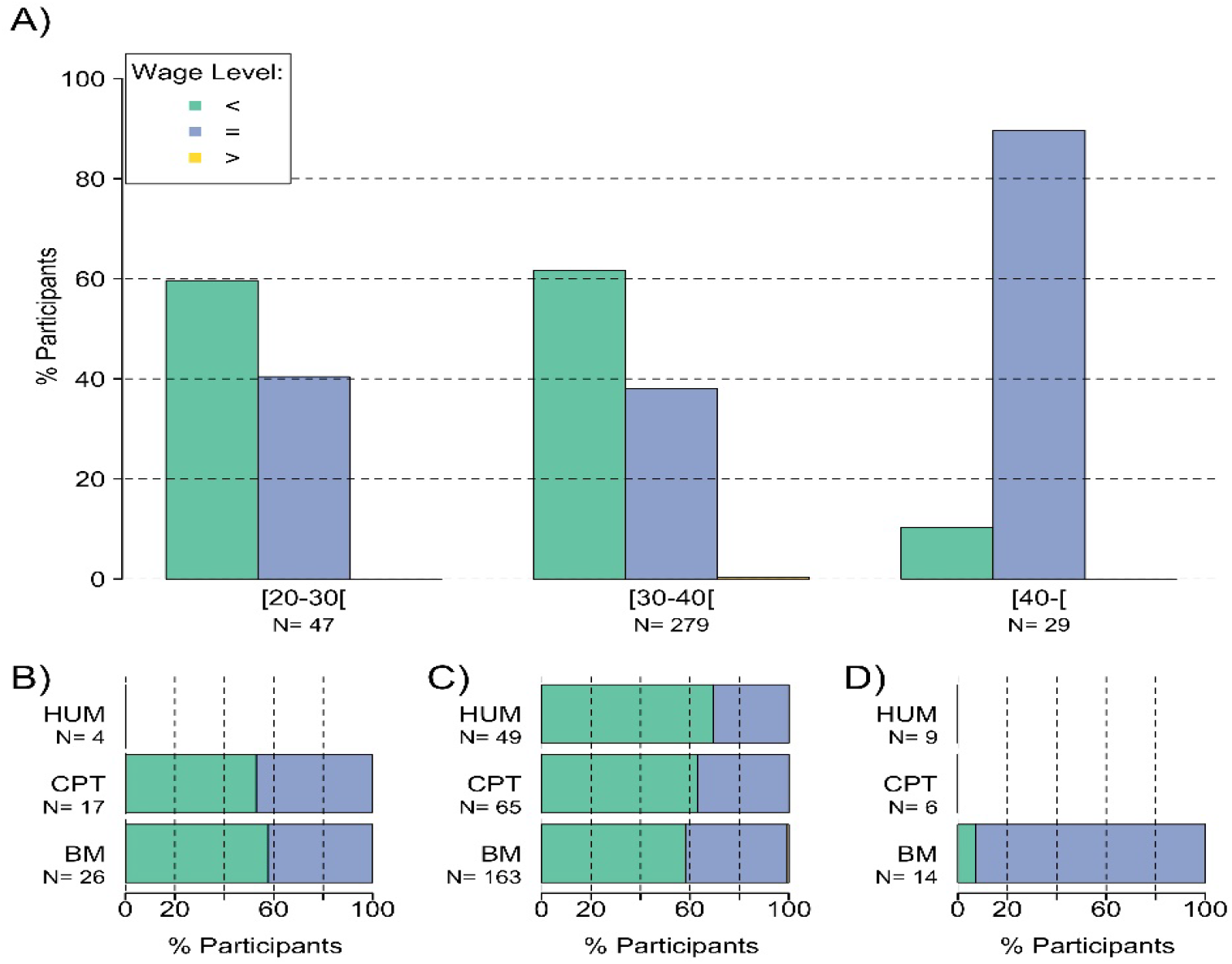
Appropriateness of wage level (*Stufe*) assignment and age. Fraction of postdocs placed in the appropriate wage level after recognition of a three-year PhD experience as a function of age (years) in the MPS **(A)** and in each MPS section **(B-D).** Postdocs found either in lower (green, <), appropriate (blue, =) or higher (yellow, >) wage level. BM, Biology and Medical section; CPT, Chemistry, Physics and Technology section; HUM, Human Sciences section.

We further confirmed the result from **Fig 29** across years of experience. In particular, taking into consideration the overall postdoctoral experience, doctoral experience was not recognized for a large fraction of postdocs (**Fig 32**). Similar results were observed also when considering only MPS postdoctoral experience (**Fig S10**).

**Figure 32.**
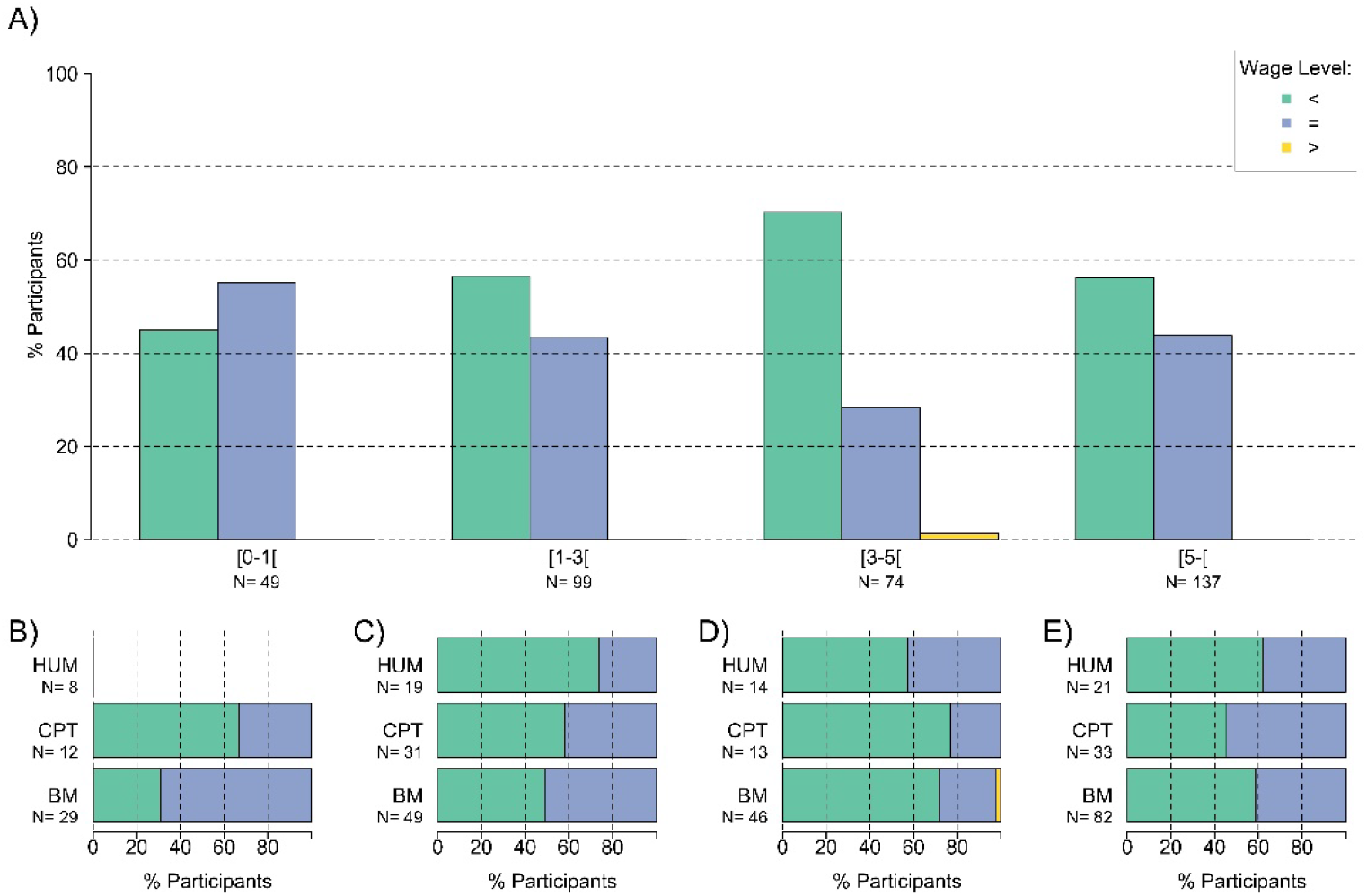
Appropriateness of wage level (*Stufe*) assignment and overall experience. Fraction of postdocs placed in the appropriate wage level after recognition of a three-year PhD experience as a function of the overall postdoctoral experience (years) in the MPS **(A)** and in each MPS section **(B-E).** Postdocs found either in lower (green, <), appropriate (blue, =) or higher (yellow, >) wage level. BM, Biology and Medical section; CPT, Chemistry, Physics and Technology section; HUM, Human Sciences section.

The differences in wage levels (*Stufe*) assignment between German, EU and non-EU postdocs (see **Fig 29**) were also visible when taking into account PhD experience. German postdocs were more likely to be assigned to the adequate wage level based on their PhD experience and were hence better paid than EU and non-EU postdoctoral researchers who tend to be placed in lower wage levels than what their years of experience would require (**Fig 33**).

**Figure 33.**
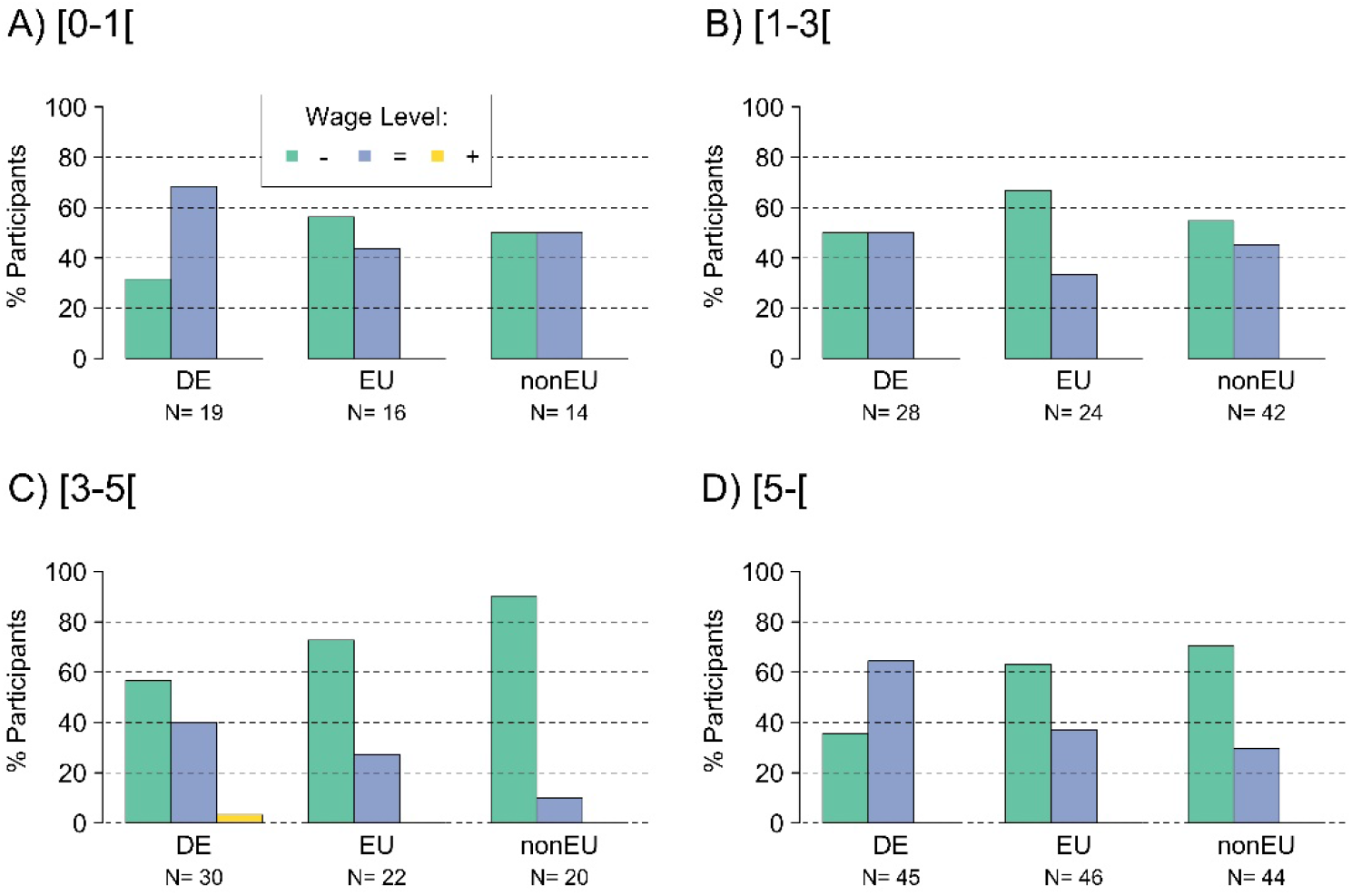
Appropriateness of wage level (*Stufe*) assignment, overall experience and origins. Fraction of postdocs placed in the appropriate wage level after recognition of a three-year PhD experience as a function of their origin, separated by overall postdoctoral experience (years): [0-1[**(A)**, [1-3[ **(B)**, [3-5[ **(C)** and [5-[ **(D)**. Postdocs found either in lower (green, -), appropriate (blue, =) or higher (yellow, +) wage level. DE, German; EU, European Union; nonEU, non-European Union.

A difference between genders was only observed for postdoctoral researchers with 1-3 years of postdoctoral experience (**Fig 34**). In particular, male postdocs with 1 to 3 years of experience tended to be placed in a lower wage level (*Stufe*) when taking into consideration their PhD experience (70% – 32/46 of male postdocs with 1-3 years of experience) as compared to female postdocs (44% – 22/50 of female postdocs).

**Figure 34.**
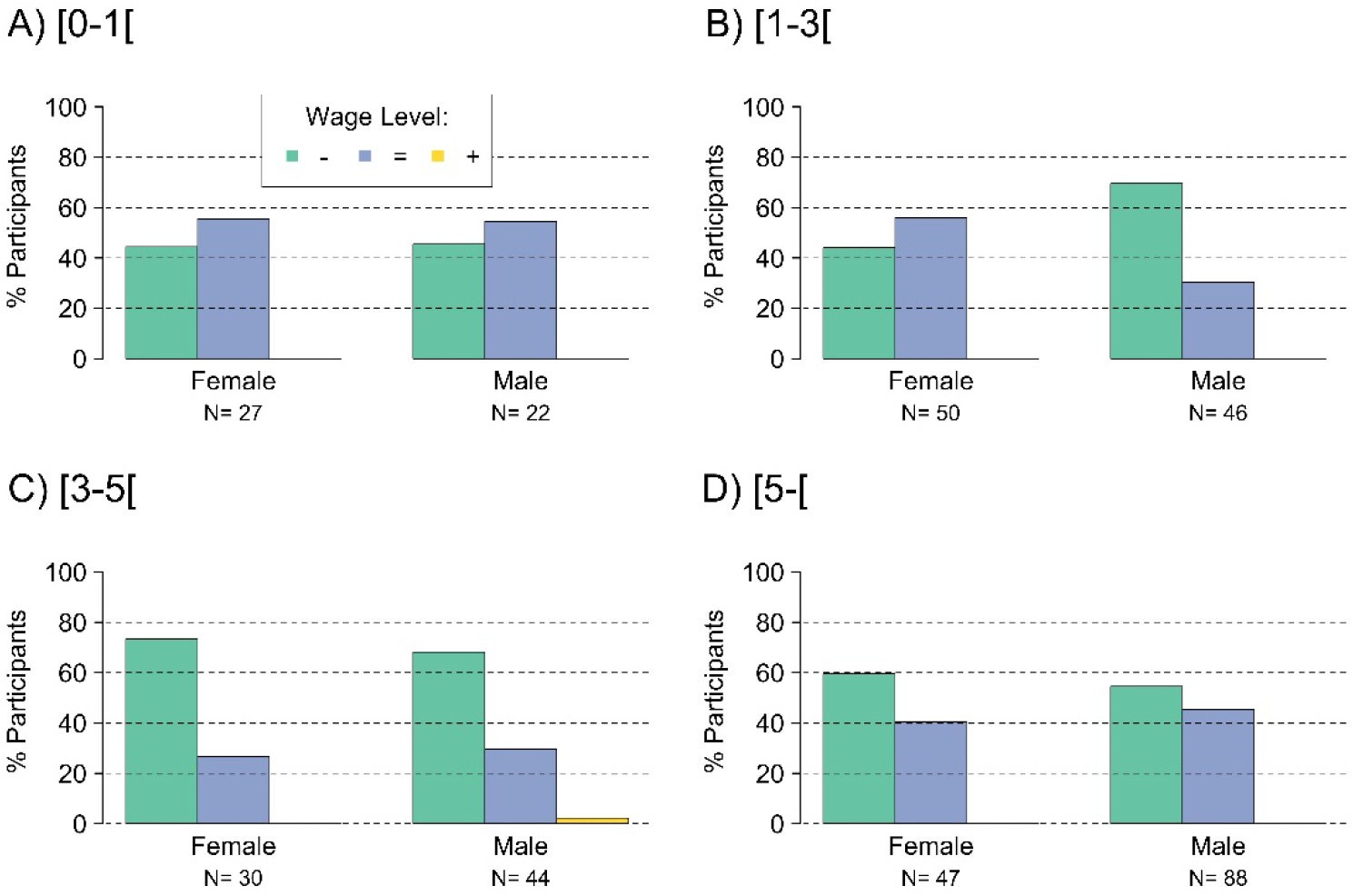
Appropriateness of wage level (*Stufe*) assignment, overall experience and gender. Fraction of postdocs placed in the appropriate wage level after recognition of a three-year PhD experience as a function of their origin, separated by overall postdoctoral experience (years): [0-1[(A), [1-3[(B), [3-5[(C) and [5-[(D). Male and female postdocs found either in lower (green, -), appropriate (blue, =) or higher (yellow, +) wage level.

##### f) Experience recognition for wage level assignment

Finally, to test the transparency of the procedures that lead to the determination of wage levels for postdocs’ salary, we asked postdocs to report: (i) whether they knew if their experience was considered for wage level (*Stufe*) assignment; (ii) whether they knew, before starting the postdoctoral position, which wage level they would receive; and (iii) the rationale or criteria for receiving it (**Fig 35**).

**Figure 35.**
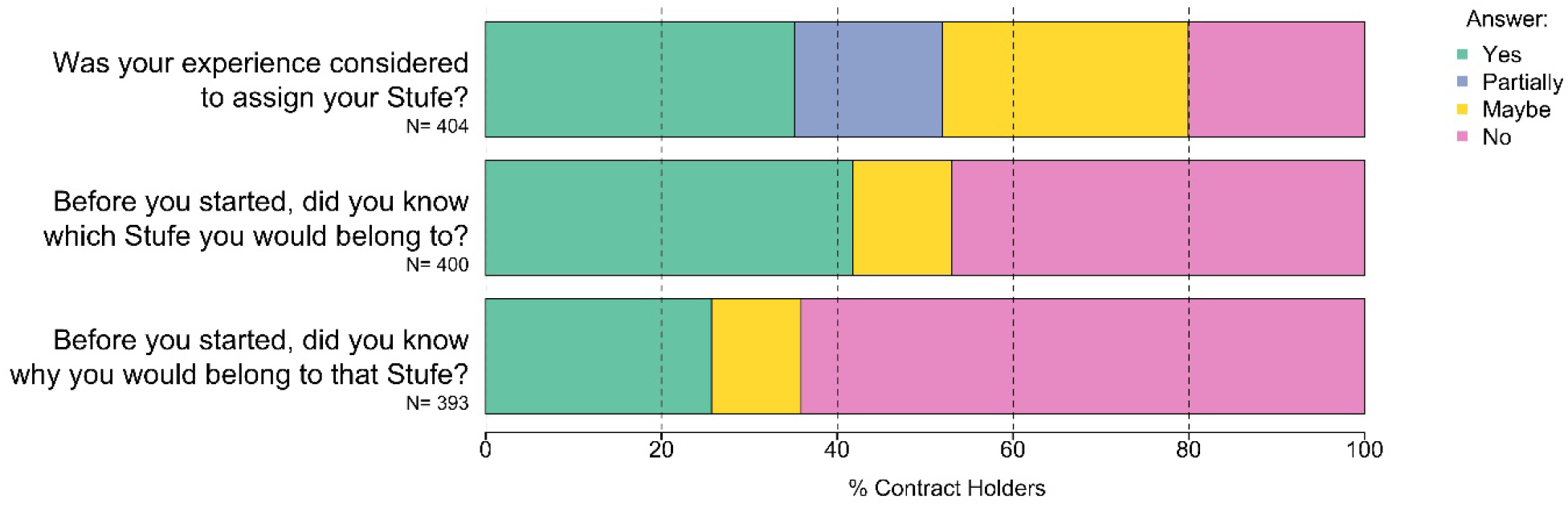
Transparency about wage level (*Stufe*) assignment. To test transparency of contractual conditions for postdocs’ employment, postdocs were asked to report whether they knew their experience was considered for wage level assignment (**top**); whether they knew the wage level they would receive (**middle**); and the reasons why they would receive it (**bottom**).

Concerning the recognition of experience in the assignment of the wage level, 34.6% (140/404) of the postdocs were certain that their experience was taken into account while 28% (113/404) were not certain of it. A lack of experience recognition was reported by 36.9% (149/404) of the postdocs. Their wage level was either not recognizing their experience (20% – 81/404) or only partially recognizing it (17% - 68/404). Importantly, the majority of postdocs did not know which wage level they would belong to before starting their postdoctoral position (42% – 167/400) or were unsure about it (11% – 45/400). Furthermore, the majority of postdocs did not know why they were assigned to the wage level they received (64% – 252/393). Only around one fourth of postdocs (26% – 101/393) knew the rationale behind their wage level assignment.

> “I only knew my salary once I got my first payslip. No one (also not HR, nor the PI) had an idea how much I would get. Pretty annoying when trying to find a flat, and you don’t know how much you will earn.”
>
> — Participant’s comment

> “That I would be E13 was clear from the beginning, but which level was not specified until I got the contract.”
>
> — Participant’s comment

#### 3. Stipends

##### a) Type of stipend

In order to evaluate the working conditions linked to being employed on a stipend, we separated stipend holders according to the type of stipend that they receive:

**- MPS scholarship**: a stipend awarded directly by the MPS, usually provided by the supervisor of the postdoc. Here, the MPS determines the employment conditions;
**- Third-party fellowships**: a stipend usually awarded by a funding agency from which the postdocs themselves receive a grant. For this stipend type, the MPS has little control over the employment conditions.

Among stipend-holders, 47.6% (39/82) received an MPS scholarship and 46.3% (38/82) received a fellowship from a third-party funding agency, while 6.1% (5/82) could not be attributed to either scholarship or fellowship due to the lack of additional information. Thus, these uncategorized stipends were excluded from further analysis, leading to an overall proportion of scholarship holders as compared to fellowship holders of about 50% (**Fig 36A**).

**Figure 36.**
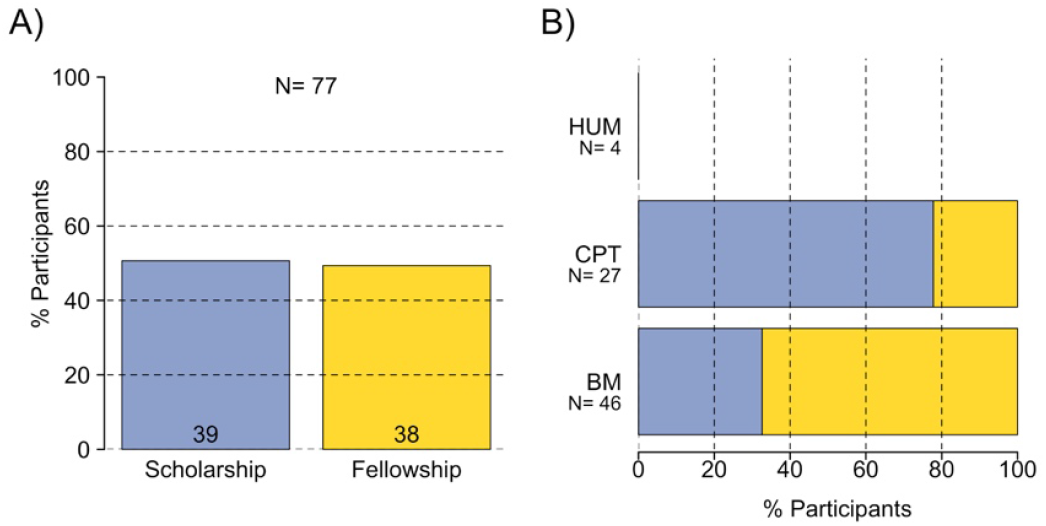
Type of stipend. Distribution of MPS scholarship and third-party fellowships in the MPS overall **(A)** and in each MPS section **(B)**. Note, statistics with less than 10 data points are not shown on graphs.BM, Biology and Medical section; CPT, Chemistry, Physics and Technology section; HUM, Human Sciences section.

Remarkably, different stipend types were preferred in different MPS sections (**Fig 36B**, p<0.001, Cramer’s V = 44%): 67.4% (31/46) of stipends were fellowships in the BM section (representing 81.6% – 31/38 of all fellowships), while 77.7% (21/27) were scholarships in the CPT section (representing 53.8% – 21/39 of all scholarships). The HUM section had a similar distribution as the CPT section (3/4 = 75% of scholarship), but the low sample size made it difficult to interpret.

Given the strong correlation between section and stipend type, and the low sample size of stipend holders in general, further correlations with other parameters will not be crossed with MPS section to avoid downsizing sample size beyond the interpretable point.

When comparing stipend type with demographics and experience parameters (**Fig 37**), we observed no significant correlations with age (**Fig 37A**, p>0.5), origin (**Fig 37C**, p>0.1), or PhD lab (**Fig 37D**, p>0.5).

**Figure 37.**
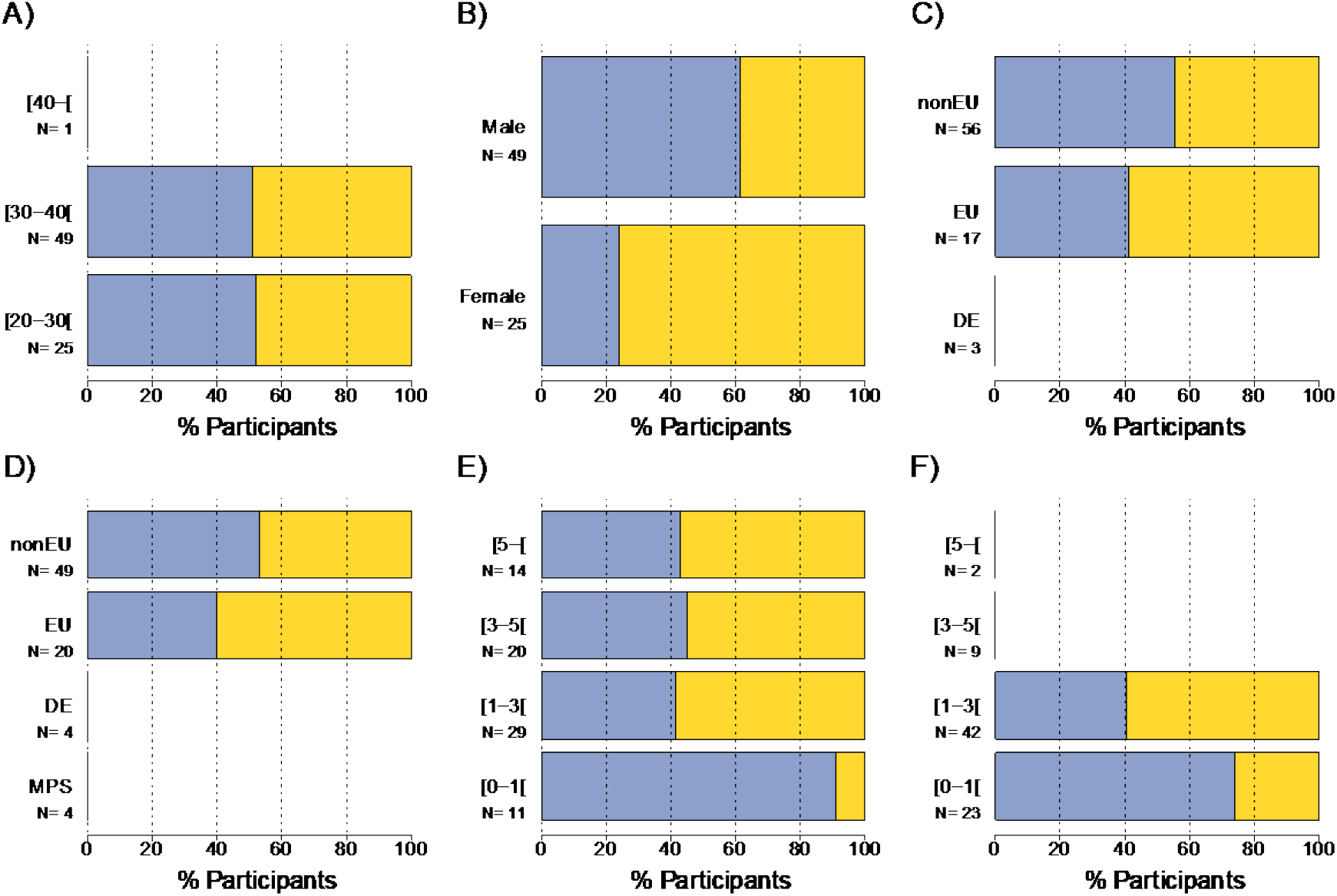
Demographics and experience information of stipend holders. Distributions of MPS scholarship (blue) and third-party fellowships (yellow) given based on age **(A)**, gender **(B)**, origin **(C)**, PhD lab **(D)**, overall experience **(E)** and MPS experience **(F)**. Age is displayed in years and both experiences in years post PhD graduation. Origin corresponds to Germany (DE), a country in the European Union (EU) or countries outside of the European Union (nonEU). PhD lab corresponds to graduation from one MPI (MPS), from a German institution (DE), from a European institution (EU) or from a non-European institution (nonEU). Note, statistics with less than 10 data points are not shown on graphs.

Indeed, both age groups with sufficient sample size had a proportion of scholarship to fellowship close to 50% (**Fig 37A**): 52.0% (13/25) for the age group between 20 and 30 years old and 51.0% (25/49) for the age group between 30 and 40 years old. Non-Europeans (**Fig 37C**) or postdocs that graduated outside the EU (**Fig 37D**) received slightly more scholarships (55.4% – 31/56 and 53.1% – 26/49 respectively) than Europeans or postdocs that graduated within the EU (41.2% – 7/17 and 40.0% – 8/20 respectively). German postdocs on a stipend all received a fellowship, but the sample size was very low. Thus, this number should be interpreted with caution. Similarly, postdocs that graduated within Germany (MPS included) and received a stipend were few, with 62.5% (5/8) of them receiving a scholarship.

However, there were strong correlations with gender (**Fig 37B**, p<0.01, Cramer’s V = 35.2%), overall experience (**Fig 37E**, p<0.05, Cramer’s V = 34.3%) and MPS experience (**Fig 37F**, p<0.05, Cramer’s V = 34.2%).

Remarkably, male stipend holders were mostly employed on an MPS scholarship (61.2% – 30/49), and represented 83.3% of all scholarship holders (30/36), while female stipend holders were mostly receiving fellowships (19/25 = 76%) and represented 50% of all fellowship holders (19/38). Interestingly, less experienced postdocs mainly received scholarships (90.9% – 10/11 for postdocs with less than one year of overall experience, and 73.9% – 17/23 of postdocs with less than one year of experience within the MPS). More experienced postdocs received more fellowships (55.6% – 36/63 for postdocs with more than one year of overall experience, and 60.4 % – 32/53 for postdocs with more than one year of experience within the MPS).

> *”The difference between having a contract and having a stipend is quite impactful. It also prima facie discriminates against male post-docs of non-EU origin.”*
>
> — Participant’s comment

##### b) Monthly allowance

The monthly allowance of a stipend depended highly on the type of stipend (p<0.001, Eta = 53.0%), with a median at 2100 euros for scholarships and at 2650 euros for fellowships (**Fig 38**). Additionally, fellowship allowances varied more than scholarship allowances. Scholarship allowances ranged from 2000 euros to 2323 euros, while fellowship allowances ranged from 1900 euros to 5000 euros.

**Figure 38.**
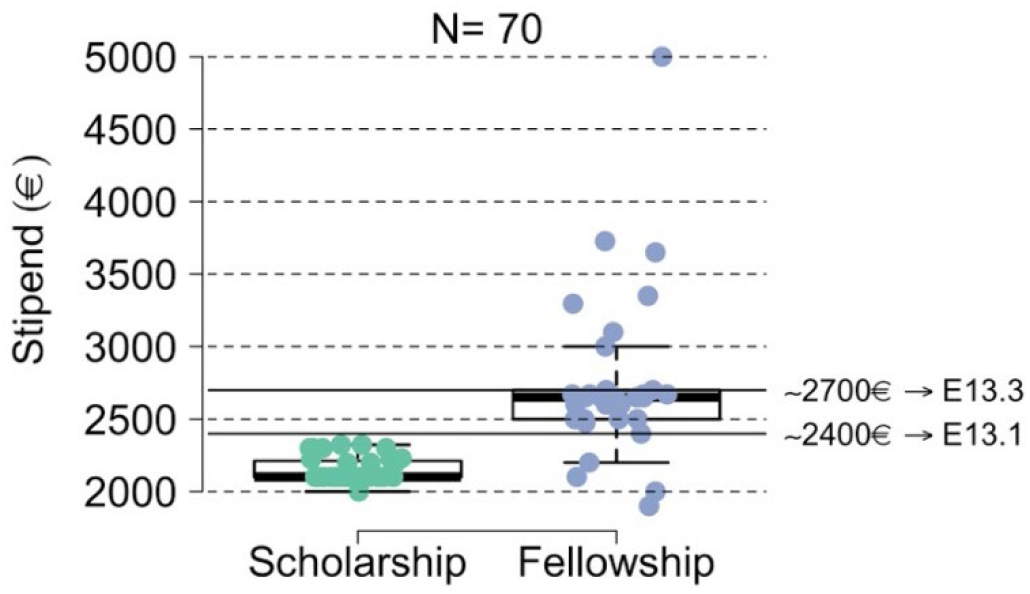
Monthly allowances of stipends. Monthly allowance according to stipend type is displayed in Euros. As reference are shown the monthly net salary of postdoc employed full-time on a TVöD contract in E13.1 and E13.3, with tax level 1 in 2019 with, respectively, 41.2% and 43% withdrawn from the gross salary.

> *“My stipend (fellowship) started after I had a contract. Though the fellowship is prestigious, I believe I am taking a paycut when considering all costs.”*
>
> — Participant’s comment

> *“If the MPS can’t even support the ‘fair salary’ system for postdocs, I doubt there is any intention to help the postdocs.”*
>
> — Participant’s comment

With respect to scholarships, we compared the monthly allowances with demographics and working experience (**Fig 39**). We observed that the median scholarship allowance did not differ with postdocs’ age (**Fig 39A**, p>0.5), gender (**Fig 39B**, p>0.8), or origin (**Fig 39C**, p>0.8). However, a trend was visible when considering the PhD lab (p=0.19, Eta = 5.7%). In particular, the median allowance of postdocs that had graduated in the EU was 2261.5 euros, while it was 2198 euros for postdocs that graduated in Germany and 2100 euros for those who graduated in the MPS or in a non-EU country (**Fig 39D**).

**Figure 39.**
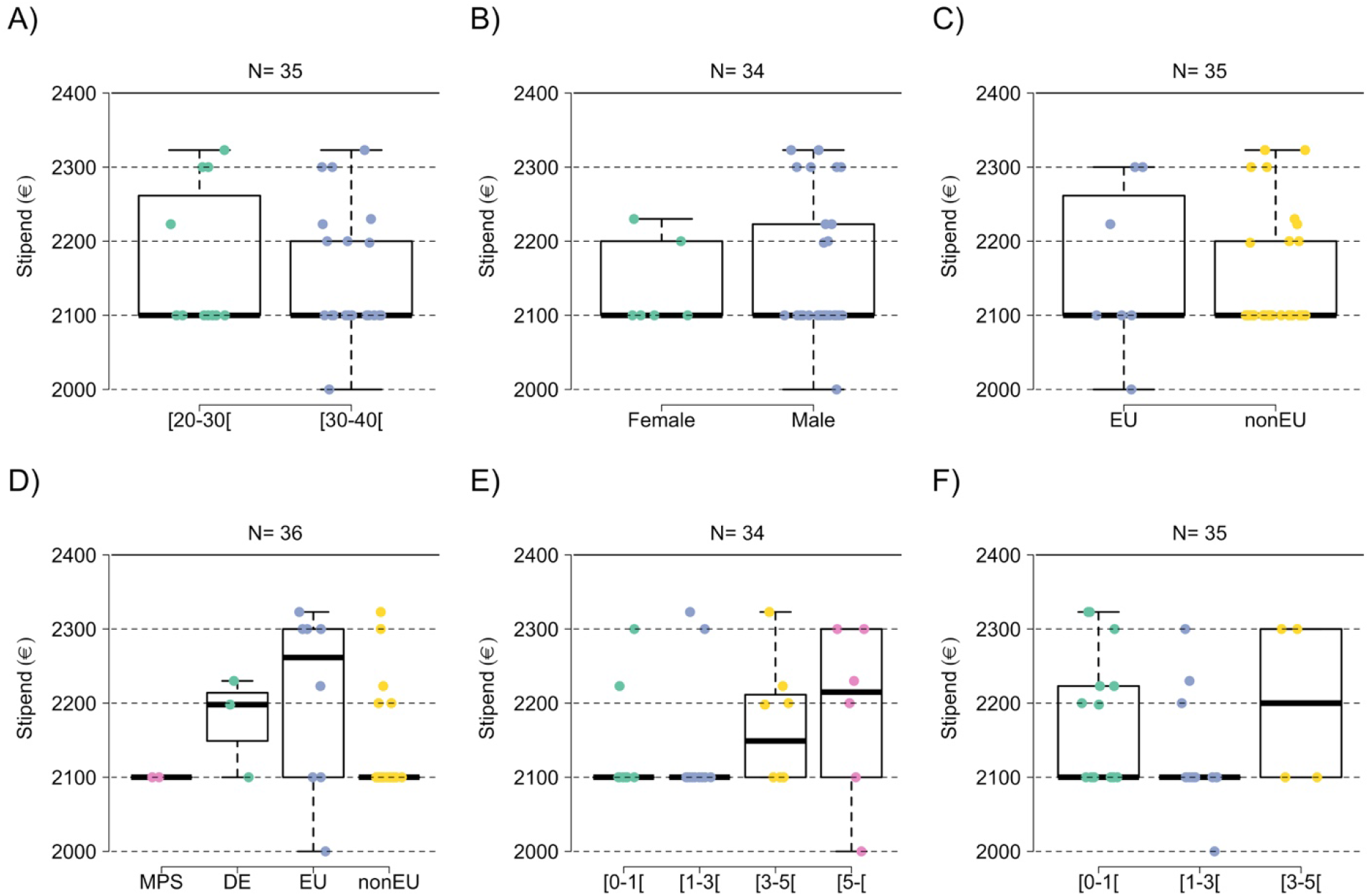
Monthly MPS scholarship allowances. Distributions given based on age **(A)**, gender **(B)**, origin **(C)**, PhD lab **(D)**, overall experience **(E)** and MPS experience **(F).** Age is displayed in years and both experiences in years post PhD graduation. Origin corresponds to Germany (DE – no postdoc), a country in the European Union (EU) or countries outside of the European Union (nonEU). PhD lab corresponds to graduation from one MPI (MPS), from a German institution (DE), from a European institution (EU) or from a non-European institution (nonEU). Monthly allowance is displayed in Euros.

Interestingly, the median scholarship allowance seemed to increase with the overall experience of postdocs, ranging from 2100 euros for those with less than three years of postdoctoral experience, to 2215 euros for postdocs with more than five years of experience (**Fig 39E**), although this was not significant (p>0.5), probably due to the low sample size. A similar trend was visible with MPS experience (**Fig 39F**, p=0.19, Eta = 3.9%), although the low sample size and high variance makes it difficult to conclude.

As for scholarships, there was no correlation between fellowship allowances and age (**Fig 40A**, p>0.3) or gender (**Fig 40B**, p>0.6). However, a correlation was found for origin (**Fig 40C**, p<0.05, Eta = 16%) and PhD lab (**Fig 40D**, p<0.05, Eta = 15.4%): European postdocs had a median fellowship allowance of 2900 euros, while non-European postdocs had a median fellowship allowance of 2000 euros. A similar trend but smaller in size was observed regarding PhD lab. Postdocs who graduated in Europe had a median fellowship allowance of 2670 euros, while the postdocs who graduated outside of Europe had a median fellowship allowance of 2600 euros. Conversely, no correlation was observed between fellowship allowances and overall experience (**Fig 40E**, p>0.6) or MPS experience (**Fig 40F**, p>0.4).

**Figure 40.**
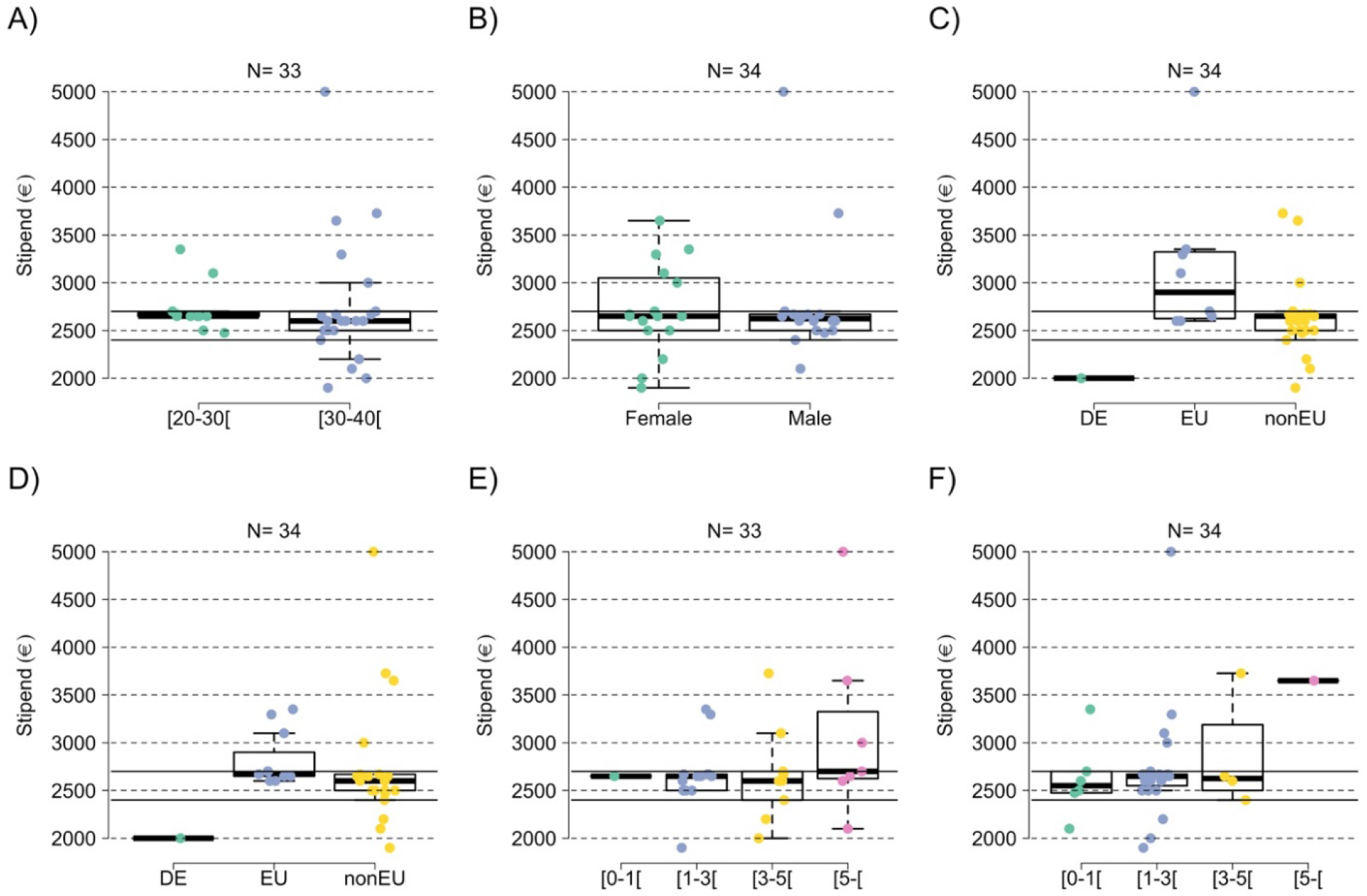
Monthly fellowship allowances. Distributions given based on age **(A)**, gender **(B)**, origin **(C)**, PhD lab **(D)**, overall experience **(E)** and MPS experience **(F).** Age is displayed in years and both experiences in years post PhD graduation. Origin corresponds to Germany (DE), a country in the European Union (EU) or countries outside of the European Union (nonEU). PhD lab corresponds to graduation from one MPI (MPS), from a German institution (DE), from a European institution (EU) or from a non-European institution (nonEU). Monthly allowance is displayed in Euros.

For fellowship allowances, the main correlation was related to the funding sources (Fig 41 – p<0.05, Eta = 24.4%). Fellowships from a European Research Foundation (ERF) had a median of 3296.1 euros, while the Alexander von Humbold (AvH) fellowships had a median of 2650 euros. International grants (IG) had a median of 2400 euros, and the German Research Foundation (GRF) fellowships had a median of 2250 euros.

**Figure 41.**
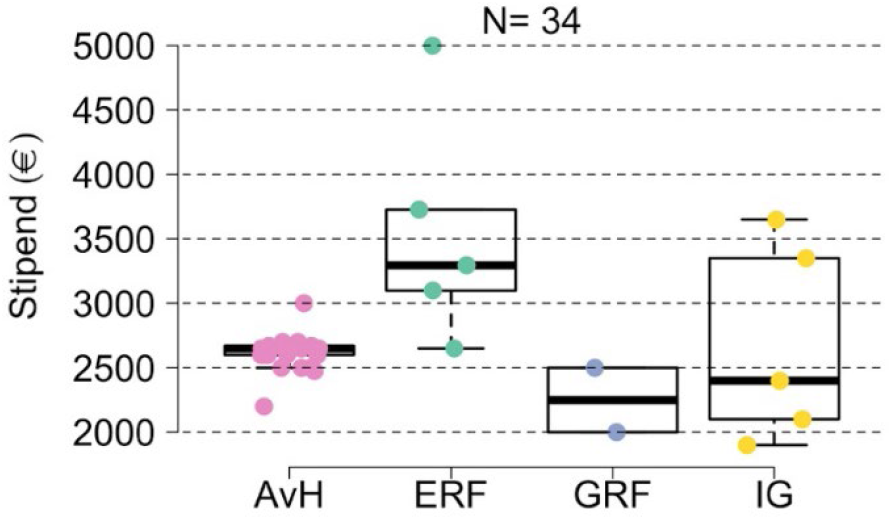
Monthly fellowships allowances according to funding source. AvH, Alexander von Humbold; ERF, European Research Foundation; GRF, German Research Foundation; IG, international grants. Monthly allowances are displayed in Euros

##### c) Initial stipend duration

All MPS scholarships and third-party fellowships started after 2015, when the MPS took the decision to ban stipends for PhD students, and limit their use to foreign scientists with short-term residency^④^.

The initial duration of stipends ranged from less than six months to more than 2 years (**Fig 42**). Most MPS scholarships had an initial duration between either 1.5 and 2 years (54.1% – 20/37) or 6 months and 1 year (29.7% – 11/37). Few scholarships were given for less than 6 months, between 1 and 1.5 years or for more than 2 years (5.4% – 2/37 in each case) (**Fig 42A**). The majority of third-party fellowships however were mostly longer than 1.5 years (75% – 27/36 were between 1.5 and 2 years, and 16.7% – 6/36 for more than 2 years), and few were shorter than 1 year (5.6% – 2/36) or shorter than 6 months (2.8% – 1/36) (**Fig 42B**).

**Figure 42.**
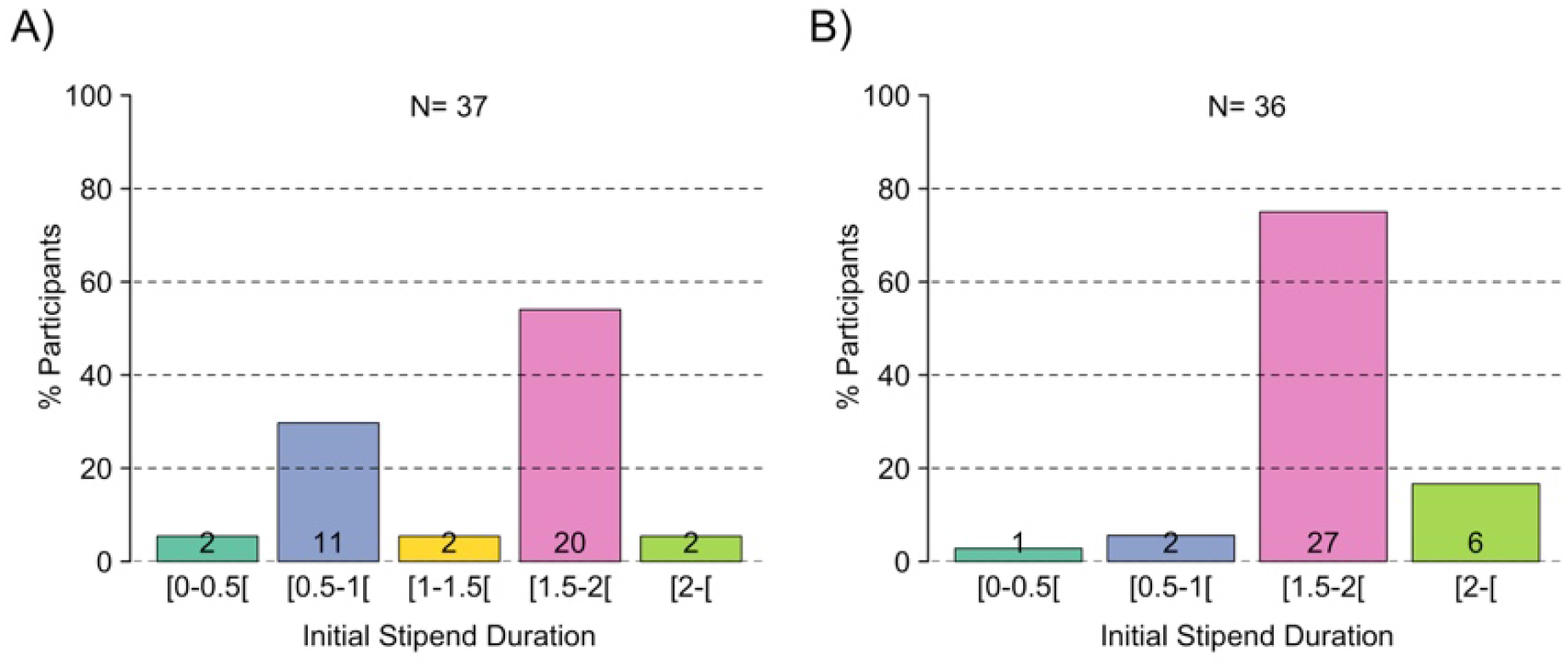
Initial duration of stipends. Distribution given for MPS scholarship **(A)** and third-party fellowships **(B).** Numbers in each bar graph refer to the number of answers given.

The open comment section suggested that stipends may be used as a first hiring strategy before providing postdocs with a work contract. It also emphasized that the accumulation of stipends exceeded by far the definition of short stay. However, related questions were not asked in the survey, making it difficult to evaluate how widespread these situations were.

> *“I was on stipend for my first 3 years before being moved to contract”*
>
> — Participant’s comment

> *“I have been on stipend (MPI and own fellowships) for several years before getting the contract”*
>
> — Participant’s comment

##### d) Social conditions

As stated in the definition given by the PostdocNet, stipends usually do not include social security coverage. To evaluate differences between stipend and contract holders, two main points were surveyed: (i) social and employee benefits (**Fig 43A**), and (ii) the prior knowledge to both the employment on a stipend and the social/work conditions associated with this type of employment (**Fig 43B**).

**Figure 43.**
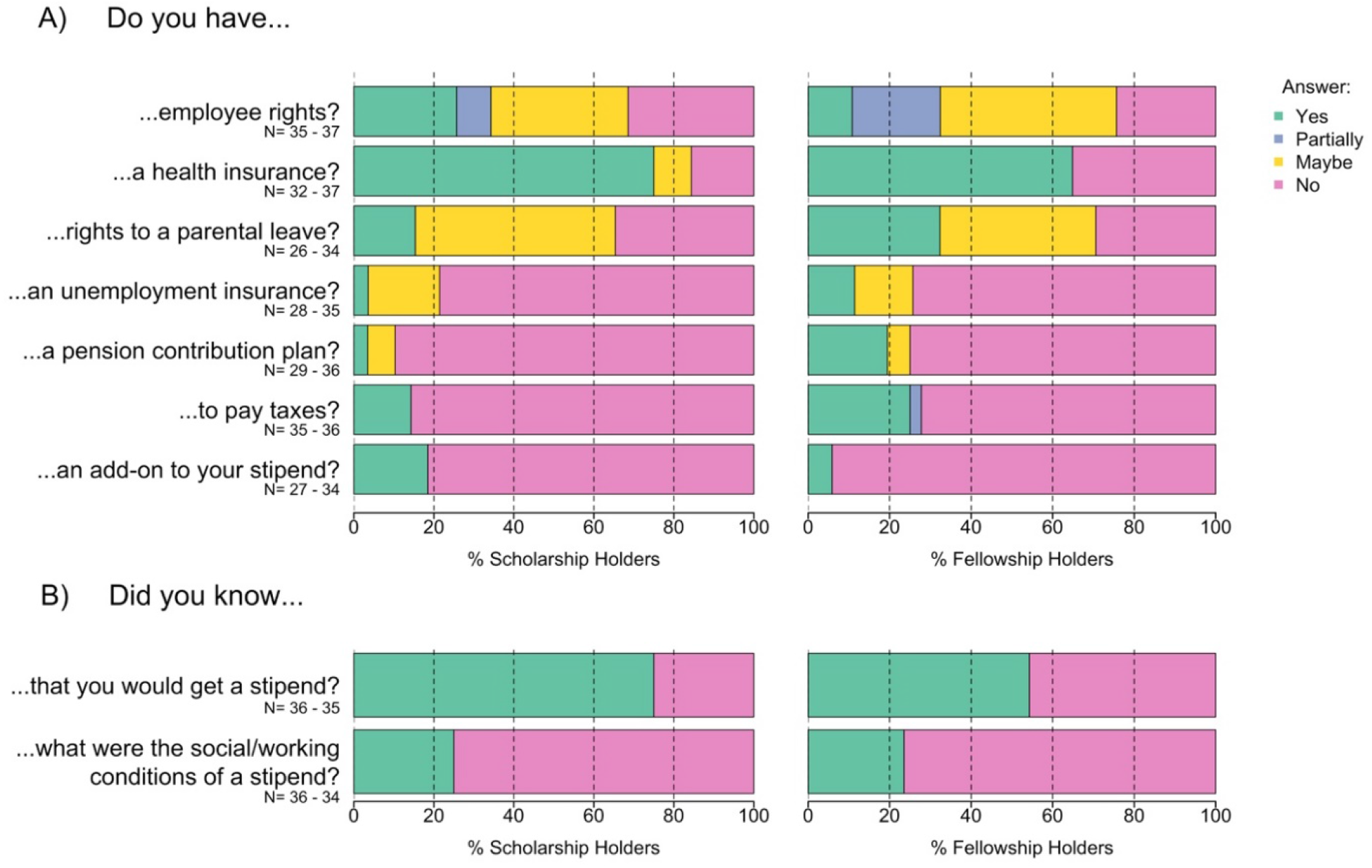
Social advantages and awareness of postdocs employed on a stipend. Responses related to the access to social rights and benefits **(A)** and related to the awareness of employment conditions **(B).**

The focus of the first question was whether employee rights were comparable between stipend and contract holders. Employee rights were defined as the opportunities given to employees to participate in their institute’s life (e.g. voting rights in the elections of the works council and Ombudsperson), to perform their work to their fullest capacity (equipment usage rights), and to benefit from the support of the MPS to balance work and private life (e.g. childcare support).

The vast majority of stipend holders reported that they did not have the same employee rights as their peers on contract. Only 25.7 % (9/35) of scholarship holders and 10.8% (4/37) of fellowship holders stated to have full employee rights in their respective institute. In contrast, 31.4% (11/35) of scholarship holders and 24.3% (9/37) of fellowship holders reported that they did not have any of these rights and 42.9 % (15/35) of scholarship holders and 64.9% (24/37) of fellowship holders reported that they were either uncertain about their rights, or had only partial rights.

The second question investigated whether stipend holders had health insurance. Surprisingly, 15.6% (5/32) of scholarship holders and 35.1% (13/37) of fellowship holders reported that they did not to have any health insurance and 9.4% (3/32) of scholarship holders reported that they were not sure about whether they had any health insurance.

Importantly, only 15.4% (4/26) of scholarship holders and 32.4% (11/34) of fellowship holders reported that they had the possibility to take a parental leave, while 34.6% (9/26) of scholarship holders and 29.4% (10/34) of fellowship holders responded that they did not have this possibility. Finally, 50% (13/26) of scholarship holders and 38.2% (10/13) of fellowship holders were unsure.

Most stipend holders reported that they did not have unemployment insurance (78.6% – 22/28 for scholarships, 74.3% – 26/35 for fellowships) nor pension contribution plan (89.7% – 26/29 for scholarships, and 75.0% – 27/36 for fellowships). Remarkably, 14.3% (5/35) of scholarship holders and 25.0% (9/36) of fellowship holders stated that they were paying taxes.

Few postdocs (18.5% – 5/27 of scholarship holders and 5.9% – 2/34 of fellowship holders) were receiving fringe benefits to their MPS scholarship from their institute.

Most postdocs knew before starting their stipends that they would be employed on a stipend (75% – 27/36 of scholarship holders and 54.3% – 19/35 of fellowship holders). Although, most of them reported that they were not informed of the work/social conditions associated with this type of employment (75% – 27/36 for scholarships holders and 76.5% – 26/34 for fellowships holders).

> *“I think the stipend of postdoc should be the same as the employment contract after tax. Because all the postdocs did a similar contribution to research and spend the same time for work. But the salary of a postdoc with employment contract has a higher salary after-tax, and their salary will increase with the years.”*
>
> — Participant’s comment

> *“Winning a fellowship, such as AvH or EMBO should be a privilege for the Postdocs. But unfortunately, postdocs with the fellowship pay their own health insurances and most of the times health insurances are so expensive.”*
>
> — Participant’s comment

> *“If I knew that I will lose all benefits after receiving a fellowship, I would have never put time and effort to apply for my fellowship.”*
>
> — Participant’s comment

> *“There should be some support to help top-up those who are able to secure their own funding, especially when the net monetary amount drops below the MPS standard.”*
>
> — Participant’s comment

> *“I think the lack of health insurance in the stipendium is critical.”*
>
> — Participant’s comment

###### Summary of employment conditions data

- Most MPS postdocs were employed on a fix-term contract
- The type of employment correlated with:

- Age and experience: younger/less experienced postdocs were more often employed on stipends than older/more experienced postdocs
- Origin: non-European postdocs were more often employed on stipends than European and German postdocs
- Fixed-term TVöD contracts:

- Variability in recognizing previous PhD and postdoctoral experience(s) o Contributing factors were origin, employment type of previous experience(s), gender
- Lack of transparency in classifying postdocs in wage groups (*Entgeltgruppe*) and wage levels (*Stufe*)
- Stipends:

- Lack of transparency in the attribution of stipends and their related work and social conditions
- On average, fellowship allowances were higher than MPS scholarship allowances
- On average, stipends allowances were lower than salaries of most fixed-term contract (E13.3)
- The allowance of all MPS scholarship holders was below the salary received by postdocs on fixed-term contract holders (E13.3)
- Female stipend holders were mainly fellowship holders while male stipend holders were mainly MPS scholarship holders

## III. Discussion/Main findings

In 2006, the MPS validated the “Guidelines for the Postdoc Stage” to clarify the postdoctoral period in the MPS, by presenting what all postdoctoral researchers can expect in terms of scientific independence, support, mentorship, coaching, working conditions and career planning during their stay at their respective MPI.

After its founding, the PostdocNet needed data on the community that it represents. These data focused exclusively on postdoctoral researchers and brought information on demographics and employment conditions.

Eventually, this report will provide insights for postdoctoral candidates, MPS postdoctoral researchers, heads of research laboratory and/or MPS administration to establish, improve and maintain optimal working conditions for postdocs.

### • Gender bias

Overall, there were slightly more male postdocs than female postdocs (see **Fig 2A**). In addition, the proportion of female postdocs dropped significantly as postdocs became more experienced (see **Fig 10A**).

These findings are consistent with the “**leaky pipeline**” effect or when women drop out of a research career at various times^⑤-⑧^.

In an article written on March 07, 2014, the MPS acknowledged the fact that more female scientists in leading positions were needed and was aiming to increase this number by five until 2017^⑨^.

Indeed, for postdocs between the age of 20 to 30 years, the proportion of female postdocs was higher than the proportion of male postdocs (see **Fig S2**). This could reflect a change in the hiring policy over the last 6 years. As an example, in November 2017, the MPS launched the Lise-Meitner excellence program that aims to “offer young female scientists unique opportunities” including free scientific development, long-term career security and clear career prospects^⑩^.

Interestingly, we observed another gender bias in the classification of wage levels for postdocs that were employed on a fixed-term TVöD-based contract. Here, male postdocs with 1 to 3 years of experience tended to be placed in a lower wage level (*Stufe*) than female postdocs with the same amount of experience (see **Fig 34B**). Additionally, male postdocs were also employed significantly more often on MPS scholarship than female postdocs, who were employed more on third-party fellowship (see **Fig 37B**).

> *“I am a bit surprised that almost all of the non-EU male gets stipend whereas all girls and EU-males get a contract. I would like to know why is this difference exists in the first place?”*
>
> — Participant’s comment

The MPS proposes a great variety of offers to support women and more broadly to favor equal opportunity^⑪^. Whether the changes in hiring and equal opportunity policy will have a consequence on the gender biases reported here will only be visible in future surveys.

### • Employment disparity between stipend and contract holders

On multiple aspects (e.g. incomes or social benefits), stipends holders are **less advantaged** than their colleagues employed on a fixed-term TVöD-based contract.

> *“Obtaining the money for my own salary for duration of total 4 years is not an easy process and it took a lot of effort to have this done. The MPI benefits from my work as my papers will be published with the name of MPI and on every international conference I will attend (paying from my own money) I will mention MPI. I am happy about my project and I can benefit from great facilities provided by MPI. However, what doesn’t feel right is that I’m cut out of any benefits employees of MPI have. This in particular includes double the price of guest house, no right to register for the language course or any other course organized by MPI. The inability to have any kind of help regarding obtaining better health insurance is particularly upsetting […]. Situation like this is heavily discouraging researchers to invest their time in obtaining the scholarships which is a necessary element on a way towards scientific independence and obviously is a beneficial for the MPI as less money goes into paying researchers salaries.”*
>
> — Participant’s comment

Postdoctoral researchers on MPS scholarships or third-party fellowships reported that they have **limited access** to social benefits, parental leave, pension and employment scheme (see **Fig 43A**).

In addition, postdocs on stipends grieved about the **discrepancies** between their own incomes and the salary received by fixed-term TVöD-based contract holders. Additionally they stated that getting a stipend does **not seem to offer them clear advantages**.

> *“I can use the equipment and the labs as equal as anyone in the department. However, I am not entitled to benefit from the health services, parental leave, Ombudsman and business car rental.”*
>
> — Participant’s comment

> *“Previously I held an EMBO LTF for 2 years, this hadn’t allowed me for any benefits and had cause a lot of issues with tax, health insurance, institute car usage etc.”*
>
> — Participant’s comment

> *“I think the stipend of postdoc should be the same as the employment contract after tax. Because all the postdocs did a similar contribution to research and spend the same time for work. But the salary of a postdoc with employment contract has a higher salary after-tax, and their salary will increase with the years.”*
>
> — Participant’s comment

> *“After many years with different combination of stipend and contract based payment, I would recommend the new comer with only contract based employment because the potential social benefit that contract holder could have is much more huge compare to the stipend holder.”*
>
> — Participant’s comment

### • Lack of transparency

There was a widespread discontent among postdoctoral researchers about a **lack of information** provided by the MPI administrations about their future employment conditions.

> *“There is no clarity on the basis of which the decision of offering a contract or stipend is made. Surprisingly, most of us are not even informed about these two distinct categories of postdocs at the time of joining.”*
>
> — Participant’s comment

> “Being from overseas and not being informed of your rights or how you are placed in the certain stüfe level is a common practice at our MPI. It’s only after some time you realize that none of your PhD or postdoc experience has actually been taken into account! But being from overseas, you place your trust in an organization that is in no way trying to get a fair deal for you.”
>
> — Participant’s comment

> “Despite years of research experience elsewhere and my publication record, nothing was taken into account in assigning me to E-13/1. I only realized later, when I began employing researchers on my own projects, that researchers with no experience beyond Ph.D., or even no Ph.D., are also employed from the same level, This seems unbalanced and unfair, and is a black mark against the MPI’s approach to pay transparency and fairness.”
>
> — Participant’s comment

In “*A career in science at Max Planck*,” the MPS had succinctly presented the different employment conditions that a postdoc candidate can expect by applying in one of the MPIs^⑫^.

However, more than 75% of stipend holders were **unaware of the work/social conditions** associated with this type of employment in their institute (see **Fig 43B**). In addition, 53% of contract holders did not know to which wage level they would be assigned to before starting their postdoctoral position (see **Fig 35**).

> *“Although I was informed about the category of my employment, there was little information given about what this means and how it effects my social benefits etc.”*
>
> — Participant’s comment

> *“If I knew that I will lose all benefits after receiving a fellowship, I would have never put time and effort to apply for my fellowship.”*
>
> — Participant’s comment

> *“It is crucial to be informed about the details of the contract before one actually decide to take the position. Once one decides to take the position and moves to the new city then gets the contact and few days to sign it, it is very late to go back in case of problematic situation.”*
>
> — Participant’s comment

Overall, these results suggested **poor transparency** and a **lack of information about work conditions in the employment procedures for postdocs**.

From the survey data, this concerned German as well as non-German postdocs, equally. However, this lack of transparency can be especially detrimental for postdocs from abroad who represented more than 72.6% of the researchers (the proportion of international found in the survey was in line with the information provided by the MPS^⑬^).

> *“Within my department, all German postdocs have a higher salary than non-German postdocs. I know this is in part because researchers with previous experience at the MPI are placed automatically into Stufe 3 and in part because our administration rarely considers PhD experience obtained elsewhere as equal to German PhDs. […]. To me this seems highly problematic and I wonder even to what extent it is lawful, given EU non-discrimination law for example.”*
>
> — Participant’s comment

Moreover, postdocs reported **inconsistencies concerning the recognition of previous experiences**, and even more so depending on postdoc’s origins (see **Fig 23, 26, 29)**.

> *“I was trying to get my previous experienced recognized and I provided my work contracts, but they were not considered. I got no explanation other than it was probably because my jobs were not at a German institution.”*
>
> — Participant’s comment

> *“I was informed about my category [Entgeltgruppe], but not about my subcategory [Stufe]. Afterwards I had some struggles with our administration staff, because it seems that they do not have consistent standards of evaluation. The classification of postdocs in a subcategory, meaning the consideration of previous work experience, seems to be rather arbitrary.”*
>
> — Participant’s comment

Additionally, PhD experience, **previous experience(s)** gained abroad or outside of the TVöD wage agreement **was not fully recognized** for the classification to the respective wage level (*Stufe*). Results show that 58% of postdocs (207/359) had a wage level (*Stufe*) lower than their overall (doctoral and postdoctoral) experience (see **Fig 30A**). This was most likely due to a lack of recognition of their doctoral experience.

> *“The two years on a stipend, did not then count as experience in the eyes of the MPG, despite having been a postdoc in the MPG those two years, so when I switched contracts, I started at the level of a newly minted German PhD student, which I clearly was not.”*
>
> — Participant’s comment

> *“PhD experience should count towards the stufe level within reason. At least 3 years of experience should be recognized. Also a postdoc means you have to have a PhD, not a master.”*
>
> — Participant’s comment

> *“During my PhD, I was receiving a scholarship, which was not acknowledged as work experience.”*
>
> — Participant’s comment

> *“I was on a stipend during Phd and the first two postdoc years. When I entered TVöD I was put in e13/1! This should not be legal since at least the former two postdoc years were on the exact same project. How can that not be count at as work experience?”*
>
> — Participant’s comment

> *“It seemed arbitrary that for some people PhD time in Germany counted towards a higher Stufe. But not in my case, I started with Stufe 1.”*
>
> — Participant’s comment

Finally, 20% of the postdocs were confident that their experience was taken into account for the determination of their wage level (Stufe) (see **Fig 35**).

> *“I had to ask for a reassessment of my ‘Stufe’ to have my previous work experience fully considered. This was granted then but it obviously required that I knew about the possibility for a reassessment and that I went to my director to ask for it (which is not self-evident after just having started working at the institute).”*
>
> — Participant’s comment

### • Current situation

A preliminary analysis of this survey led to intense discussions within the postdoctoral community and within the MPS. These discussions have **brought awareness to the working conditions and inequalities** that currently exist among postdocs, and has shed some light on the underlying causes.

The PostdocNet 2019-2020 steering group and the speakers of scientific staff representatives of the three MPS sections agreed in June 2020 on a proposal of measures to improve the employment conditions of MPS postdocs. Hopefully, these propositions will be implemented and **guide future actions by the MPS**.

More recently, in November 2020, the MPS announced an increase of the monthly allowance of MPS-scholarships, a first encouraging step towards more equality within the MPS postdoc community. However, this measure leaves unchanged (i) the reduced social benefits of stipend-holders; (ii) the gap between scholarship-allowance and the income of a contract holder (see **Fig 38**) and (iii) the origin’s bias for MPS scholarship attribution (see **Fig 19A** & **37C**).

The difficulties faced by the postdoctoral community are in fact not limited to the MPS. A recent publication in Nature^⑬^ recognizes postdoctoral researchers as the “**research precariat**”, a situation that contrasts with their highly-valued works.

We hope that this report by documenting postdocs concerns, will contribute to improve future working and social conditions for the postdocs in the Max Planck Society, one of the most prestigious research societies worldwide. Today, the MPS has a unique chance to set an example in the academic world by actively participating to the global betterment of an increasingly difficult period of academic research: the postdoctoral phase.

## Methods

The survey contained 32 questions, all of which were made optional, to ensure that each participant could decide whether an answer could jeopardize their anonymity. The full list of the questions is available in supplementary **File S1**. 623 postdocs answered the survey.

The 623 answers were edited to facilitate analysis:

– free answers were edited to the same format (e.g. all numbers were modified to use the same decimal marker);
– when participants answered their MPI without the MPS Section, the second was inferred from the first;
– when participants on stipends answered “I don’t know” to the funding source of the stipend, and “My boss” to the origin of the stipend, and the monthly allowance corresponded to that of an MPS stipend (2100 euros), the funding source was inferred to be MPS (N=5);
– when participants answered the wage category (*Entgeltgruppe)*, the work load and the salary, but not the wage level (*Stufe)*, the later was inferred (N=3);
– when the funding source was a combination of MPS and another institution, it was binned in the category “MPS & Other” to avoid increasing the number of categories containing <5 occurrences (N=10);
– for the precise origin of the stipend, all funding sources cited except the Alexander von Humbold Foundation (N=22) had at most 4 occurrences and thus could not be analyzed, so we only considered the AvH Foundation.

Of those 623 participants, not all responded to all questions, so a set of “key” questions was defined to evaluate representativeness, excluding the MPI membership, and the detailed questions about previous experiences, filled in with free text. A summary of the answers per question and the “key”/“accessories” questions is available in supplementary **File S2**.

Statistical tests were done in R version 3.6.3. Correlations between categorical parameters were calculated using chisq test from the base package, together with Cramer’s V from package “vcd” version 1.4-7 to evaluate effect size. Correlations between categorical and numerical parameters were calculated using Kruskall-Wallis test from the base package, together with Eta to evaluate effect size. Eta was calculated as:

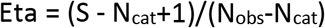

with S as the Kruskall-Wallis test statistic, N_cat_ the number of categories of the parameter and N_obs_ the number of observations.

Figures were made in R version 3.6.3, and groups with less than 10 observations were displayed as empty to avoid biased interpretations from low sample sizes.

Aggregated data for each question and associated tests are provided in supplementary **Files S3-5**, details for groups with samples size lower than 15 were not displayed to protect anonymity of the respondents.

Comments from contract-holder were sorted by “topics” as follow:

<u>Experience recognition:</u>

– PhD years non recognized = 15 comments
– Work on stipend non recognized = 7 comments
– Previous positions non recognized = 20 comments
– Discrepancy with other ‘equal-experienced’ fellow = 7 comments
– Experience taken totally or partially in consideration = 7 comments
– ‘Positive’ situation => often when postdocs asked for = 14 comments

<u>Information received:</u>

– Not enough = 26 comments
– Information was given correctly = 1 comment

<u>Miscellaneous:</u>

– General comments = 11 comments
– Work conditions / Family-life balance = 9 comments

## Supporting information

Supplemental File 3_Demographics

Supplemental File 4_Contract

Supplemental File 5_Stipend

## IV. Acknowledgments

The authors are grateful to all the postdocs who participated to the survey.

The authors also want to thank the 2019-2020 PostdocNet ‘Steering’ group, ‘Social need’ group and the former members of the ‘Survey’ group, especially Dr Abhirup Ghosh and Dr Carlos Mora Duro, for their constructive feedback.

## VI. Supplementary data

### • Supplementary tables

**Abbreviations (alphabetical order): BM**, Biology and Medical section; **CPT**, Chemistry, Physics and Technology section; **DE**, Germany; **EU**, European Union; **HUM**, Human Sciences section; **MPS**, Max Planck Society; **nonEU**, countries outside the European Union; **PhD lab**, geographic region of PhD graduation.

**Table S1.**
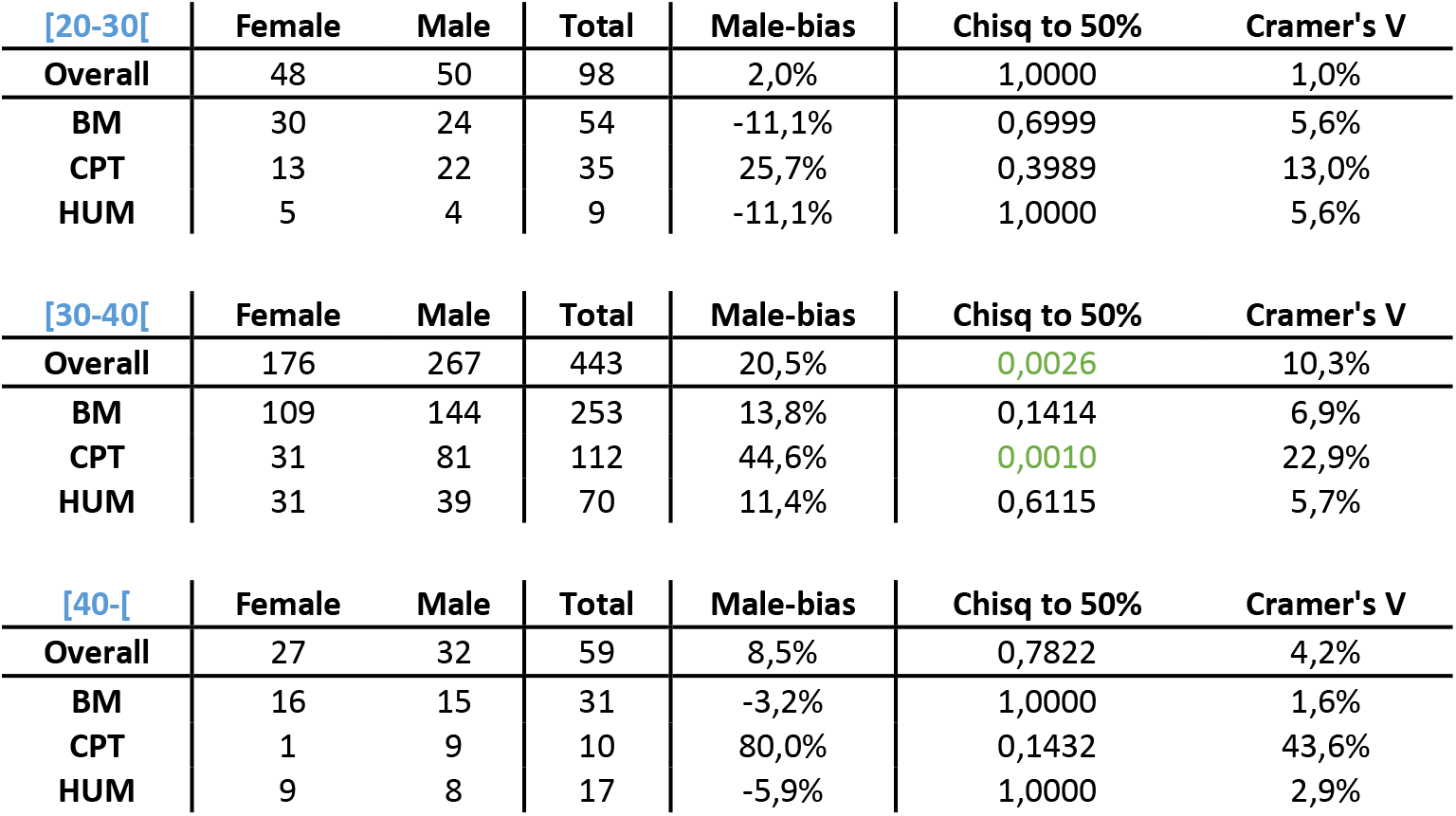
Gender and age groups (in years): 20 to 30 years old group **(top)**, 30 to 40 years old group **(middle)** and older than 40 years old group **(bottom)**.

**Table S2.**
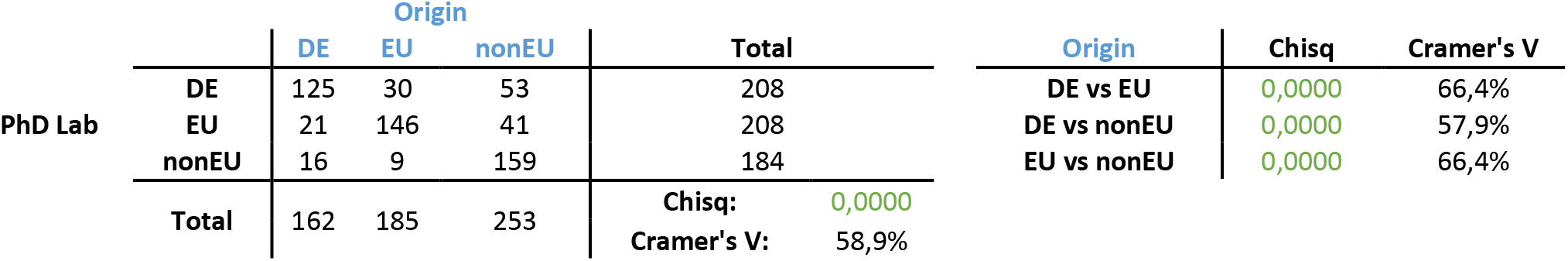
Correlation of origin and PhD lab.

**Table S3.**
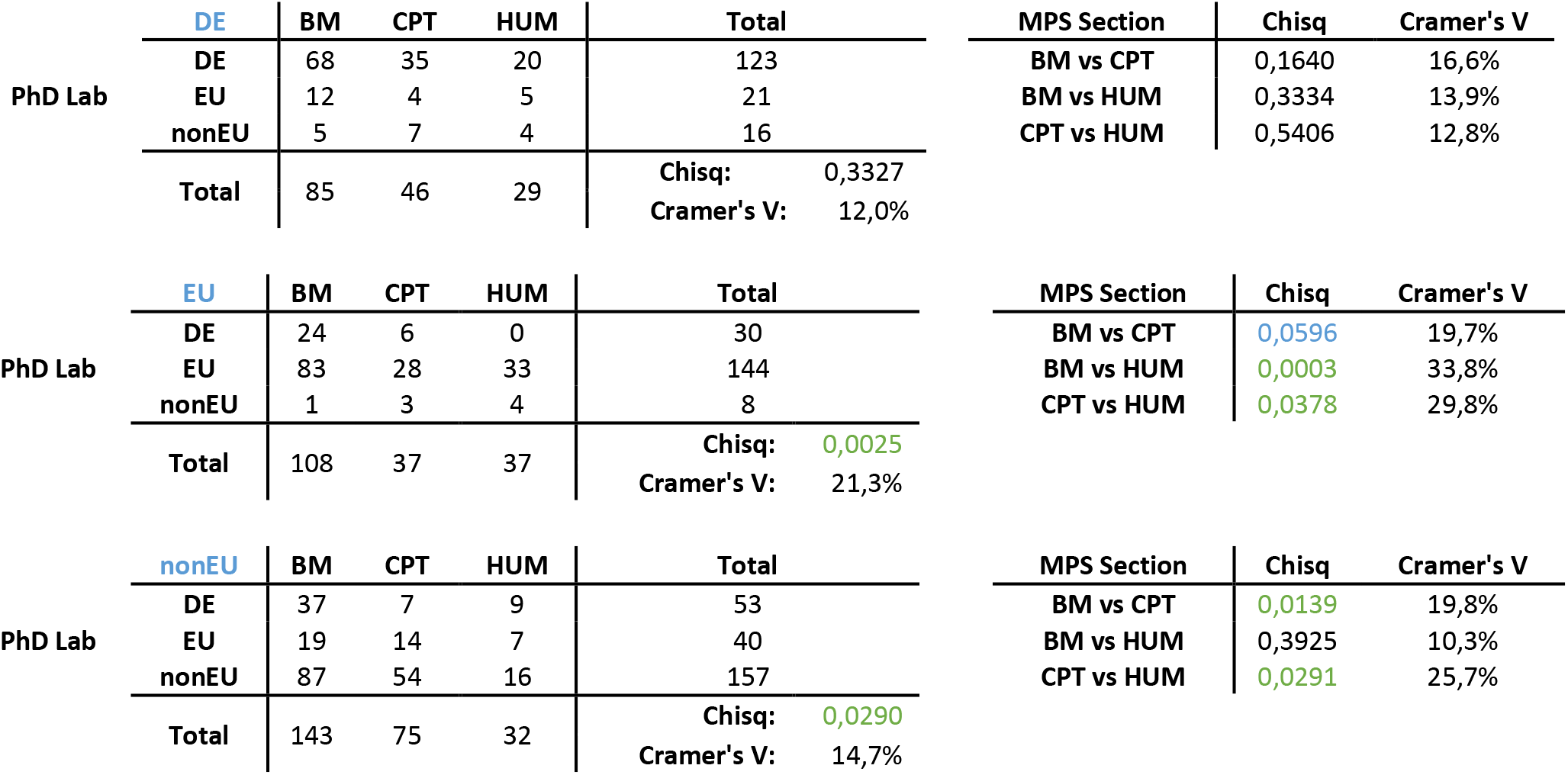
PhD lab and MPS section distributions of German (DE), European (EU) and non-European (nonEU) postdocs.

**Table S4.**
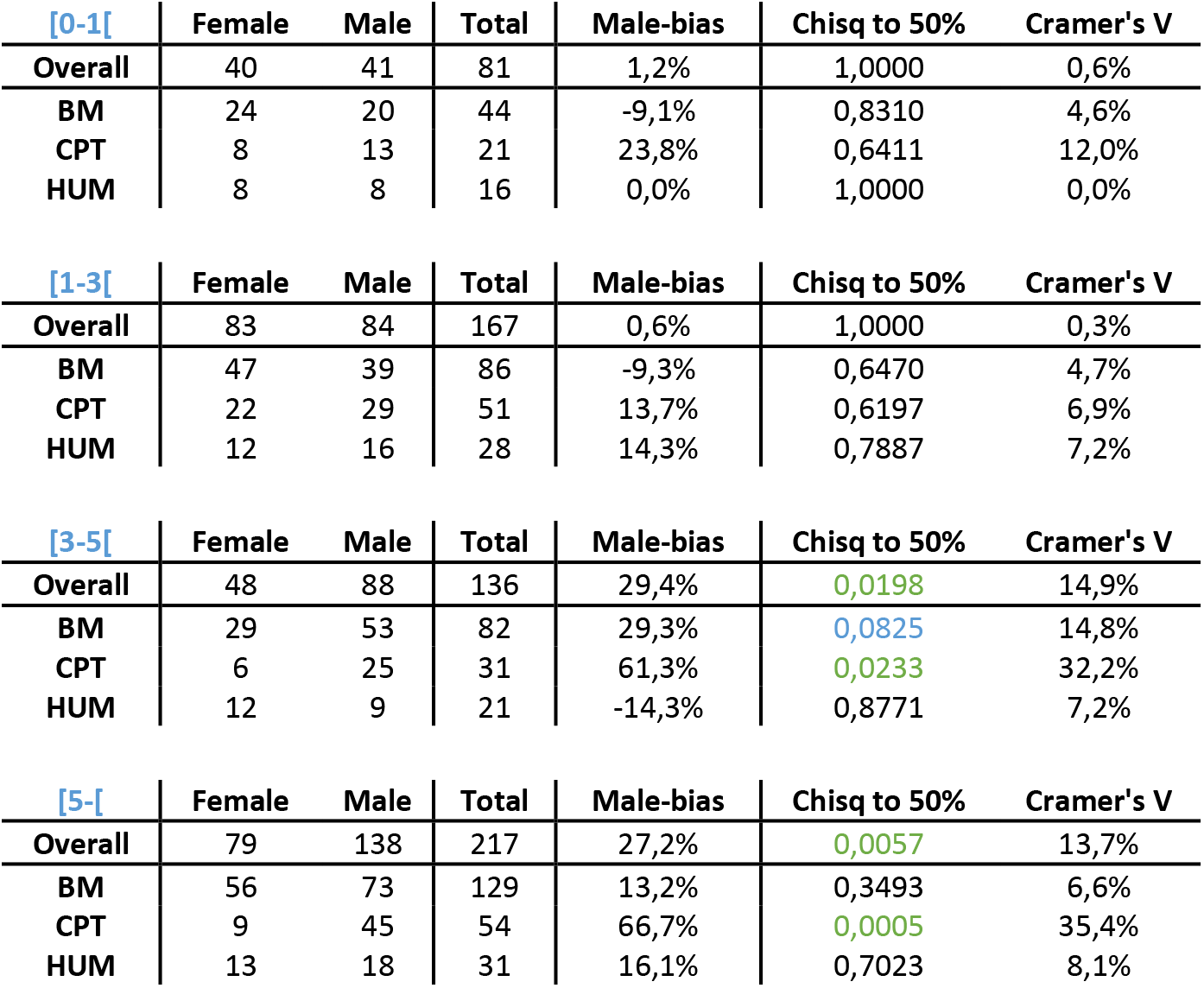
Gender and overall experience. Overall experience corresponds to the years worked after PhD graduation.

**Table S5.** Correlation of overall postdoctoral experience and MPS experience (in year). Overall experience corresponds to the years worked after PhD graduation while MPS experience corresponds to the years worked in one MPI after PhD graduation.

|  |  | Overall experience (in years) |  |  |  |  |  |
| --- | --- | --- | --- | --- | --- | --- | --- |
|  |  | [0-1[ | [1-3[ | [3-5[ | [5-[ | Total |  |
| MPS experience | [0-1[ | 75 | 37 | 16 | 17 | 145 |  |
|  | [1-3[ | 5 | 133 | 54 | 51 | 243 |  |
|  | [3-5[ | 3 | 1 | 66 | 51 | 121 |  |
|  | [5-[ | 0 | 1 | 1 | 95 | 97 |  |
|  | Total | 83 | 172 | 137 | 214 | Chisq<br>Cramer's V | 2,01E-112<br>0,549431 |

**Table S6.** Gender and MPS experiences. MPS experience corresponds to the years worked in one MPI after PhD graduation.

| [0-1[ | Female | Male | Total | Male-bias | Chisq to 50% | Cramer's V |
| --- | --- | --- | --- | --- | --- | --- |
| Overall | 58 | 86 | 144 | 19,4% | 0,1237 | 9,8% |
| BM | 34 | 39 | 73 | 6,8% | 0,8038 | 3,4% |
| CPT | 12 | 30 | 42 | 42,9% | 0,0739 | 21,9% |
| HUM | 12 | 16 | 28 | 14,3% | 0,7887 | 7,2% |

| [1-3[ | Female | Male | Total | Male-bias | Chisq to 50% | Cramer's V |
| --- | --- | --- | --- | --- | --- | --- |
| Overall | 115 | 123 | 238 | 3,4% | 0,7833 | 1,7% |
| BM | 62 | 59 | 121 | -2,5% | 0,9487 | 1,2% |
| CPT | 27 | 45 | 72 | 25,0% | 0,1790 | 12,6% |
| HUM | 23 | 19 | 42 | -9,5% | 0,8271 | 4,8% |

| [3-5[ | Female | Male | Total | Male-bias | Chisq to 50% | Cramer's V |
| --- | --- | --- | --- | --- | --- | --- |
| Overall | 36 | 83 | 119 | 39,5% | 0,0029 | 20,1% |
| BM | 24 | 53 | 77 | 37,7% | 0,0267 | 19,2% |
| CPT | 4 | 18 | 22 | 63,6% | 0,0564 | 33,6% |
| HUM | 7 | 12 | 19 | 26,3% | 0,6235 | 13,3% |

| [5-[ | Female | Male | Total | Male-bias | Chisq to 50% | Cramer's V |
| --- | --- | --- | --- | --- | --- | --- |
| Overall | 41 | 56 | 97 | 15,5% | 0,3492 | 7,8% |
| BM | 35 | 32 | 67 | -4,5% | 0,9311 | 2,2% |
| CPT | 2 | 19 | 21 | 81,0% | 0,0114 | 44,3% |
| HUM | 4 | 4 | 8 | 0,0% | 1,0000 | 0,0% |

### • Supplementary figures

**Figure S1.**
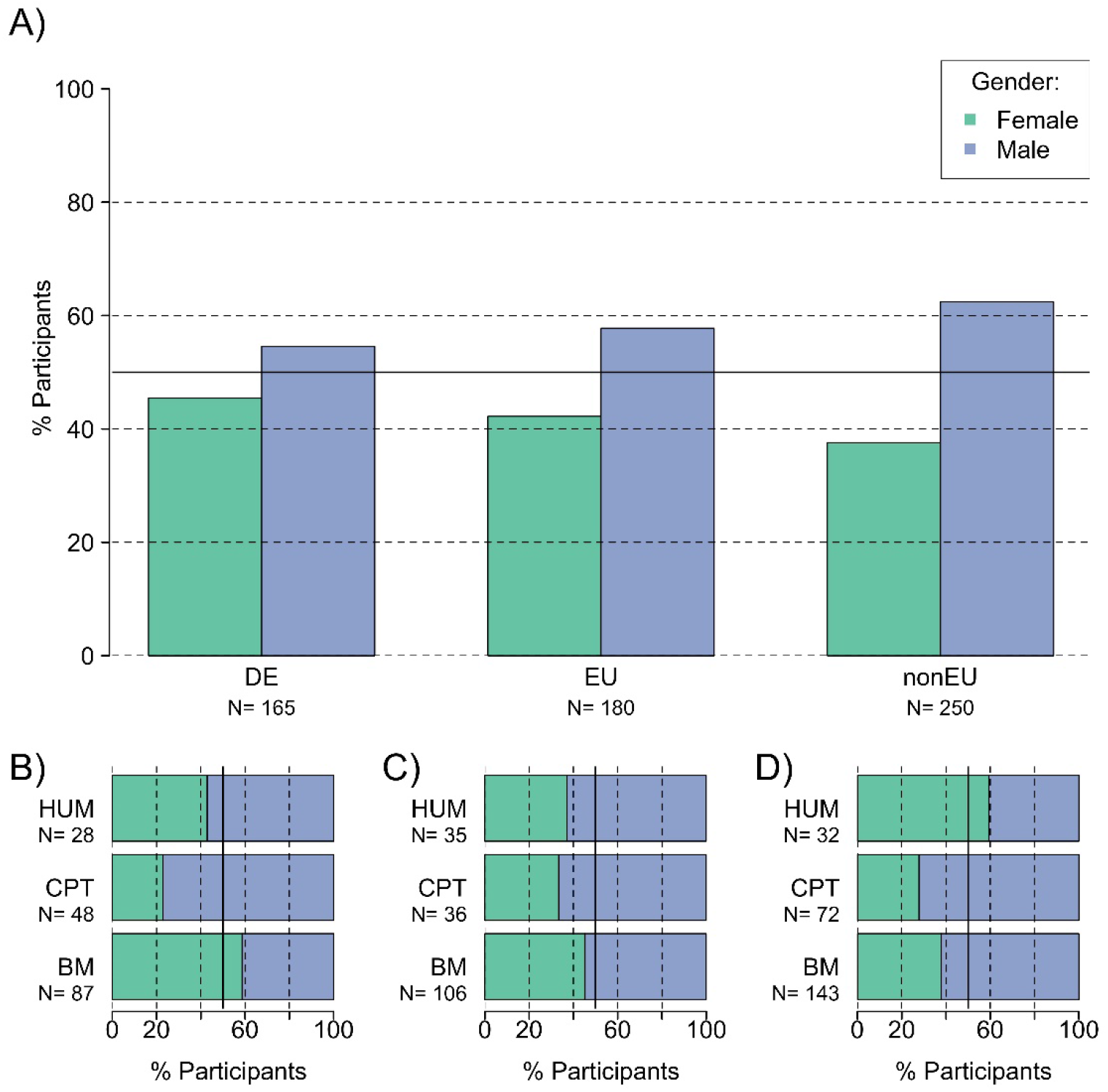
Gender and origin. Comparison of gender distribution based on origin in the MPS overall **(A)** and in each MPS section: German (DE) postdocs **(B)**, EU postdocs **(C)**, non-EU postdocs **(D)**. The thick line corresponds to 50%. BM, Biology and Medical section; CPT, Chemistry, Physics and Technology section; HUM, Human Sciences section; EU, European Union; nonEU, countries outside the European Union.

**Figure S2.**
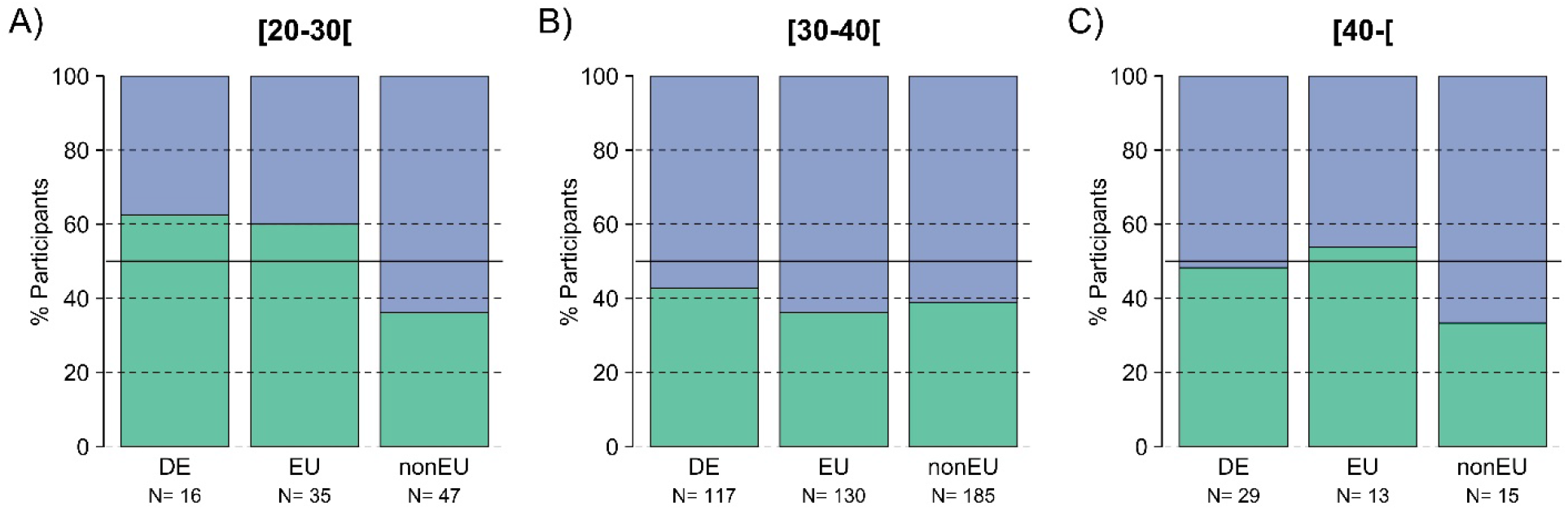
Gender, age groups and origin. Comparison of gender distribution based on origin and age in the MPS overall: 20 to 30 years old group **(A)**, 30 to 40 years old group **(B)** and older than 40 years old group **(C)**. Female postdocs proportion is green while male postdocs proportion is blue. The thick line corresponds to 50%. DE, Germany; EU, European Union; nonEU, countries outside the European Union.

**Figure S3.**
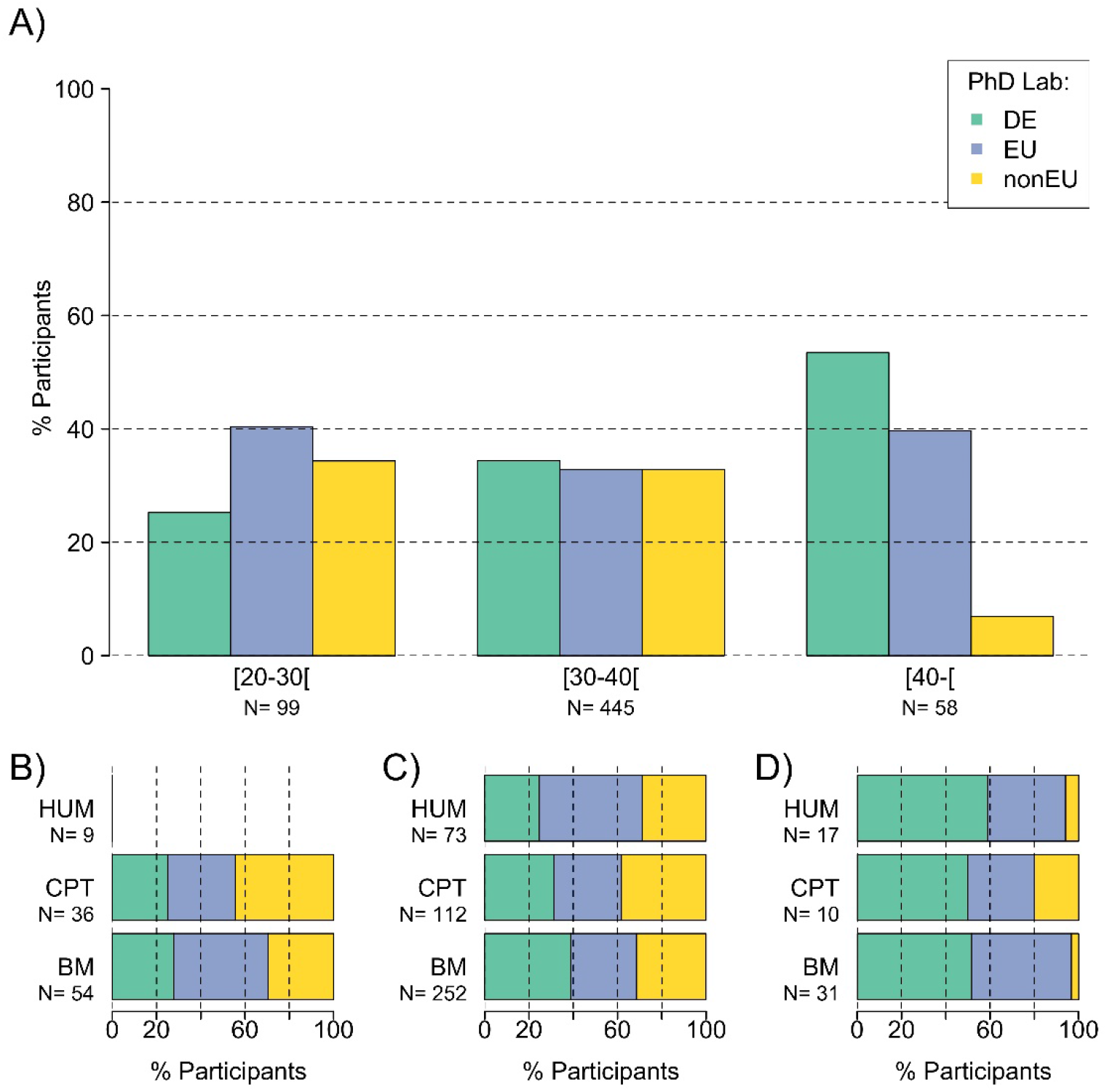
PhD lab and age groups. Comparison of the geographic region of PhD graduation (PhD lab) based on age of the postdocs in the MPS overall (**A**) and in the MPS sections: the 20-30 years old group (**B**), the 30-40 years old group (**C**) and the older than 40 years old group (**D**). Note, statistics with less than 10 data points are not shown on graphs. BM, Biology and Medical section; CPT, Chemistry, Physics and Technology section; HUM, Human Sciences section; DE, Germany; EU, European Union; nonEU, countries outside the European Union.

**Figure S4.**
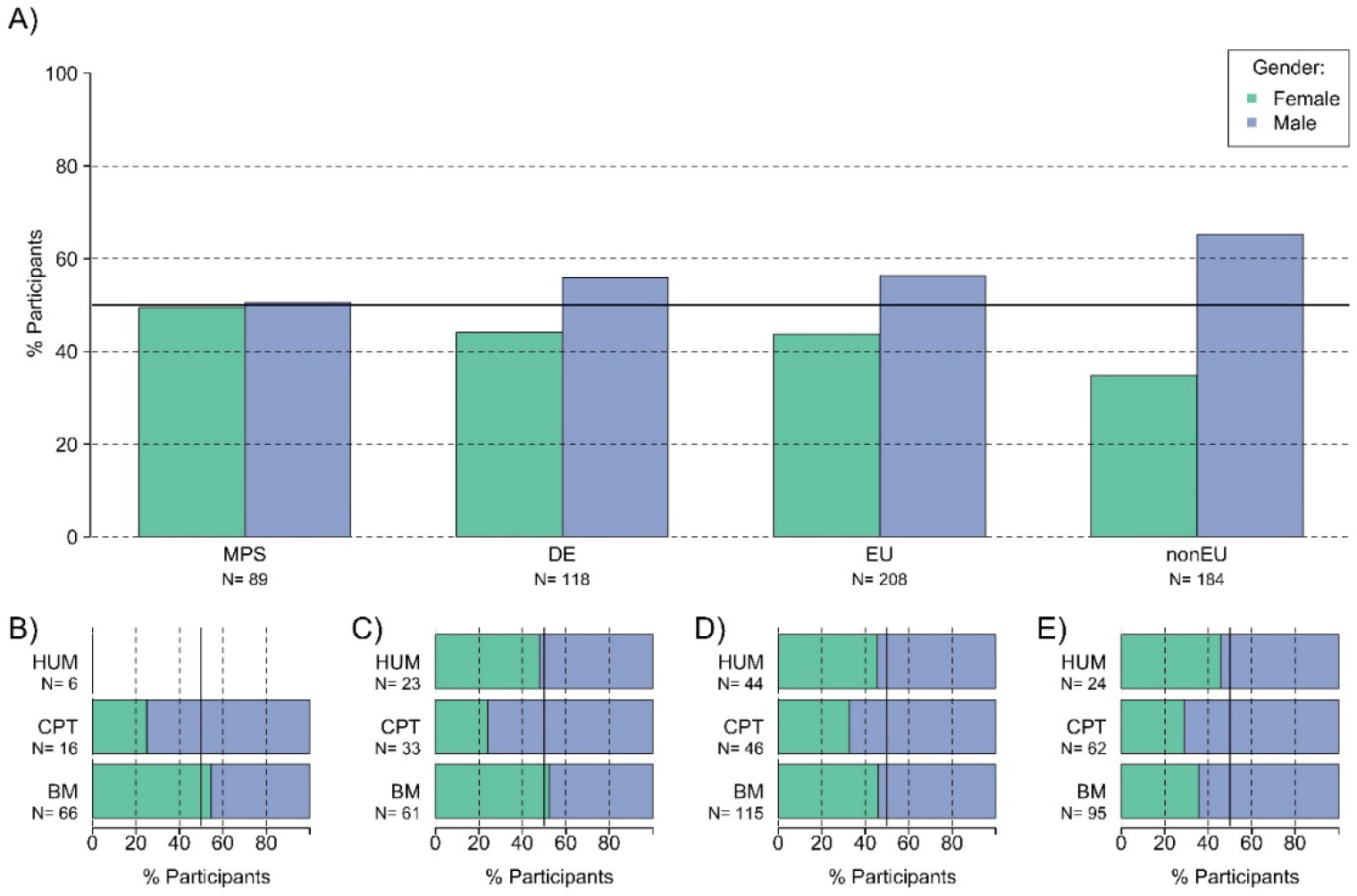
Gender distribution and PhD lab. Comparison of gender distribution based on the geographic region of PhD graduation (PhD lab) in the MPS overall (**A**). In each section, the gender distribution of postdocs is shown for PhD graduation from the MSP **(B)**, from a German institution **(C)**, from an institution in the European Union **(D)**, or from a non-EU institution **(E)**. The thick line corresponds to 50%. Note, statistics with less than 10 data points are not shown on graphs. BM, Biology and Medical section; CPT, Chemistry, Physics and Technology section; HUM, Human Sciences section; DE, Germany; EU, European Union; nonEU, countries outside the European Union.

**Figure S5.**
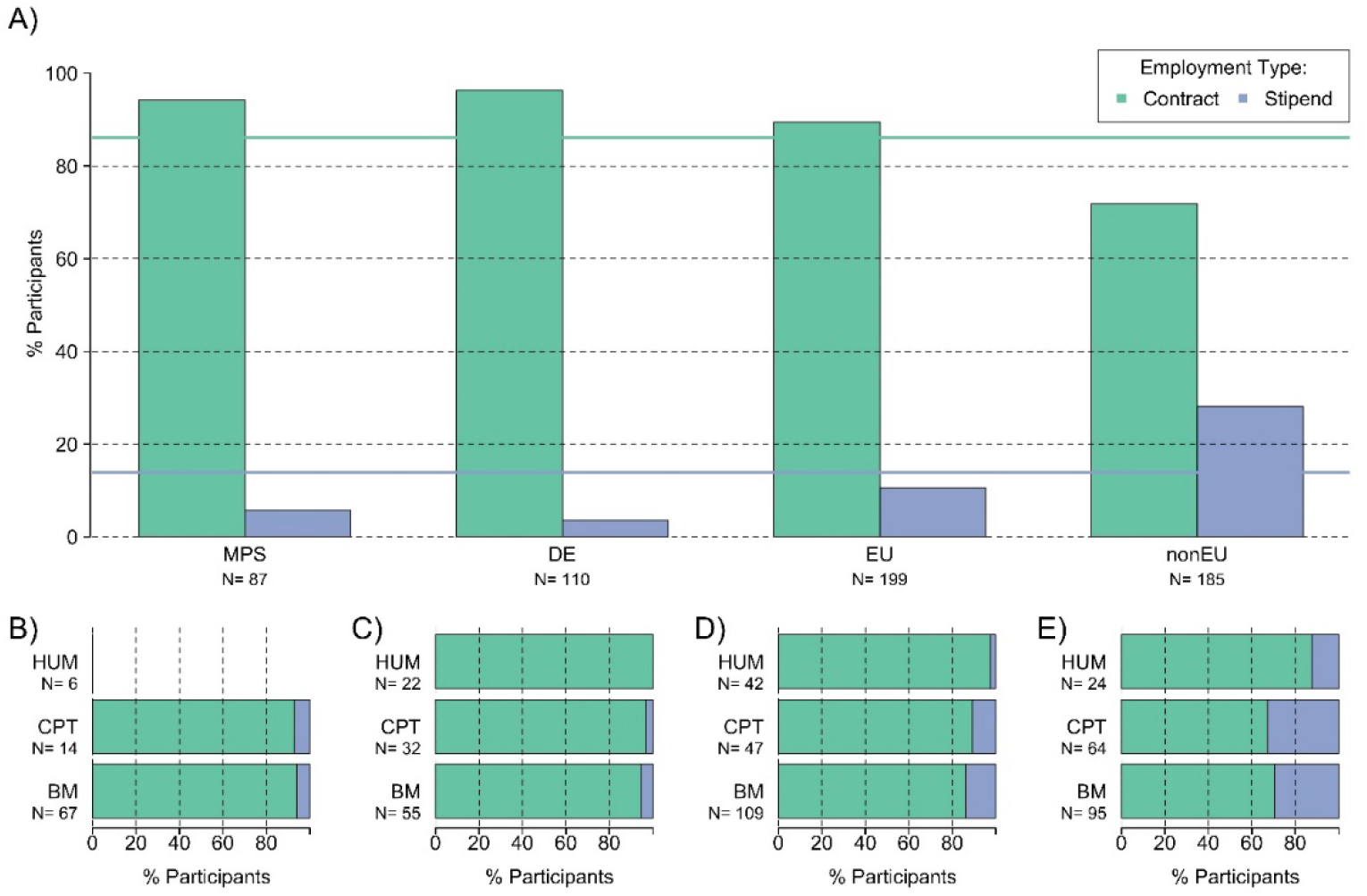
Employment type and PhD lab. Comparison of postdocs employment type (contract or stipends) across the geographic region of PhD graduation (PhD lab) in the MPS overall **(A)**. In each section, the distribution of postdocs being on contract or stipend is shown for PhD graduation from the MSP **(B)**, from a German institution **(C)**, from an institution in the European Union **(D)**, or from a non-EU institution **(E)**. The green and blue lines correspond to the proportion in the overall population of fixed term TVöD-based contracts and stipends respectively. Note, statistics with less than 10 data points are not shown on graphs. BM, Biology and Medical section; CPT, Chemistry, Physics and Technology section; HUM, Human Sciences section; MPS, Max Planck Society; DE, Germany; EU, European Union; nonEU, countries outside the European Union.

**Figure S6.**
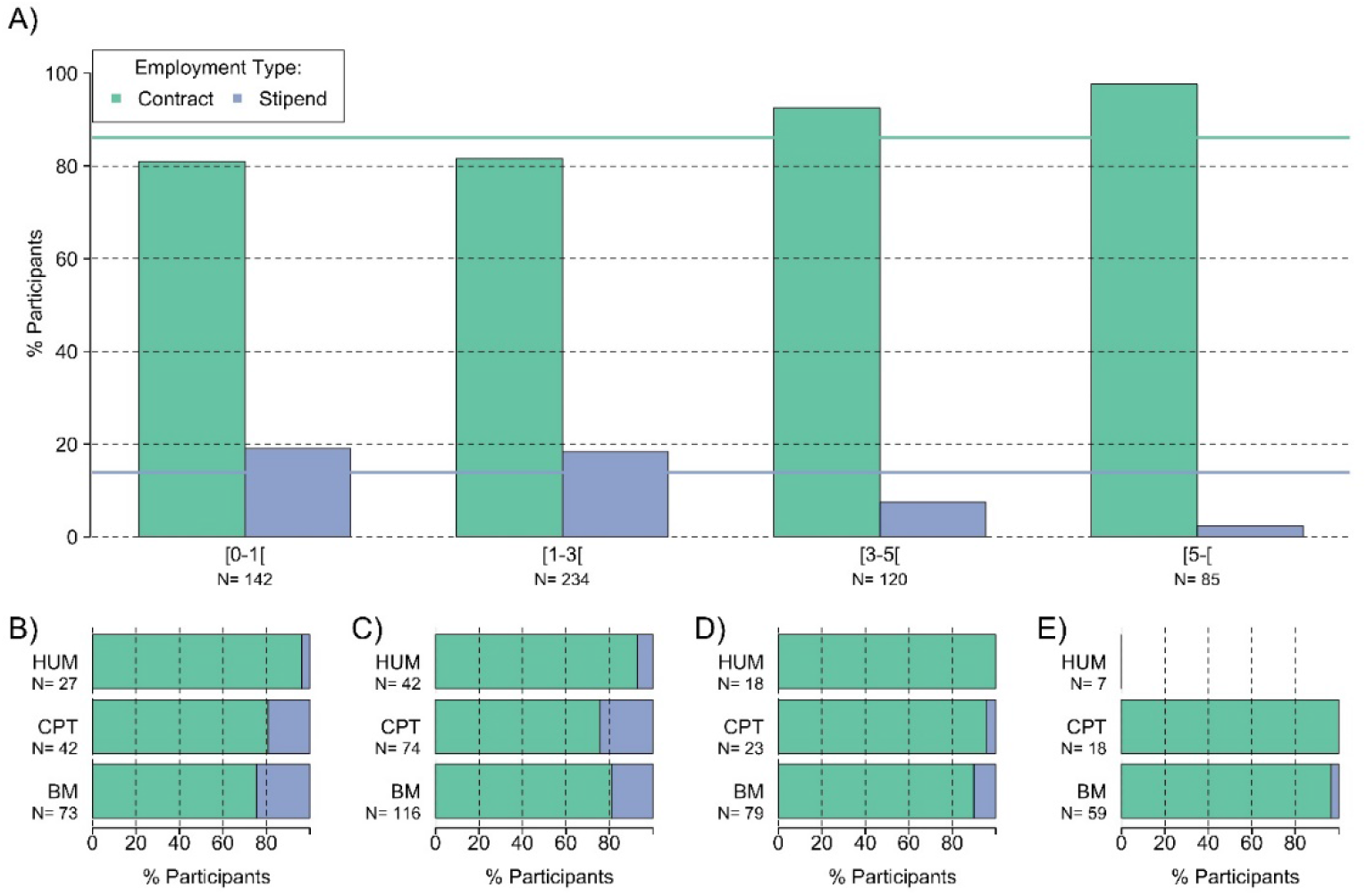
Employment type and MPS experience. Comparison of postdocs employment type (contract or stipends) at the MPS in the MPS overall (**A**) and in each MPS section depending on the MPS experiences (in year): less than one year of experience **(B)**, less than 3 years **(C)**, less than 5 years **(D)** and more than 5 years of MPS experience **(E)**. MPS experience represents the number of years worked within the MPS after PhD graduation. The green and blue lines correspond to the proportion in the overall population of fixed term TVöD-based contracts and stipends respectively. Note, statistics with less than 10 data points are not shown on graphs. BM, Biology and Medical section; CPT, Chemistry, Physics and Technology section; HUM, Human Sciences section.

**Figure S7.**
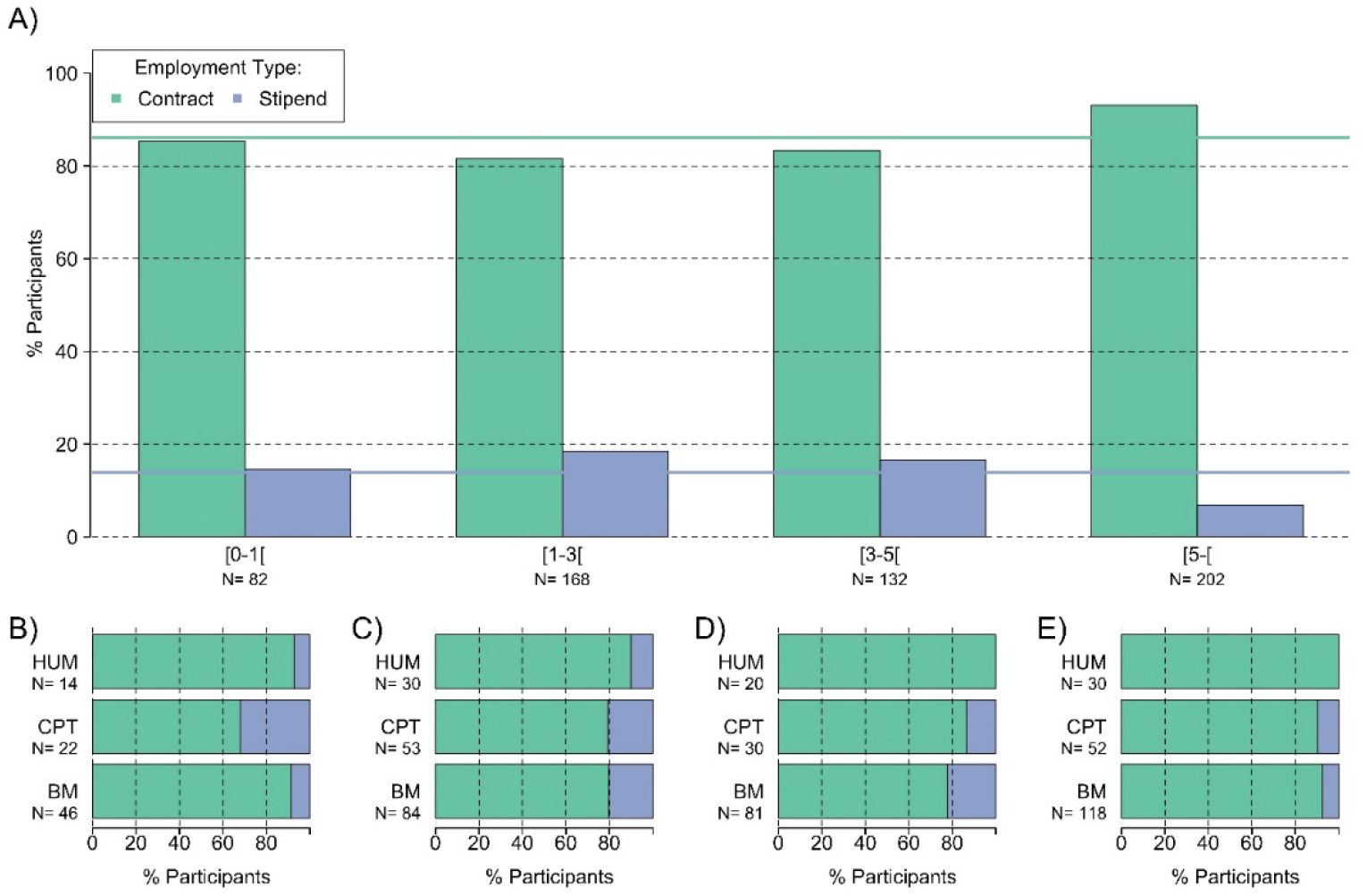
Employment type and overall experience. Comparison of postdocs employment type (contract or stipends) in the MPS overall (**A**) and in each MPS section depending on the overall postdoctoral experiences (in year): less than one year of experience **(B)**, less than 3 years **(C)**, less than 5 years **(D)** and more than 5 years of experience **(E)**. Overall experience represents the number of years worked after PhD graduation. The green and blue lines correspond to the proportion in the overall population of fixed term TVöD-based contracts and stipends respectively. BM, Biology and Medical section; CPT, Chemistry, Physics and Technology section; HUM, Human Sciences section.

**Figure S8.**
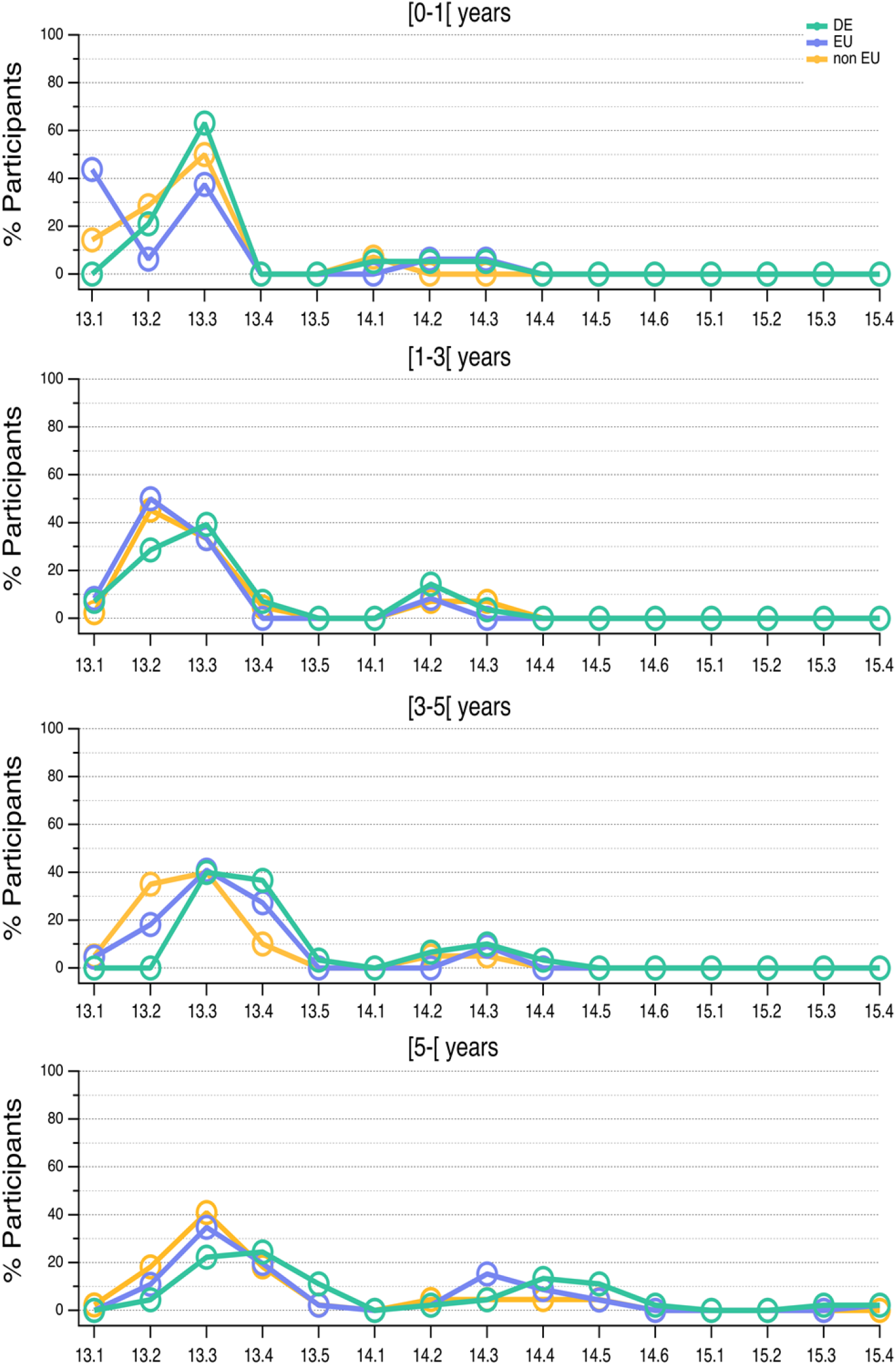
Wage group (*Entgeltgruppe*) and wage level (*Stufe*) assignment, origin and overall experience. Whole distribution of the fraction of German (DE, green), European Union (EU, blue) and non-European Union (non-EU, orange) postdocs in wage levels and groups E13, E14 and E15 for each year of overall experience. In proportion, more international postdocs were found in lower wage levels as opposed to German postdocs.

**Figure S9.**
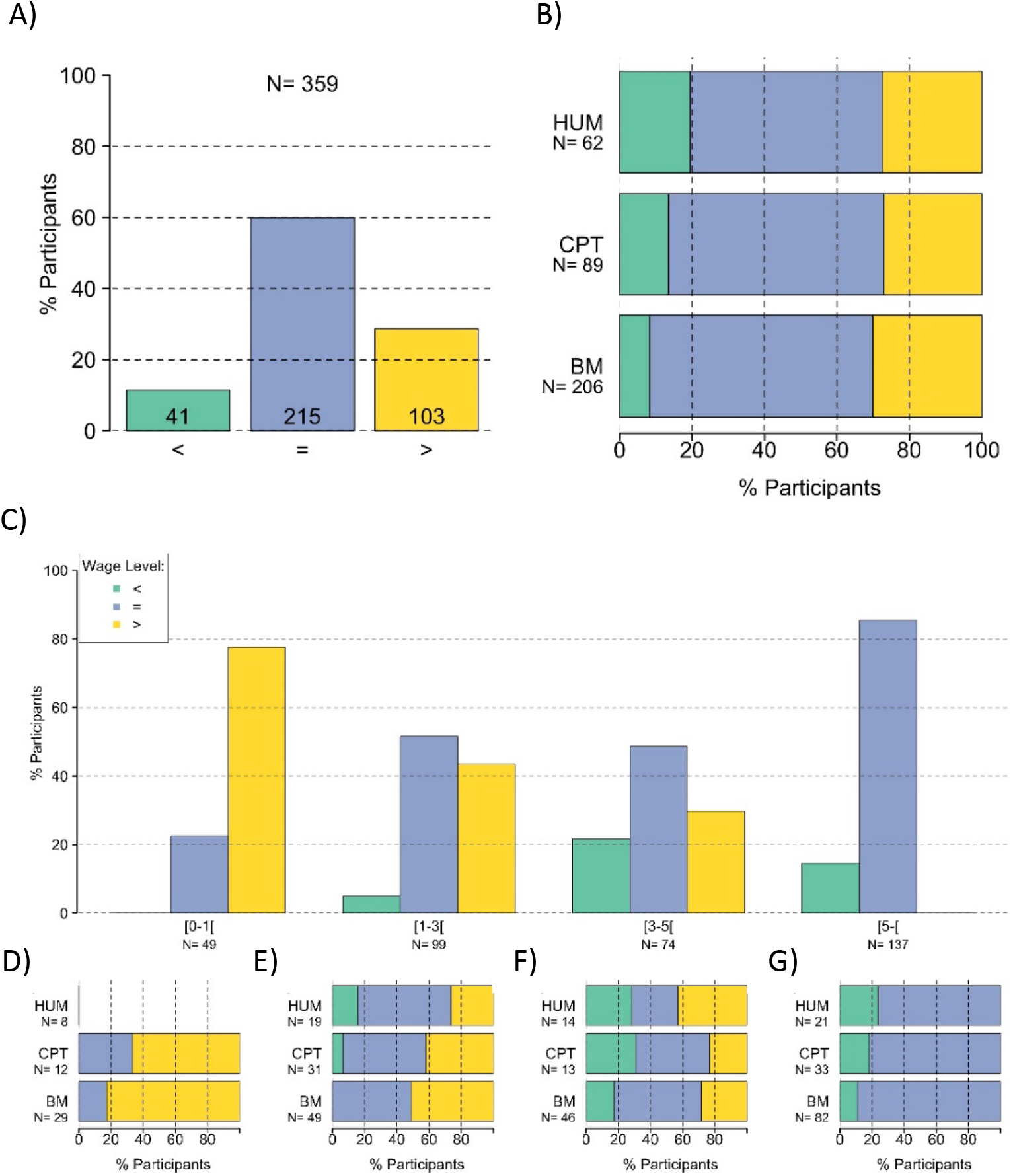
Appropriateness of wage level (*Stufe*) assignment. (**A & B**) Fraction of postdocs placed in the appropriate wage level in the MPS **(A)** and in each MPS section **(B)**. (**C** – **G**) **Appropriateness of wage level (*Stufe*) assignment and overall experience.** Fraction of postdocs placed in the appropriate wage level in the MPS **(C)** and in each MPS section depending on the overall postdoctoral experience (in year): less than one year of experience **(D)**, less than 3 years **(E)**, less than 5 years **(F)** and more than 5 years of overall experience **(F)**. Overall experience represents the number of years worked after PhD graduation. Postdocs found either in lower (green, <), appropriate (blue, =) or higher (yellow, >) wage level. BM, Biology and Medical section; CPT, Chemistry, Physics and Technology section; HUM, Human Sciences section.

**Figure S10.**
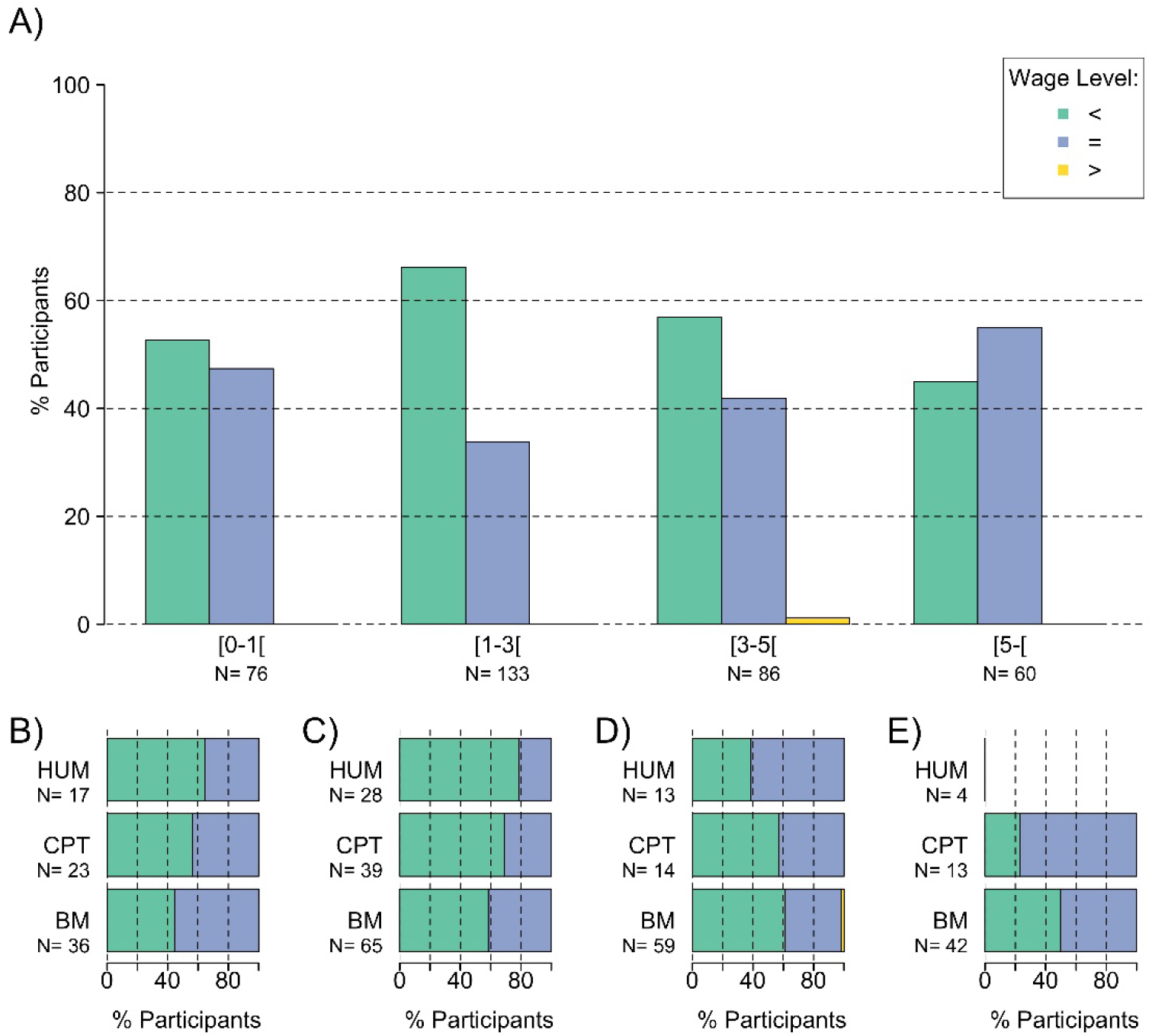
Appropriateness of wage level (*Stufe*) assignment and MPS experience. Fraction of postdocs placed in the appropriate wage level after recognition of a three-year PhD experience in the MPS **(A)** and in each MPS section depending on the MPS experiences (in year): less than one year of experience **(B)**, less than 3 years **(C)**, less than 5 years **(D)** and more than 5 years of MPS experience **(E)**. MPS experience represents the number of years worked within the MPS after PhD graduation. Postdocs found either in lower (green, <), appropriate (blue, =) or higher (yellow, >) wage level. This analysis confirms the general pattern that doctoral experience was not recognized for a large fraction of postdocs. Note, statistics with less than 10 data points are not shown on graphs. BM, Biology and Medical section; CPT, Chemistry, Physics and Technology section; HUM, Human Sciences section.

## Notes

### Competing Interest Statement

The authors have declared no competing interest.

