## Supplemental File 1 for "Demographics and Employment of Max-Planck Society’s Postdocs"

### Fair pay and employment for Postdocs in the Max Planck Society

In order to prepare the meeting of the PostdocNet steering group with various members of the Max Planck Society (MPS) headquarters in Munich mid-September, we would like to have some data about the PostdocNet community and the employment types and salary categories in the different institutes of the MPS.

The Postdoctoral working group in charge of the conduction of this survey would like to emphases two points:

1. We decide to make all questions optional, to let you define if giving certain answers may jeopardize your anonymity. However, we would like to stress that we are working carefully to ensure anonymity and that no-one can be single out.
2. The more answers we can obtain thanks to your participation, the better our evaluation of Postdoc needs will be. Therefore, our statements in front of MPS headquarters and the further improvement for our community will be boosted!

There are 32 questions in this survey.

### I. General information - demographics

In the following questions, the abbreviation "MPS" refers to any Institute, Research unit, Research group, Associated institute that make up the Max Planck Society (https://www.mpg.de/institutes_map).

##### 1. What is your gender?

*Choose one of the following answers*

- Female
- Male
- Other: ________________________

##### 2. How old are you?

*Choose one of the following answers*

- 20-30
- 30-40
- 40+

##### 3. Where are you from?

Should you have multiple nationalities/origins, please select the one you feel best represents you.

*Choose one of the following answers*

- Germany
- Europe - within EU
- Europe - outside EU
- Asia
- North America
- South America
- Africa
- Oceania

##### 4. Where did you obtain your PhD?

*Choose one of the following answers*

- At the MPS
- At another German institution
- At another European institution
- At a non-European institution

##### 5. How many years of research experience do you have, excluding your PhD years?

*Choose one of the following answers*

- Less than 1 year
- Between 1 and 3 years
- Between 3 and 5 years
- More than 5 years

##### 6. How long have you been working at the MPS as a Postdoc?

If you were employed several times at the MPS, please count your overall stay and do NOT count years spent as a PhD.

*Choose one of the following answers*

- Less than 1 year
- Between 1 and 3 years
- Between 3 and 5 years

- More than 5 years

##### 7. How many years of Postdoc experience did you have when you started working at the MPS?

If you were employed in another Max Planck Institute (MPI), please answer based on your current position.
Please write your answer here: ________________________

##### 8. Of those Postdoc years, how many were:

At another MPI: ________________________
At another German institution: ________________________
At another European institution: ________________________
At another institution: ________________________

##### 9. To which MPS section do you belong?

*Choose one of the following answers*

- Biology and Medicine Section (BMS)
- Humanities Section (HS)
- Chemistry, Physics and Technology Section (CPTS)

##### 10. To which particular MPI do you belong?

MPI are listed below in alphabetical order. If the name of your MPI is not in the list, select "Other" and specify.
*Choose one of the following answers*

- Associated Institute - Ernst Strüngmann Institute (ESI) for Neuroscience
- Associated Institute - Research Center caesar (center of advanced European studies and research)
- Bibliotheca Hertziana - MPI for Art History
- Friedrich Miescher Laboratory of the Max Planck Society
- Fritz Haber Institute of the Max Planck Society
- Kunsthistorisches Institut in Florenz – MPI
- Max Planck Florida Institute for Neuroscience
- MPI for Astronomy
- MPI for Astrophysics
- MPI for Biogeochemistry
- MPI for Biological Cybernetics
- MPI for Biology of Ageing
- MPI for Biophysical Chemistry
- MPI for Brain Research
- MPI for Chemical Ecology
- MPI for Chemical Energy Conversion
- MPI for Chemical Physics of Solids
- MPI for Chemistry
- MPI for Coal Research
- MPI for Comparative and International Private Law
- MPI for Comparative Public Law and International Law
- MPI for Demographic Research
- MPI for Developmental Biology
- MPI for Dynamics and Self-Organization
- MPI for Dynamics of Complex Technical Systems
- MPI for Empirical Aesthetics
- MPI for European Legal History
- MPI for Evolutionary Anthropology
- MPI for Evolutionary Biology

- MPI for Experimental Medicine
- MPI for Extraterrestrial Physics
- MPI for Foreign and International Criminal Law
- MPI for Gravitational Physics
- MPI for Gravitational Physics (Hannover)
- MPI for Heart and Lung Research
- MPI for Human Cognitive and Brain Sciences
- MPI for Human Development
- MPI for Infection Biology
- MPI for Informatics
- MPI for Innovation and Competition
- MPI for Intelligent Systems, Stuttgart site
- MPI for Intelligent Systems, Tübingen site
- MPI for Iron Research
- MPI for Marine Microbiology
- MPI for Mathematics
- MPI for Mathematics in the Sciences
- MPI for Medical Research
- MPI for Metabolism Research
- MPI for Meteorology
- MPI for Molecular Biomedicine
- MPI for Molecular Genetics
- MPI for Nuclear Physics
- MPI for Ornithology
- MPI for Physics
- MPI for Plant Breeding Research
- MPI for Plasma Physics
- MPI for Plasma Physics (Greifswald)
- MPI for Polymer Research
- MPI for Psycholinguistics
- MPI for Radio Astronomy
- MPI for Research on Collective Goods
- MPI for Social Anthropology
- MPI for Social Law and Social Policy
- MPI for Software Systems, Kaiserslautern site
- MPI for Software Systems, Saarbrücken site
- MPI for Solar System Research
- MPI for Solid State Research
- MPI for Tax Law and Public Finance
- MPI for Terrestrial Microbiology
- MPI for the History of Science
- MPI for the Physics of Complex Systems
- MPI for the Science of Human History
- MPI for the Science of Light
- MPI for the Structure and Dynamics of Matter
- MPI for the Study of Religious and Ethnic Diversity
- MPI for the Study of Societies
- MPI Luxembourg for International, European and Regulatory Procedural Law
- MPI of Animal Behavior
- MPI of Biochemistry
- MPI of Biophysics
- MPI of Colloids and Interfaces
- MPI of Immunobiology and Epigenetics

- MPI of Microstructure Physics
- MPI of Molecular Cell Biology and Genetics
- MPI of Molecular Physiology
- MPI of Molecular Plant Physiology
- MPI of Neurobiology
- MPI of Psychiatry
- MPI of Quantum Optics
- Max Planck Research Unit for Neurogenetics
- Max Planck Research Unit for the Science of Pathogens
- Research Group Social Neuroscience
- Other: ________________________

### II. Employment and salary at the MPS

Note: the below definitions were established by the PostdocNet and do not engaged in any case the responsibility of the MPS.

A **stipend** is a payment made for living expenses (e.g. fellowship, grant). As it is not considered wages, you are not paying for Social Security or Medical Taxes on it and your employer will not withhold any income taxes from the stipend. However, it can still be counted as taxable income for income tax purposes in certain cases.
An **employment contract** is a salaried position governed by the Collective Wage Agreement for the Civil Service (e.g. TVöD13). You are paying for Social Security or Medical Taxes on it and your employer will withhold some income taxes from the contract.

##### 11. Currently, how are you employed at the MPS?

Please, select the option that is covering your main income.
*Choose one of the following answers*

- Stipend = no social benefits included (health insurance, unemployment money, retirement money), no taxes. You are paying your health insurance on your own. Please check this box if you are on stipend, regardless the origin of it (MPS or funding agency).
- Fixed-term employment contract = you pay taxes and social charges (health insurance, unemployment money, retirement money…) on your salary (brutto vs netto), the institute pays half of your health insurance. Your contract is time-limited.
- Permanent employment contract = same as fixed-term but without end.
- Other: ________________________

##### 12. What is the origin of the funding sources that pay you?

*Choose one of the following answers*

- Max Planck Society
- German Research Foundation (e.g. DFG)
- European Research Foundation (e.g. ERC)
- Industry
- Private Foundations
- I do not know
- Other: ________________________

### III. You are a stipend holder

In the case that you were several times on a stipend at the MPS, please consider only your current position.

##### 13. What is the monthly allowance of your stipend in euros?

________________________

##### 14. Where is your stipend coming from?

*Choose one of the following answers*

- From a grant you personally obtained from a funding agency (e.g. AvH or EMBO fellowship)
- From the MPS or a grant that your superior obtained

##### 15. Please, specify the exact origin of your stipend.

________________________

##### 16. When did your stipend start?

*Choose one of the following answers*

- Before 2015
- After 2015

##### 17. What is the initial duration of your stipend?

*Choose one of the following answers*

- 6 months
- Between 6 months and 1 year
- Between 1 year and 1.5 year
- Between 1.5 year and 2 years
- Over 2 years

##### 18. Does the MPS add in any way to your actual stipend (e.g. mini contract)?

- Yes
- No
- No answer

##### 19. Do you have (either supported by you or your current MPS):

| | Yes | Uncertain | No | No answer |
|---|---|---|---|---|
| • A general health insurance? | | | | |
| • Right to a parental leave? | | | | |
| • An unemployment insurance? | | | | |
| • A pension contribution? | | | | |

##### 20. Do you pay taxes?

*Choose one of the following answers*

- Yes
- Yes, I should have but I obtained an exception from the Taxes declaration office (Finanzamt)
- No

##### 21. Do you have the same rights as contract holders in your institute (e.g. electoral rights for institute works council and Ombudsperson, equipment usage right, Childcare support, etc.)?

*Choose one of the following answers*

- Yes
- Partially (specify if you want, in the general comments)
- No
- I do not know

##### 22. Were you aware that you would be employed on a stipend before you started your position?

Please, consider the first time where you received a stipend at the MPS.

*Choose one of the following answers*

- Yes
- No

##### 23. Were you aware of the difference between stipend and contract before you started your position?

Please, consider the first time where you received a stipend at the MPS.

*Choose one of the following answers*

- Yes
- No

##### 24. If you wish to add a comment, fell free to use this section.

________________________

#### IV. You are a contract holder

To answer the following questions, you might need your most recent income statement. The information about your salary category (Tarifgr./-stufe) should be in a small box on the top right of your income statement along with the legal work load hours (Wöch AZ); your brutto salary should appear at the beginning of the detailed accounts.

##### 25. Are you paid in the E13 category of the TVöD?

*Choose one of the following answers*

- Yes
- No, please specify your category: ______________________

##### 26. Which 'Stufe' (subcategory) do you belong to?

______________________

##### 27. What is the work load in hours (Wöch AZ) stated on your income statement?

______________________

##### 28. What is the monthly brutto salary stated on your income statement?

______________________

##### 29. Were your previous work experiences considered to decide your salary category?

*Choose one of the following answers*

- Yes
- Partially
- No
- I do not know

##### 30. Were you informed about the category you would belong to before you started?

*Choose one of the following answers*

- Yes
- No
- I do not remember

##### 31. Were you informed why you would belong to that category before you started?

*Choose one of the following answers*

- Yes
- No
- I do not remember

##### 32. If you wish to add a comment, fell free to use this section.

______________________
